# Mechanism of bacterial predation via ixotrophy

**DOI:** 10.1101/2024.01.29.577165

**Authors:** Yun-Wei Lien, Davide Amendola, Kang Soo Lee, Nina Bartlau, Jingwei Xu, Go Furusawa, Martin F. Polz, Roman Stocker, Gregor L. Weiss, Martin Pilhofer

## Abstract

Predation allows bacteria to access alternative substrates in low-nutrient conditions. Ixotrophy has been proposed as a predatory lifestyle of multicellular filamentous bacteria in aquatic environments; however, the molecular mechanism remains unknown.

Here we uncover by a multidisciplinary approach that ixotrophy requires the interplay of multiple cellular machineries and a regulatory mechanism. Attacker-prey contacts are established by gliding motility and extracellular grappling hook-like structures that bind prey flagella. Cryo-electron microscopy identifies the grappling hooks as a heptameric assembly of a Type 9 Secretion System substrate. Cryo-electron tomography and functional assays show that killing is mediated by puncturing of the prey cell using a Type 6 Secretion System, possibly triggered by extracellular antennae. Single-cell analyses with stable isotope-labeled prey demonstrate that prey components are taken up by the attacker. Depending on nutrient availability, ixotrophy is switched off by endogenous Insertion Sequence Elements and re-activated through their excision. A marine metagenomic time series provides evidence for coupled dynamics of ixotrophic bacteria and their prey.

Our study reveals the complex mechanism of a conserved microbial predatory lifestyle and indicates the need for its regulation in conditions where the expression of costly pathways is dispensable.

## Introduction

Freshwater and ocean habitats are often heterogeneous environments with nutrient concentrations varying drastically across space and time (Fenchel, 2002; Stocker, 2012). Predation is a strategy for bacteria to access substrate that is bound in living microbes. The number of known bacterial predatory mechanisms is surprisingly low – distinct examples were observed in *Myxococcus xanthus*, *Bdellovibrio*-like organisms, *Candidatus* Uab amorphum, and *Vampirococcus* species (Thiery & Kaimer, 2020; Laloux, 2020; Bratanis *et al*, 2020; Shiratori *et al*, 2019; Moreira *et al*, 2021).

Ixotrophy has been proposed as a widespread predatory strategy for strains of the phylum *Bacteroidota* (Lewin, 1997; Furusawa *et al*, 2015a). Ixotrophy is a facultative predatory lifestyle and the supplementation with the prey cells can be sufficient for the ixotrophic bacteria to grow. Known ixotrophic bacteria are filamentous and multicellular, and – just like flypaper – they catch prey cells and stick them to their cell surface, followed by the lysis of the prey. This antagonism towards the prey seems to be specific, with possible prey cells including bacterial species from *Vibrio*, *Photobacterium*, diverse cyanobacterial strains (e.g., *Anabaena*, *Synechosystis*, *Synechococcus*, *Microcystis*), and even eukaryotic organisms such as diatoms (e.g., *Chaetoceros ceratosporum*) (Lewin, 1997; Furusawa *et al*, 2015a; Shi *et al*, 2006; Ashton & Robarts, 1987; Furusawa *et al*, 2003; Svercel *et al*, 2011). While prey flagella have been reported to play a crucial role in ixotrophy, the underlying mechanisms of specificity, catching, and killing are not understood.

Some ixotrophy-positive strains contain intriguing rod-shaped particles, called “rhapidosomes” (Correll & Lewin, 1964; Lewin & Kiethe, 1965), which were speculated to be involved in gliding motility. Later, it was shown that these structures comprise a protein with similarities to a contractile injection system (CIS) from *Serratia entomophila* (Furusawa *et al*, 2005). Interestingly, CISs from other microorganisms have been shown to mediate diverse cell-cell interactions (Basler *et al*, 2012; Shikuma *et al*, 2014; Desfosses *et al*, 2019; Böck *et al*, 2017; Weiss *et al*, 2022; Xu *et al*, 2022). CISs are macromolecular injection devices with homologies to the contractile tails of phages, comprising a contractile sheath-tube module that is assembled on a baseplate structure. Upon firing, contraction of the sheath expels the inner tube and propels it into a target cell. Based on their mode of action, CISs are classified into two major groups: extracellular CISs (eCISs) and Type 6 Secretion Systems (T6SSs). eCIS are released into the medium and bind to a target cell, followed by firing and translocation of cargo (Desfosses *et al*, 2019; Jiang *et al*, 2019; Shikuma *et al*, 2014; Xu *et al*, 2022). T6SSs are anchored to the inner membrane via their baseplate and accessory components, and they can fire into neighboring cells in a cell-cell contact-dependent fashion (Brunet *et al*, 2015; Rapisarda *et al*, 2019; Basler *et al*, 2012). While “rhapidosomes” have similarities to CISs, their role and mode of action in ixotrophy are unclear.

Here we set out to explore the molecular mechanism of ixotrophy to answer the following questions: (i) How is the prey caught? (ii) How is the prey killed? (iii) How is ixotrophy regulated? (iv) Why do cells employ ixotrophy?

## Results

### Ixotrophy killing is efficient, contact-dependent and functions on solid and in liquid media

We studied ixotrophy in the strain *Aureispira* sp. CCB-QB1 (Furusawa *et al*, 2015b) (hereafter referred to as “*Aureispira*” or attacker) and established it as a model organism. DAPI staining depicted that filaments typically comprised ∼5-10 individual cells (**Fig. S1a**). Previous studies using co-culturing assays showed an ixotrophic behavior of *Aureispira* against *Vibrio* species (Furusawa *et al*, 2015a; Yeoh *et al*, 2021). To understand the dynamics at a single-cell level, we performed time-lapse light microscopy (LM) imaging of *Aureispira* cells co-incubated with *Vibrio campbellii* (hereafter “*Vc*”) as prey on an agarose pad. *Aureispira* filaments were frequently seen to either bulge out to approach a *Vc* cell (**Fig. 1a, Movie S1**) or to glide “head-on” toward a *Vc* cell (**Fig. S1b, Movie S2**). In ∼78% of such cell-cell contacts (*n* = 130), the *Vc* cells subsequently showed signs of cell lysis (**Fig. 1b**). Among them, many *Vc* cells (∼60%) immediately lysed within seconds after the cell-cell contact, while in ∼18% of the cell-cell contacts the *Vc* cells showed signs of rounding and often lysed at a later stage (**Fig. 1b**). Such antagonistic effects were not seen in the absence of cell-cell contact (*n* = 102). The need for contact was further supported by a bulk killing assay on a solid medium, in which killing of *Vc* was only observed by co-incubation of *Vc* with live *Aureispira* cells, but not with culture supernatant or heat-inactivated attacker cells (**Fig. 1c/S1c**). Interestingly, even though killing appears to be contact-dependent, ixotrophy also functions in a liquid medium. We hypothesized that *Aureispira* must be able to make contact with prey cells efficiently. In fact, consistent with previous reports (Lewin, 1997; Furusawa *et al*, 2015a), *Aureispira* was decorated by multiple *Vc* cells within only a few seconds from the onset of co-incubation (**Fig. 1d**). Interestingly, the specificity towards a certain prey occurs only in liquid medium and a wider range of prey was killed indiscriminately on solid agar surface (**Fig. S1d**). We conclude that ixotrophy-mediated killing functions in liquid and on solid media and requires direct cell-cell contact.

**Figure 1.**
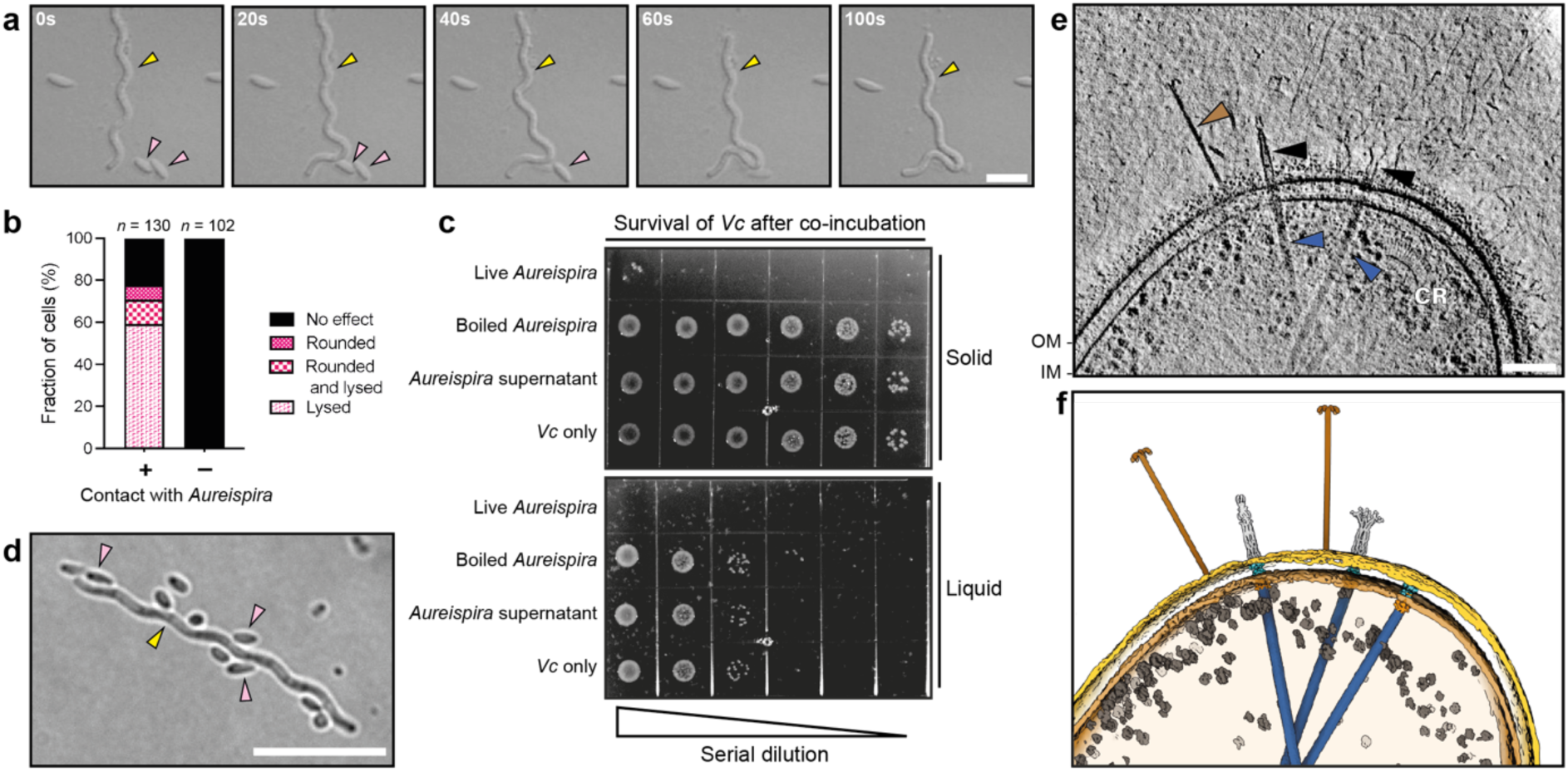
*Aureispira* shows contact-dependent killing in liquid and on solid media, and expresses an arsenal of supramolecular machines. **a:** Time-lapse light microscopy (LM) images show an *Aureispira* filament (yellow arrowhead) approaching *Vc* prey cells (pink arrowheads) followed by rapid lysis of the prey. Bar: 5 μm. **b:** Quantification of the fate of *Vc* prey cells in time-lapse LM movies revealed that ∼80 % of *Vc* cells showed signs of cell lysis after cell-cell contact with *Aureispira*. No signs of cell lysis were observed for *Vc* cells without direct contact with *Aureispira* filaments. Sample size *n* is indicated at the top of the chart. **c:** Serial dilution of prey cells after co-incubation killing assays suggests that ixotrophy killing is contact-dependent. *Vc* was co-incubated either in liquid culture (top panel, 6 h of co-incubation) or on a solid agar plate (bottom panel, 24 h of co-incubation) with differently treated *Aureispira* cells (indicated on the left). The cultures were serially diluted and dropped onto an agar plate to visualize the survival of *Vc*. See **Fig. S1b** for the quantification of *Vc* before the start of the co-incubation assay. **d:** LM image shows an *Aureispira* filament (yellow arrowhead) decorated with *Vc* cells (pink arrowheads) after co-incubation in liquid culture. Bar: 10 μm. **e:** Slice through a cryo-tomogram of *Aureispira* reveals several supramolecular machineries. Attached to the OM were grappling hook-like structures and multiple IM-bound T6SS-like apparatuses. T6SSs were often associated with extracellular antenna-like densities that were seen in two distinct conformations. CR: chemoreceptor arrays. Bar: 100 nm. Thickness of the slice: 13.8 nm. **f:** Shown is a segmentation of the tomogram in (**d**). OM: yellow, IM: light brown, grappling hooks: brown, T6SS sheaths: blue, T6SS baseplates: orange, T6SS trans-envelope complexes: turquoise, antennae: light gray and ribosomes: dark gray.

### *Aureispira* cells express elaborate cell envelope-associated supramolecular machines

In order to observe ultrastructures that may be involved in catching and killing of the prey, we plunge-froze *Aureispira* cells on electron microscopy (EM) grids and imaged them by cryo-electron tomography (cryoET). The cells had a typical Gram-negative cell envelope that was associated with distinct supramolecular machines (**Fig. 1e/f, Movie S3**). Attached to the outer membrane (OM) were numerous rigid “grappling hook”-like structures. Furthermore, all cells exhibited multiple T6SS-like apparatuses in mostly extended and rarely contracted conformations. Interestingly, T6SSs were often associated with extracellular “antenna”-like structures. Some cells also showed putative chemoreceptor arrays (CR) in the cytoplasm.

### Grappling hooks interact with prey flagella and mediate cell-cell contacts in liquid medium

To test whether the grappling hooks made contact with *Vc* cells, we imaged a mixture of attacker and prey cells. Due to the thickness of the sample, the frozen cells had to be thinned by cryo-focused ion beam (FIB) milling prior to cryoET imaging (Marko *et al*, 2007; Rigort & Plitzko, 2015; Zachs *et al*, 2020). Our cryo-tomograms revealed that grappling hooks interacted with the flagella of the prey cells (**Fig. 2a**). To test the relevance of this interaction for ixotrophy, we co-incubated *Aureispira* with *Vc* wild-type (WT) and *Vc* flagellin deletion mutants (lacking flagella; **Fig S2a**), respectively. LM imaging immediately after mixing showed that *Aureispira* was heavily decorated by *Vc* WT cells (on average 5.8 cells per *Aureispira* filament, *n* = 320), whereas *Vc* Δ*flagellin* showed a significant reduction of decoration of *Aureispira* (on average 0.9 cell per *Aureispira* filament, *n* = 290) (**Figs. 2b, S2b**). Finally, a killing assay with either *Vc* WT or *Vc* Δ*flagellin* indicated slower killing rates for the mutant in liquid (**Fig. 2c, S2c**) but no difference when co-incubated on solid agar (**Fig. S2d**). These results suggest that, in liquid culture, cell-cell contacts between *Aureispira* and *Vc* are facilitated via the interaction of *Aureispira* grappling hooks with *Vc* flagella.

**Figure 2.**
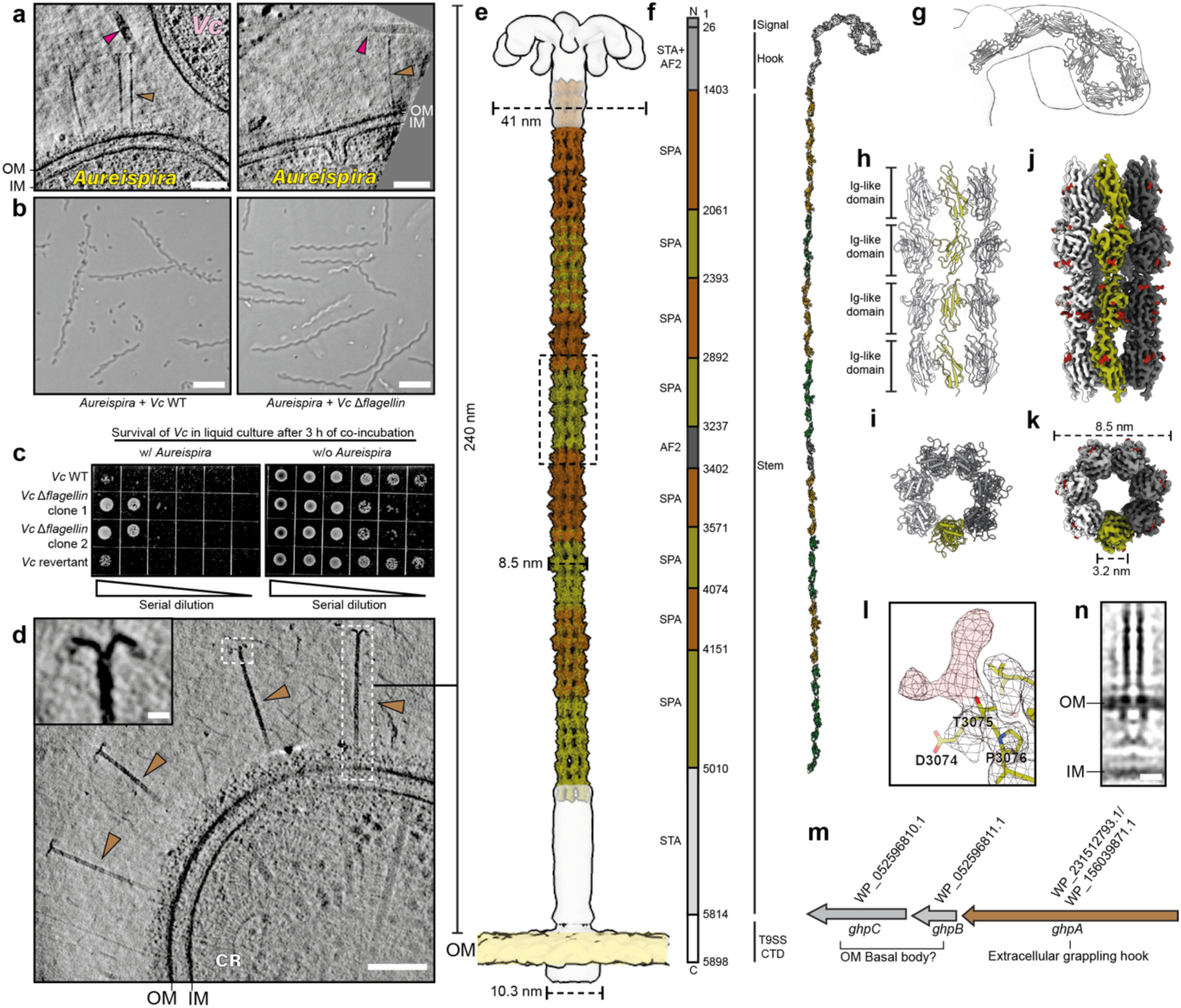
Heptameric grappling hooks interact with prey flagella. **a:** CryoET of cryoFIB-thinned *Aureispira*–*Vc* prey mixture reveals interactions between *Aureispira* grappling hooks (brown arrowhead) and prey flagella (magenta arrowhead). Shown are 13.8 nm thick slices. Bar: 100 nm. **b:** LM imaging of *Aureispira* co-incubated in liquid culture with either *Vc* WT (left) or *Vc* flagella-mutant (Δ*flagellin*, right). Prey without flagella were not caught by *Aureispira*. Bar: 10 μm. **c:** Serial dilution drop assay of killing assays in liquid showed that the *Vc* Δ*flagellin* mutants were killed slower by *Aureispira* compared to flagellated *Vc* WT. Prey cells with different genotypes were co-incubated with or without *Aureispira* for 3 h, respectively. See **Fig. S2c** for assays showing the survival of the prey at different time points during co-incubation. **d:** A slice through a cryo-electron tomogram showing an *Aureispira* cell with several homogenous extracellular grappling hooks (brown arrowhead). Bar: 100 nm. Shown is 13.8 nm thick slice. Magnified view of the distal end of a grappling hook is provided in the inset. Scale bar: 10 nm. **e:** A composite density map combining results of subtomogram averaging and single particle cryoEM to show the overall structure of a grappling hook. Density maps obtained by subtomogram averaging are shown in light gray, and density maps obtained from single particle cryoEM are shown in brown or green. OM: yellow. **f:** Shown is a GhpA schematic representation (left), indication of structural modules (middle) and composite atomic model (right). Methodologies utilized to determine the structure of GhpA fragments and their corresponding amino acid residues are indicated next to the schematic (AF2: Alphafold2 prediction. STA: subtomogram averaging. SPA: single particle cryoEM). **g:** Atomic model of the distal hook was predicted by Alphafold2 (residues 26-1,402, gray) and fitted into the subtomogram averaging density map. **h/i:** Shown is the atomic model (side view in **h**, top view in **i**) of the stem region of GhpA (residue 2,892-3,236, corresponding to the dashed box in **e**), revealing seven parallel strands of GhpA with a hollow lumen and individual strands composed of successive Ig-like domains. See **Fig. S9** for atomic models of other fragments. Individual GhpA strands are shown in different shades of gray/green. **j/k:** Surface rendering of the SPA density map of the fragment of the grappling hook stem region (side view in **j**, top view in **k**; corresponding to dashed box in **e**). Bulky densities, which could not be assigned to an amino acid side chain are shown in red and likely correspond to glycosylation sites. See **Fig. S9** for surface renderings of SPA density maps of other fragments. Color code as in **h/i**. **l:** An example showing the SPA density map of a potential branched glycan on the grappling hook stem at Thr-3,075. The potential glycan is highlighted in light red. **m:** Schematic of the grappling hook gene cluster. Next to GhpA, it encodes putative OM basal body proteins GhpB (homolog of PorP) and GhpC (homolog of PorE). **n:** Longitudinal slice through C7 rotationally symmetrized subtomogram average of the OM basal body of grappling hooks. Bar: 10 nm.

### Grappling hooks are formed by a heptamer of a Type 9 Secretion System substrate

We set out to identify and structurally characterize the grappling hook. Multiple grappling hooks with a uniform length (240 ± 4 nm; mean ± s.d.; *n* = 55) were distributed along the cell body in every cryo-tomogram (*n* = 146) (**Fig. 2d**). Sub-tomogram averaging of the distal end revealed a terminal structure consisting of seven hook-like densities (**Fig. 2e, S3**). For protein identification, grappling hooks were sheared off, further purified, and subjected to single particle cryo-electron microscopy (cryoEM) (**Fig. S4-S6**). We determined eight distinct density maps at a global resolution between 3.4 – 3.8 Å that correspond to different regions along the grappling hook stem module (shown in different colors in **Fig. 2e**; local resolution of density maps is shown in **Fig. S7**). Consistent with the presence of seven terminal hooks, the structures of all stem regions show a C7-symmetry with a hollow lumen. For identification of involved proteins, the program ModelAngelo (Jamali *et al*, 2023) was used to predict the primary amino acid sequence based on the density maps. These sequences were then blasted against the *Aureispira* genome. The eight density maps all correspond to the same protein (WP_231512793.1/WP_156039871 [spread across two contigs]), which we will refer to as Grappling Hook Protein A (GhpA). Seven protomers of GhpA align longitudinally to form the grappling hook emanating from the OM.

GhpA is a 5,898 amino acid residue-long protein with a predicted molecular weight of 610 kDa (**Fig. 2f**). Sequence and structure homology analyses revealed a N-terminal Sec secretion system signal peptide (residue 1 to 25 residues), followed by a bacterial adhesin-like domain that corresponds to the distal hook module. The hook module could not be resolved by single particle cryoEM due to its flexibility. An Alphafold2-prediction of the N-terminal region (residues 26 to 1402), however, resulted in a loop-like structure, which fits with high confidence into the sub-tomogram average (**Fig. 2g**).

The stem module is composed of seven longitudinally aligned GhpA strands, which form a straight and hollow structure (**Fig. 2f/h/i**). The GhpA stem region comprises 52 slightly varying Ig-like repeats. Lateral interactions between the protomers are mediated by inter-molecular hydrogen bonds (**Fig. S8a/b**). Furthermore, a multitude of intra-molecular disulfide bonds and hydrogen bonds were found between nearby Ig-like domains (**Fig. S8**). Together, these interactions are likely the basis for the straight and rigid architecture of the stem. In addition, the surface of the stem module shows bulky densities nearby threonine and serine residues that likely correspond to O-glycosylation sites (**Fig. 2j/k, S9**). These potential glycans are found in all the Ig-like domains, and they are often branched (**Fig. 2l, S9**), consistent with the general glycosylation pattern found in other *Bacteroidota* species (Coyne *et al*, 2013; Veith *et al*, 2023). The proximal region of the stem was less well resolved, presumably due to the fact that grappling hooks were sheared off during purification, inducing structural flexibility in this region.

The GhpA C-terminus encodes a Type 9 Secretion System (T9SS) substrate sorting domain. In *Porphyromonas gingivalis*, a T9SS substrate is speculated to be anchored on the cell surface by interacting with PorP and PorE (Gorasia *et al*, 2022b), which encode OM beta-barrel and OmpA-like peptidoglycan binding domains, respectively. We also detected such homologs downstream from *ghpA* (here referred to as *ghpB/C*) (**Fig. 2m**). Sub-tomogram averaging of the anchoring region of grappling hooks revealed a structure located in the OM/periplasm (**Fig. 2n**), which may account for GhpB/C. Altogether, we conclude that grappling hooks are assembled from a heptameric T9SS substrate.

### *Aureispira* assembles Type 6 Secretion Systems (T6SSs) with unique features

Next, we set out to investigate the structure and the role of *Aureispira* T6SSs (**Fig. 1e/f**). We first identified a candidate T6SS gene cluster (**Fig. 3a**) in the *Aureispira* genome (Furusawa *et al*, 2015b). The cluster comprises genes for typical structural components, which we renamed *cis1*-*cis16* according to previously characterized CISs (Weiss *et al*, 2022; Xu *et al*, 2022; Desfosses *et al*, 2019) (**Table S1**). Proteomics of a crude purification confirmed that *Aureispira* T6SSs were encoded by these genes (**Fig. 3a, S10a, Table S2**). Close relatives of the *Aureispira* system are the eCIS in *Algoriphagus machipongonensis* (Xu *et al*, 2022), the eCIS in *Pseudoalteromonas luteoviolacaea* (Shikuma *et al*, 2014), the thylakoid-anchored CISs in cyanobacteria (Weiss *et al*, 2022), and the T6SS*^iv^* in *Ca.* A. asiaticus (Böck *et al*, 2017) (**Fig. S11**). Based on these phylogenetics and cryoET imaging, we classify the *Aureispira* T6SS as subtype *iv* (T6SS*^iv^*). The previously reported “rhapidosomes” (Lewin & Kiethe, 1965) are likely T6SS*^iv^* structures.

**Figure 3.**
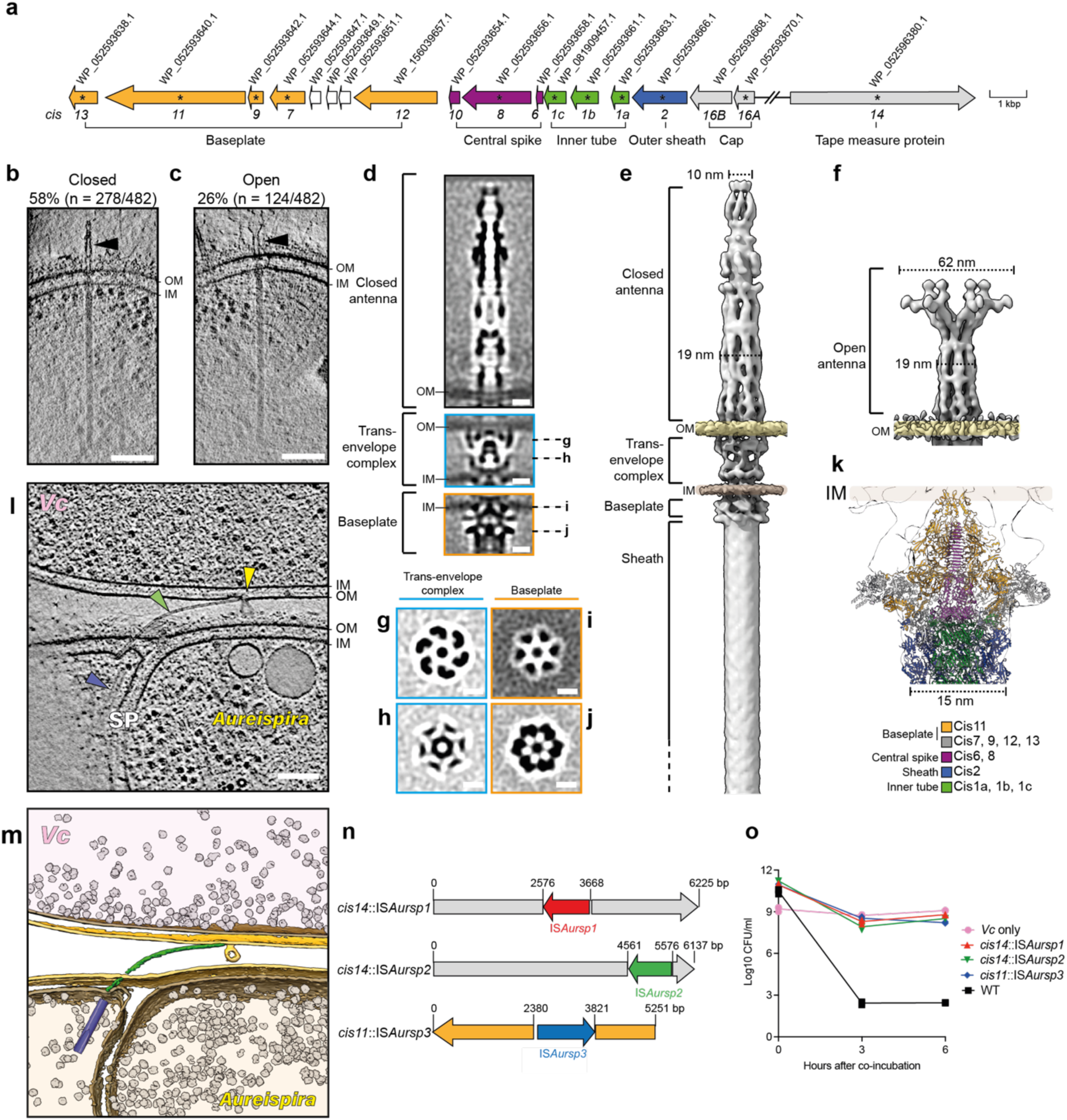
A type 6 secretion system with unique features kills prey. **a:** Schematic representation of the *Aureispira* T6SS*^iv^*gene cluster. Accession numbers are indicated above, and genes were renamed *cis1* – *cis16*. The assignment of genes to functional modules is indicated below. Asterisks indicate components that were detected in purification via mass spectrometry. **b/c:** Examples of cryo-tomograms of *Aureispira* T6SSs with extracellular antennae (black arrowheads) in either “closed” (**b**) or “open” conformations (**c**). Quantification of observed antenna conformation is indicated above. Bar: 100 nm. Thickness of the cryoET slices: 13.8 nm. **d:** Slices through subtomogram averages of extracellular antenna, trans-envelope complex and baseplate of *Aureispira* T6SSs, respectively. Bar: 10 nm. **e:** Composite isosurface representation of the subtomogram averages of closed antenna, trans-envelope complex, baseplate and sheath-tube modules. **f:** Isosurface representation of the subtomogram average of the open antenna. **g-j:** Perpendicular slices through the C6-symmetrized subtomogram average of the trans-envelope complex (**g/h**) and the baseplate (**i/j**). The positions of the slices are indicated in **d**. Bar: 10 nm. **k:** Docking of the atomic model of thylakoid-anchored CIS (PDB: 7B5H) into the density map of the subtomogram average of the *Aureispira* T6SS baseplate reveals conserved elements like a cage (Cis11, orange) surrounding the spike (Cis8, purple). **l:** Slice through a cryo-tomogram of a cryoFIB-thinned *Aureispira*-*Vc* mixture reveals a contracted T6SS sheath (blue arrowhead) and an associated expelled inner tube (green arrowhead). The inner tube is seen penetrating the OM of *Vc* (yellow arrowhead), apparently resulting in membrane vesiculation. SP: septum. Thickness of the slice: 13.8 nm. Bar: 100 nm. **m:** Shown is a segmentation of the tomogram shown in **l.** OM: yellow. IM: light brown, contracted T6SS sheath: blue, T6SS inner tube: green, ribosomes: gray. **n:** Shown is a schematic illustrating the interruption of T6SS genes by insertion sequences (ISs) in T6SS-negative *Aureispira* derivatives. **o:** Quantification of colony forming units (CFUs) of prey cells during killing assay (in liquid culture) with *Aureispira* WT or *Aureispira* T6SS-negative derivatives. CFUs of *Vc* were quantified at several time points by serial dilution of the culture and plating on agar, revealing that T6SS-negative *Aureispira* derivates were unable to kill *Vc*.

In cryo-tomograms, T6SSs were almost always seen anchored to the inner membrane (IM) (**Fig. 3b/c**). Extended T6SSs had a homogeneous length of 420 ± 8 nm (mean ± s.d.; *n* = 45), consistent with the presence of a putative tape measure protein (*cis14* in **Fig. 3a**). On average, we detected 4.6 extended T6SSs per tomogram (**Fig. S10b**). Interestingly, *Aureispira* T6SSs were often associated with extracellular antenna-like structures. These antennae showed two distinct conformations that we term “closed” (58 %; 278 out of 482) and “open” (26 %; 124 out of 482) (**Fig. 3b/c, S10c**).

We then performed subtomogram averaging of the individual T6SS modules, i.e. antenna, trans-envelope complex, baseplate and sheath. The antenna in the closed conformation revealed a complex C6-symmetric architecture, a length of 132 nm, and a width of 10-19 nm (**Fig. 3d-f/p, S12a-c, S13a/c-e/i**). The open conformation was much shorter (82 nm) and featured a similar architecture at the base (**Fig. S3f, S13b/f-h/i**). The distal part of the open conformation showed a significant widening (width of 62 nm) of the six arms that each split into two further protruding densities (**Fig. 3f, S13f/j**). To our knowledge, except for the much simpler extracellular structures connected to a T6SS in *Myxococcus* (Chang *et al*, 2017), such elaborate extracellular T6SS-associated structures have not been reported.

The C6-symmetric trans-envelope complex consisted of three main parts (**Fig. 3d/e/g/h, S12a/d/e, S14a-k**). Two components were associated with the OM and IM, respectively. A third periplasmic part complements the structure, resulting in a continuous OM-IM spanning complex. This trans-envelope complex is fundamentally different from the C5-symmetric TssJLM (no homologs in *Aureispira*) trans-envelope complex from the canonical T6SS*^i^* (Rapisarda *et al*, 2019). The structure also presents a major variation compared to the T6SS*^iv^* in *Ca.* A. asiaticus, which lacks a trans-envelope complex altogether (Böck *et al*, 2017).

The T6SS baseplate is also C6-symmetric, and its architecture is generally consistent with high-resolution structures of thylakoid-anchored CISs from cyanobacteria (Weiss *et al*, 2022) and eCISs from *Algoriphagus* (Xu *et al*, 2022) (**Fig. 3d/e/i/j, S12a/f/g, S14l-r**). All of these systems feature an extension of Cis11, which was shown to form a cage-like structure around the spike. This extension is conserved in the *Aureispira* Cis11, and a similar cage structure around the spike is seen in the baseplate average (**Fig. 3k**). The comparison with the T6SS*^iv^* in *Ca.* A. asiaticus (Böck *et al*, 2017) reveals that in both systems contact to the IM seems to be mediated by a set of six anchoring densities as well as by the cage (**Fig. 3d/e/i**).

### The *Aureispira* T6SS is required for the killing of prey

Next, we set out to understand whether the novel T6SS was involved in ixotrophy killing. Physical puncturing by the T6SS inner tube has long been hypothesized but never visualized (Pukatzki *et al*, 2007; Hernandez *et al*, 2020; Vettiger & Basler, 2016). Thus, we co-incubated *Aureispira* and *Vc* for 2-8 min and performed cryoFIB-milling and cryoET. Strikingly, we captured three intriguing attacker-prey contacts, showing a contracted sheath and its expelled inner tube protruding from the attacker and puncturing the OM of the prey (**Fig. 3l/m, Movie S4**). In one example, the OM showed signs of vesiculation and disruption at the puncturing site. The detection of three puncturing events in only 86 cryo-tomograms of ∼150-200 nm-thick lamellae indicates that these events are relatively long-lived or highly frequent.

Having established that the T6SS has a disruptive effect on the prey cell envelope, we further investigated whether it was also responsible for killing. Our attempts to establish a genetic system by site-directed mutagenesis were unsuccessful, given the multicellularity of the strains and the absence of genetic tools for close relatives. In the process, however, we identified several *Aureispira* derivatives (T6SS-negative) from which we could not purify T6SS sheaths and which did not show any assembled T6SSs in cryo-tomograms (**Fig. S15a/b, Table S2**).

Genome sequencing of the WT and T6SS-negative derivatives revealed that each mutant strain had an insertion of an “insertion sequence” (IS) into T6SS genes (**Fig. 3n, S14d-g**). All three ISs were different, and we therefore named them IS*Aursp1* (for IS *Aureispira* sp. 1), IS*Aursp2* and IS*Aursp3*, respectively. ISs are the simplest mobile genetic elements that encode only a transposase gene that can insert into various sites in the genome (Siguier *et al*, 2014). While IS*Aursp1* and IS*Aursp2* were inserted into different positions of gene *cis14* (tape measure protein, strains *cis14*::IS*Aursp1* and *cis14*::IS*Aursp2*), IS*Aursp3* was inserted into gene *cis11* (baseplate component, strain *cis11*::IS*Aursp3*). Since all three ISs are present elsewhere in the genomes of WT with an additional copy in each mutant strain, the origins of the T6SS-disrupting ISs were likely duplication events (**Table S3**).

Finally, we performed killing assays with WT and T6SS mutant strains and observed that the ability to kill was completely abolished in the T6SS-negative strains (**Fig. 3o**). Time-lapse imaging showed that the mutants were still able to glide and make repeated contact with prey cells, however, lysis of prey cells was never observed (**Movie S5**). In summary, *Aureispira* expresses abundant T6SSs with unique structural features that are required for killing and the system can be deactivated by endogenous ISs.

### Insertion sequences (ISs) switch T6SS activity dependent on nutrient availability

The spontaneous inactivation of the T6SS was observed in full medium, which posed the question of whether ISs may be a mechanism to down-regulate T6SS activity in nutrient-rich conditions. We, therefore, set out to investigate whether inactivation was reversible upon starvation. First, we determined the transcription levels of two genes involved specifically in IS excision. The IS-excision enhancer *iee* is an error-prone DNA polymerase previously found to enhance IS excision rate (Kusumoto *et al*, 2011; Calvo *et al*, 2023), whereas *recB*, a factor of the DNA double-strand break repair machinery, is required for successful IS excision (Sinha *et al*, 2020; Dorman, 2020). We compared transcription levels of *iee* and *recB* in cells that were cultured in full medium *versus* minimal medium and found that transcription of both genes was significantly upregulated during starvation (**Fig. 4a, S16a**).

**Figure 4.**
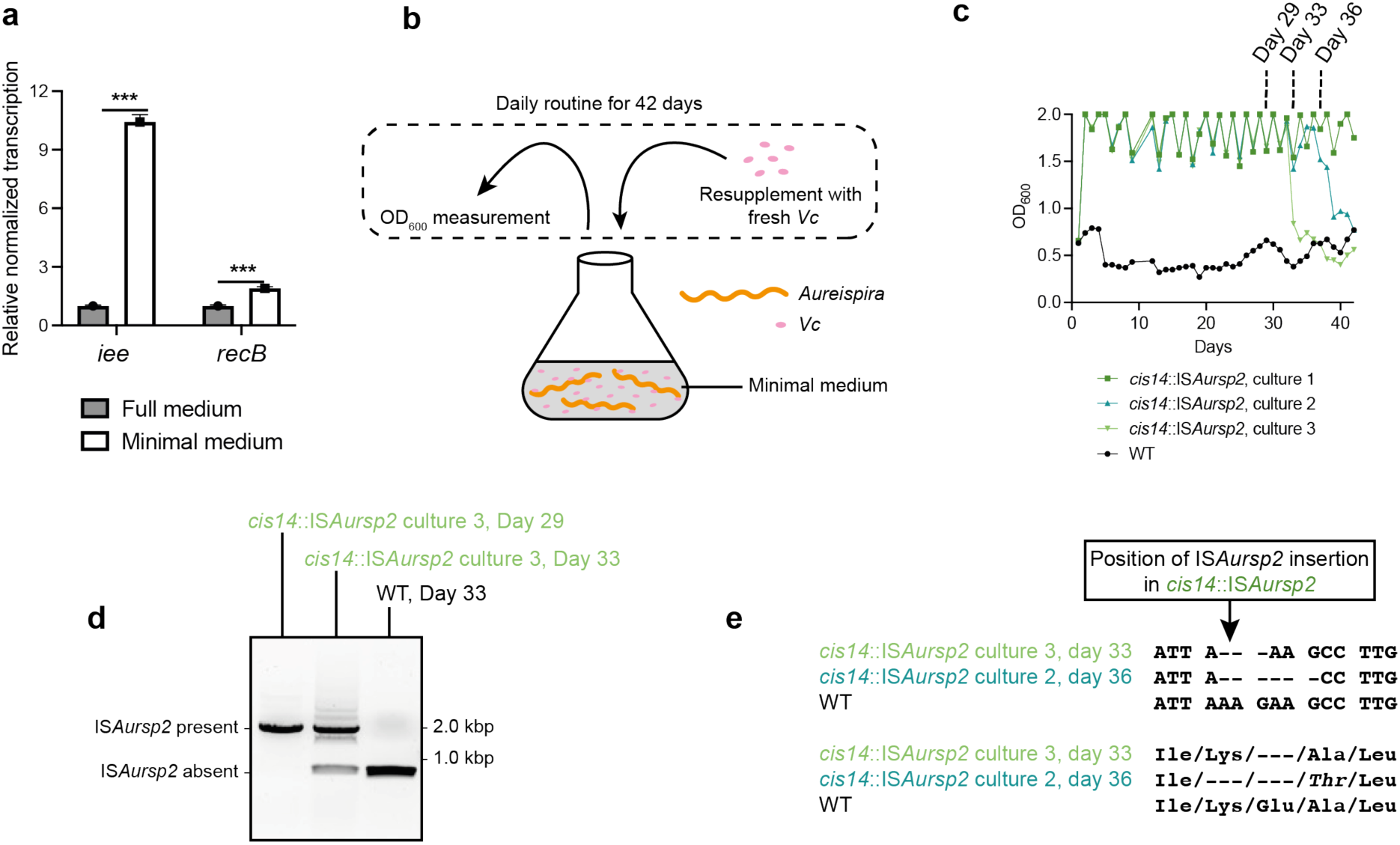
Insertion sequences control T6SS activity. **a:** Genes involved in IS excision (*iee*/*recB*) were upregulated during starvation. Total RNA from *Aureispira* cultured in full or minimal medium was extracted, and *iee/recB* transcripts were quantified. The transcription level was normalized using the housekeeping gene *gyrB*. **b:** Schematic showing the experimental setup that was used to analyze IS excision during starvation. OD_600_ values of the cultures were measured every day and afterwards re-adjusted to OD_600_ = 2 by adding prey cells. Experimental data is shown in (**c**). **c:** Shown are OD_600_ measurements of the *Aureispira-Vc* starvation experiment described in (**b)**. Until Day 32, only the WT culture showed killing activity (low OD_600_ values), while cultures 1-3 (T6SS-negative derivatives) did not kill (evidenced by high OD_600_ values). On Day 33 and 36, respectively, OD_600_ values of cultures 2 and 3 dropped, indicating re-activation of T6SS killing activity. **d:** IS*Aursp2* excision was confirmed by PCR amplification of a *cis14* fragment (primers shown in **Fig. S15d**). At Day 29, IS*Aursp2* was homogenously present in *cis14* of *Aureispira* culture 3. On Day 33, we detected a mixture of *cis14* fragments with inserted IS*Aursp2* as well as *cis14* WT fragments. **e:** PCR products from (**d)** and **Fig. S16b** were sequenced, revealing that IS excision generated 3 or 6 bp deletions in *cis14*. Sequencing results are shown in the top panel, and the corresponding translated amino acid sequence is indicated in the bottom panel.

We therefore hypothesized that under nutrient-limiting conditions, T6SS-negative strains may activate T6SS by IS excision. To test this, we cultured T6SS-positive (WT) and a T6SS-negative (*cis14*::IS*Aursp2*) strain under nutrient-limiting conditions but in the presence of *Vc* (**Fig. 4b**). The optical cell density (OD_600_) was measured daily and served as a general indicator for *Vc* survival. As expected, the T6SS-positive WT efficiently killed *Vc*, resulting in very low OD_600_ values (**Fig. 4c**). In contrast, in three independent cultures of the T6SS-negative mutant *cis14*::IS*Aursp2*, the OD_600_ values remained high over the course of ∼30 days, indicating the absence of killing activity. Strikingly, on days 33 and 36, respectively, the OD_600_ values in two of the mutant cultures suddenly dropped, indicating the re-activation of killing. Indeed, the excision of IS*Aursp2* in *cis14* was confirmed by PCR and sequencing (**Fig. 4d/e, S16b**). Interestingly, compared to the WT, both excision events resulted in the deletion of three and six base pairs, respectively (**Fig. 4e**). We conclude that IS elements constitute a previously unrecognized mechanism to deactivate T6SS expression in nutrient rich conditions by duplication events, and to activate T6SS expression in nutrient-limiting conditions by excision events.

### *Aureispira* takes up nutrients from killed prey cells

Next, we turned to the question of why *Aureispira* kills prey cells. To test whether *Aureispira* takes up nutrients from prey cells upon killing, *Vc* was labeled with deuterium (D). We first confirmed that D-labeled *Vc* can still be killed by *Aureispira* (**Fig. S17a**). The D-labeled prey cells were then co-incubated for 1 h with unlabeled *Aureispira* cells and dried on a sample support. We then measured the D-labeling status of the *Aureispira* cells using single-cell Raman microspectroscopy (**Fig. 5a**). For each *Aureispira* filament, we quantified the deuterium labeling status by normalizing the signal intensity of the carbon-deuterium (C-D) peak with that of the carbon-hydrogen (C-H) peak (a proxy for cell biomass) (Lee *et al*, 2021). The deuterium labeling status of T6SS-positive *Aureispira* was significantly higher compared to the T6SS-negative strain (**Fig. 5b, S17b/c**). These results suggest that *Aureispira* takes up and likely metabolizes nutrients from killed *Vc*.

**Figure 5.**
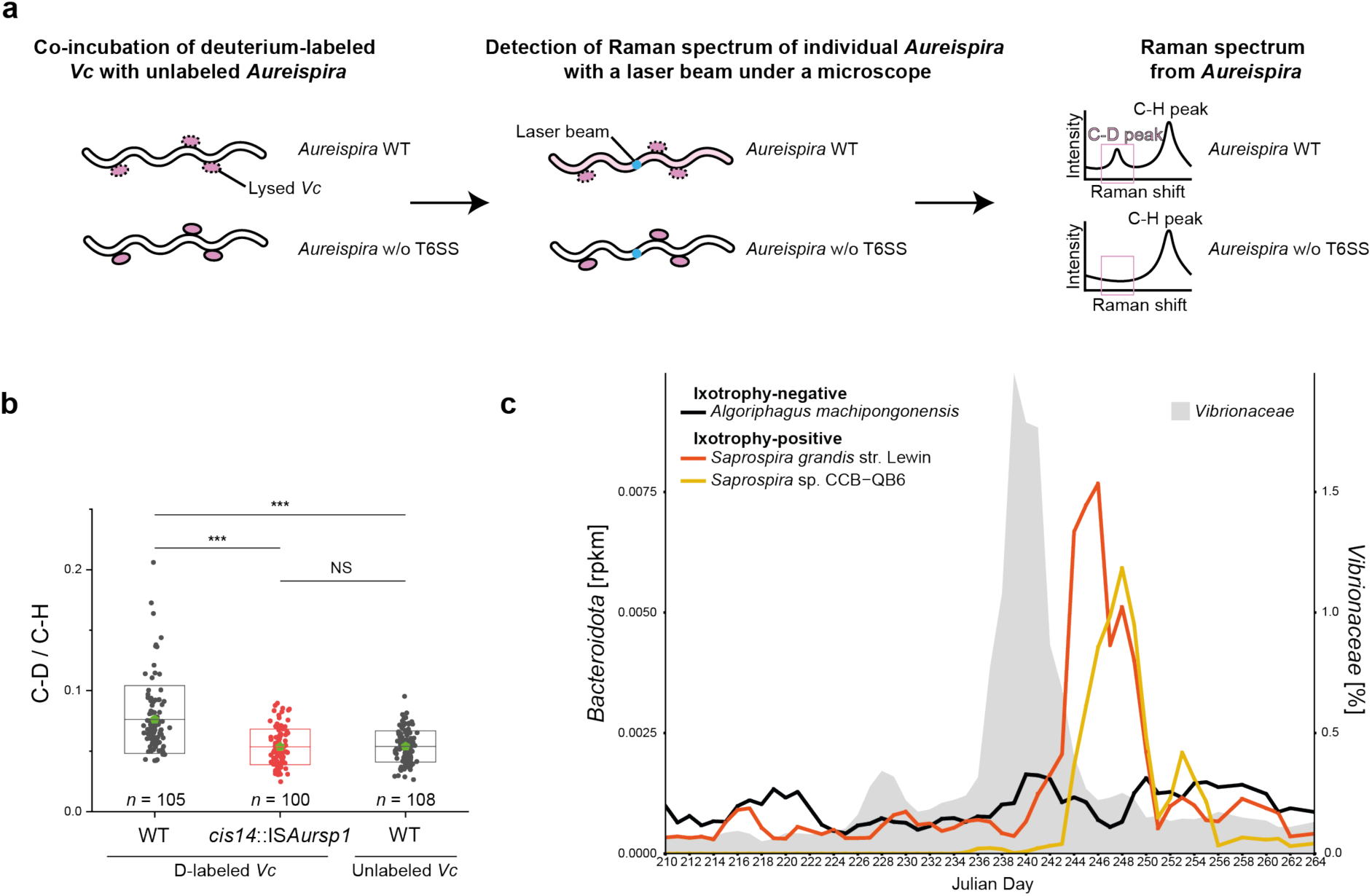
Ixotrophy-positive bacteria take up substrate from labeled prey and their abundance correlates with *Vibrionaceae* in the ocean. **a:** Schematic showing the experimental setup of Raman microspectroscopy measurements for testing substrate uptake by *Aureispira* from *Vc* prey cells. The right panel shows illustrative Raman spectra. See **Fig. S17b/c** for example of experimental spectra. **b:** After co-incubation, the deuterium (D) signal can be detected in T6SS-positive (but not in T6SS-negative) *Aureispira* cells using Raman microspectroscopy. The deuterium signal intensity of *cis14*::IS*Aursp1* cells was comparable with that of *Aureispira* WT co-incubated with unlabeled *Vc*. The box represents mean (also indicated by the light green pentagon) ± s.d. — 0.076 ± 0.028 for WT co-incubated with D–labeled *Vc*; 0.054 ± 0.015 for *cis14*::IS*Aursp1* co-incubated with D–labeled *Vc*; and 0.054 ± 0.013 for WT co-incubated with unlabeled *Vc*. Statistical significance between the samples was assessed using Kruskal-Wallis analysis of variance (ANOVA) followed by post hoc Dunn’s multiple comparison test (***P < 0.001; NS [not significant] P > 0.05). **c:** Analysis of a time-resolved metagenomic dataset (Nahant time series) shows correlation between the abundance of *Vibrionaceae* (gray) and ixotrophic predators (*Saprospira*; orange and yellow) in the environment. The time-lag in the correlation resembles a dynamic predator-prey interaction. In contrast, an ixotrophic-negative *Bacteroidota* (*Algoriphagus machipongonensis*; black) does not show a correlation to the *Vibrio* bloom.

### **The abundance of ixotrophic predators correlates with *Vibrionaceae* in the ocean**

Finally, we asked whether the analysis of an environmental dataset would support the hypothesis that ixotrophy was an important predatory activity in the wild. Since *Vibrionaceae* are common prey for ixotrophic predators (Lewin, 1997; Furusawa *et al*, 2015a; Yeoh *et al*, 2021), we analyzed the Nahant time series, which consists of coastal ocean metagenomes sampled daily over a 93-day period (Martin-Platero *et al*, 2018), for correlated occurrences. We first identified 35 pairs of *Vibrionaceae* and *Saprospiraceae* amplicon sequence variants, whose abundance significantly correlated to each other with a characteristic time-lag indicating coupled dynamics over the entire time series (**Supplementary Material S1**). This behavior is consistent with a predator-prey relationship. Because amplicon sequences typically encompass relatively broad phylogenetic groups, we further analyzed the dynamics using deeply sequenced metagenomes encompassing one very large *Vibrionaceae* bloom from Julian day 238 to 242 (Martin-Platero *et al*, 2018) to test the specificity of the response from ixotrophic predators. This analysis confirmed that the relative abundance of ixotrophy-positive strains (*Saprospira*) increases after the *Vibrionaceae* bloom. In contrast, the abundance of *A. machipongonensis*, which is an ixotrophy-negative *Bacteroidota* strain (Xu *et al*, 2022; Alegado *et al*, 2013), does not respond (**Fig. 5c, Fig. S18**). Overall, these analyses are strongly suggestive that ixotrophy might be an overlooked important predatory lifestyle in this environment.

## Discussion

We propose an integrative model for bacterial ixotrophy shown in **Fig. 6**. Our insights impact different fields of research, which are discussed below.

**Figure 6.**
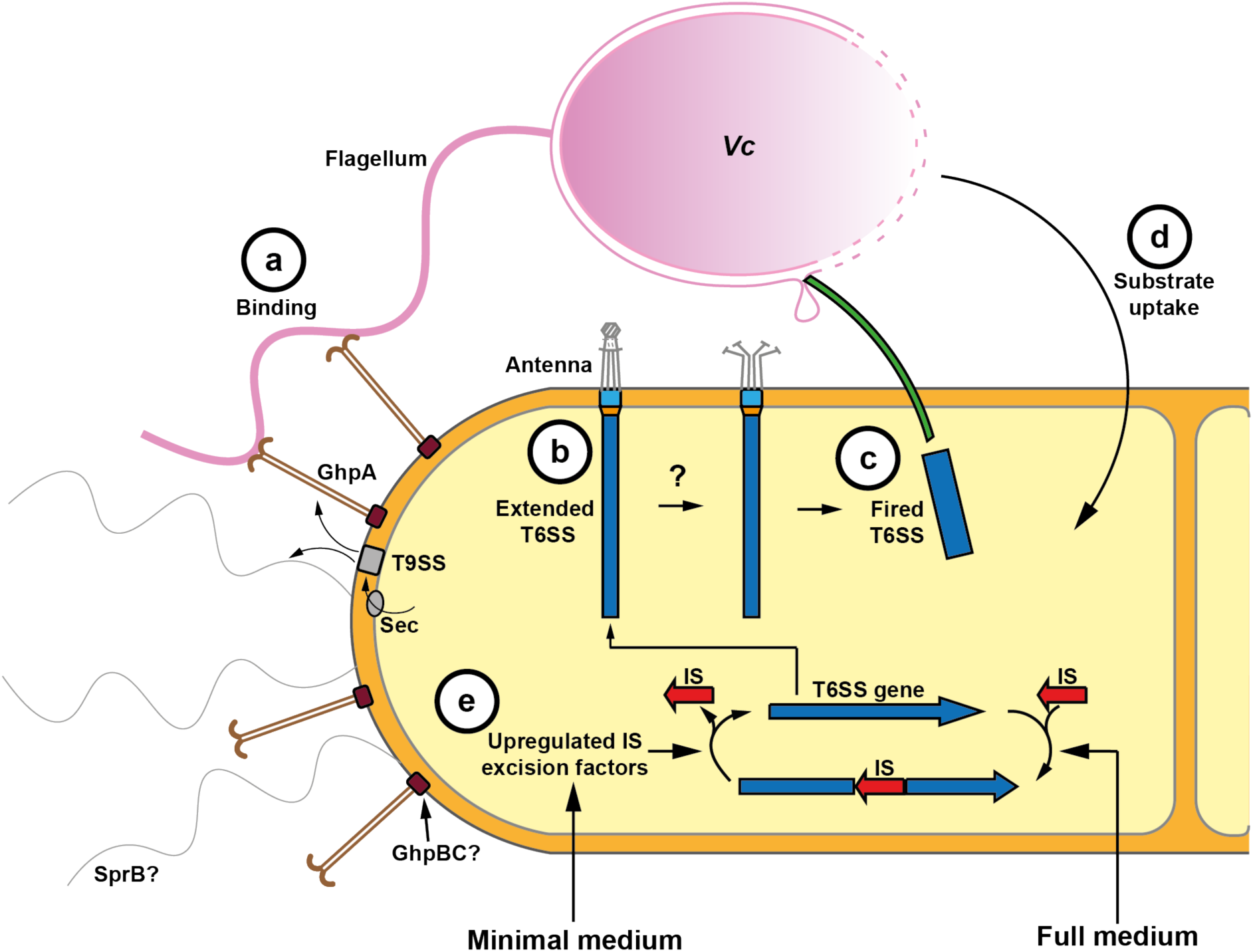
Integrative model of ixotropy. **a:** GhpA is translocated by the Sec/T9SS and assembles into heptameric OM-attached grappling hooks (brown). These grappling hooks mediate the catching of *Vc* flagella in liquid culture. On solid medium, SprB-mediated gliding motility generates cell-cell contacts. **b:** For the subsequent killing of *Vc*, *Aureispira* expresses multiple T6SS*^iv^* (sheath-tube module: blue; baseplate: orange; trans-envelope complex: turquoise). T6SS*^iv^*-associated extracellular antennae (gray; seen in two distinct conformations) may be involved in signal transduction and triggering of firing. **c:** T6SS*^iv^* firing expels the inner tube and propels it into *Vc*, where it causes cell envelope defects and lysis. **d:** Released *Vc* components are taken up by *Aureispira*. **e:** Under nutrient rich conditions, ixotrophy is switched off by endogenous ISs that integrate into T6SS genes. Upon encountering nutrient-limiting conditions, IS excision factors are upregulated, and ixotrophy is re-activated by the excision of ISs.

### The mechanism underlying ixotrophy integrates different functional modules and is conserved

We show that at its core, ixotrophy employs a short-range contact-dependent T6SS for the killing of prey. With the goal of taking up substrate in an aquatic environment, a short-range system is likely advantageous over the secretion of a toxin or an eCIS into the medium. To achieve the fascinating efficiency of killing – being much higher than for any known T6SSs – ixotrophy integrates the action of a T9SS in different ways, depending on the environment. On solid medium (*Aureispira* was isolated from seaweed), T9SS/SprB-mediated motility (likely combined with chemotaxis) allows cells to glide towards prey cells to establish cell-cell contact. Motility has recently been shown to significantly increase the use of short-range weapons (Booth *et al*, 2023). In liquid medium, T9SS/GhpA-mediated catching of prey cells establishes contact that is required for ixotrophy. Biotechnological applications could make use of these different functional modules in the future.

Ixotrophy-like behavior has been reported in filamentous multicellular bacterial species of the *Sparospiria* class from the phylum *Bacteroidota*. Furthermore, a member of the class *Cytophagia* have also been reported to exhibit contact-dependent killing (Svercel *et al*, 2011). Our analysis of available genomes of ixotrophy-positive bacteria (including *Cytophagia*) revealed the presence and coexistence of gene clusters encoding T6SSs, T9SSs, SprB and grappling hook protein GhpA in these organisms (**Table S4**). This indicates a conserved molecular mechanism for ixotrophy and its role in mediating access to nutrients in heterogeneous aquatic environments. In fact, ixotrophy-positive organisms show a broad distribution across diverse sampling sites (**Table S6**). The spectrum of prey includes important aquatic microbes such as *Vibrio* species, cyanobacterial strains, and diatoms, which can undergo dramatic changes in abundance, for instance, during blooms (Martin-Platero *et al*, 2018). The strong correlation between abundances of ixotrophic predators and *Vibrionaceae* in an environmental dataset (**Fig. 5c**) indicates that ixotrophy plays a previously unrecognized role in shaping the marine microbiome. Our study will serve as a framework for studying the impact of ixotrophy on these population dynamics in the future.

### Grappling hooks: establishing cell-cell contacts by an OM-anchored conserved adhesin

Our data reveal grappling hooks as an elegant solution to the challenge for bacteria to establish efficient cell-cell interactions in liquid culture. Interestingly, grappling hook-like surface-anchored cell appendages were previously found in archaea (Moissl *et al*, 2005; Probst *et al*, 2014; Perras *et al*, 2015). These “hami” are believed to be involved in the formation of networks consisting of multiple sister cells. The absence of hami homologs from the *Aureispira* genome indicates convergent evolution, leading to “sticky” hook-like structures that establish cell-cell connections.

*Aureispira* grappling hooks are assembled by protomers of the protein GhpA. In bioinformatic searches, we detected a high abundance of close relatives in other related *Bacteroidota* species (**Table S4**). CryoET imaging of one representative, *Microscilla marina*, confirmed the presence of grappling hooks (**Fig. S19a**), and structure prediction confirmed the characteristic hook-like architecture at the N-terminus (**Fig. S19b**). This result suggests the possibility of broad conservation of the grappling hook structure and its function across ixotrophy-positive bacteria. The formation of a heptameric palm tree-like module at the N-terminal distal end may be critical to expose seven potential binding sites towards the target. This N-terminus shows similarities to adhesins from the Antigen I/II family of *Streptococci* (PDB IDs: 3QE5, 4TSH, 2WQS). Interestingly, many of these adhesins bind Ca^2+^ ions (Hall *et al*, 2014), which is consistent with ixotrophy being dependent on Ca^2+^ (Furusawa *et al*, 2015a).

The grappling hook stem module is characterized by its straight architecture and stability (remaining intact throughout purification). The structural basis for this is likely attributed to several types of interactions between the Ig-like domains of the seven parallel protomers, including hydrogen bonds, disulfide bonds, and glycosylation sites (**Fig. 2j-l, S8, S9**) (Koomey, 2019). Interestingly, GhpA also has similarities to the protein SprB, the key extracellular factor involved in gliding motility in members of *Bacteroidota* (Nelson *et al*, 2008). Even though we lack a high-resolution structure, SprB has been proposed to form filamentous, flexible extracellular appendages (Liu *et al*, 2007; Nakane *et al*, 2013). SprB and GhpA are large proteins (5,898 amino acid residues for *Aureispira* GhpA and 6,497 for *F. johnsoniae* SprB) with Ig-like repeats and a C-terminal T9SS sorting domain. An SprB homolog is also found in the genome of *Aureispira* (WP_052599872.1), which is consistent with the observed gliding motility (**Fig. 1a** and (Furusawa *et al*, 2015a)) and potentially corresponds to the thinner extracellular filaments observed in cryo-tomograms of *Aureispira* cells (**Fig. S20**). Altogether, these data indicate that the straight architecture of GhpA grappling hooks is mediated by heptamerization and interactions between parallel protomers, while SprB may not multimerize and may thus form thinner and more flexible structures. These adaptations are likely due to their different roles in catching prey and motility. Besides understanding a bacterial adhesin with potential biotechnological applications, our structure of GhpA may also serve as a framework for understanding SprB. Next, we discuss a possible pathway for translocating and assembling such supramolecular structures.

### Translocation and anchoring of a T9SS substrate

The T9SS is conserved in the bacterial phylum *Bacteroidota* and plays a critical role in virulence and gliding motility (Lasica *et al*, 2017; Paillat *et al*, 2023). T9SS substrates reach the periplasm by the Sec pathway (signal peptide found in GhpA, **Fig. 2f**). They are then recognized by the T9SS via a C-terminal sorting sequence (present in GhpA, **Fig. 2f**). The core T9SS machinery is composed of an IM rotary motor and an OM-associated complex, both of which are encoded in the *Aureispira* genome (**Table S5**). The OM complex consists of a 36-stranded beta-barrel OM protein (SprA/Sov) that functions as a 70 Å-wide translocon that allows the passage of folded substrate (Lauber *et al*, 2018). Considering the rigidity and the diameter of assembled grappling hooks, the heptamerization of GhpA is likely to happen in the extracellular space. Interestingly, in the pathogen *P. gingivalis*, eight SprA translocons were found to multimerize in the OM (Song *et al*, 2022). It is therefore tempting to speculate that seven GhpA monomers may be concomitantly translocated across the OM and assembled in extracellular space. Here GhpA is potentially anchored to the OM by the homologs of PorE/P (GhpB/C) (Gorasia *et al*, 2022b). Altogether, our high-resolution structure of a T9SS substrate combined with *in situ* imaging will help to understand translocation and anchoring of T9SS substrates in the future.

### T6SS*^iv^*: a contractile injection system with unique structural features kills prey

Besides the role of the T9SS in adhesin translocation, we found that ixotrophy also requires a functional T6SS subtype *iv* for killing the prey. In comparison to other T6SSs, *Aureispira* shows conserved sheath-tube and baseplate modules, while the anchoring to the cell envelope is fundamentally different. The *Aureispira* T6SS features a unique trans-envelope complex and elaborate extracellular antennae in two different conformations. We speculate that these adaptations could function in signal transduction from extracellular space to the T6SS baseplate, thereby triggering firing when a prey cell is in close proximity. This hypothesis is supported by our observation that contracted T6SSs were almost always absent from *Aureispira* cells that were cultured without prey, while contracted T6SSs were frequently seen in cells that were co-incubated with prey (**Fig. S21**).

Another striking feature of the *Aureispira* T6SS is the inner tube. For classical T6SSs, the tube-spike complex is thought to carry effectors that are punctured into the target cell (Shneider *et al*, 2013). The *Aureispira* inner tube may be much more stable and, for the first time, could be seen inflicting cell envelope defects on the prey cell (**Fig. 3l/m**). Differences in tube stability are supported by the observation of expelled inner tubes in purifications of T6SSs from *Aureispira* (**Fig. S11a**) but not from organisms with other T6SSs (Basler *et al*, 2012; Wu *et al*, 2020; Böck *et al*, 2017). It remains open whether the mechanical stress may be complemented by additional effectors. *Aureispira* encodes potential effector candidates that comprise DUF4157 domains, which are associated with CIS effectors in other organisms (Wood *et al*, 2019b; Geller *et al*, 2021; Xu *et al*, 2022). Our LM observation of prey cell rounding upon attacker-prey contacts (**Fig. S1b**) is consistent with the action of a peptidoglycan-degrading effector (Whitney *et al*, 2013).

### Insertion sequence elements: regulation of ixotrophy depending on substrate availability

We found that ixotrophy can be switched off by the insertion of ISs into structural T6SS genes and reactivated by IS excision. Intriguingly, we detected three or six base pair deletions as a result of excision (**Fig. 4e**), an imperfection that is a known phenomenon for IS excision (Kanai *et al*, 2022). While introducing mutations in structural components may in principle cause malfunction of the molecular machine, this mechanism may also promote the evolution of the machinery.

Although ISs are considered selfish genes, they can be utilized by different pathogens for the regulation of capsule production (*Acinetobacter baumannii* (Whiteway *et al*, 2022)), for the regulation of metabolic pathways (Shiga toxin-producing *Escherichia coli* (Nakamura *et al*, 2022)) and the variation of O1 antigen (*Vibrio cholerae* (Nesper *et al*, 2000)); however, how environmental factors regulate the excision of ISs is not well understood. For ixotrophy, our data suggest that the levels of available nutrients may regulate IS activity. In particular, we show that the transcription of factors that are critical for IS excision (*recB*/*iee*) is upregulated under starvation conditions. Cultivation in full medium, on the other hand, leads to deactivation of T6SS by IS integration. The incentive for bacteria to evolve such a genomic switch, could be the possibility to deactivate the expression of dispensable and energy-costly molecular machineries. Recent studies on *Vibrio*, *Pseudomonas*, and *Bacteroides* strains suggest that T6SS activity does in fact come with a fitness cost and it may be beneficial to regulate its activity (Drebes Dörr *et al*, 2022; Septer *et al*, 2023; Perault *et al*, 2020; Robitaille *et al*, 2023).

## Materials and Methods

### Bacterial culture conditions

*Aureispira* sp. CCB-QB1, *Vibrio campbellii* and their derivatives were cultured in artificial seawater medium [ASWM; 2.4% Instant Ocean (Aquarium Systems), 10 mM HEPES, 0.5% peptone, pH 7.6] at 30 °C with constant shaking at 200 rpm or on ASWM plates solidified with 1.5 % agar. Marine agar and marine broth (Condalab) were used for culturing *Vc*. *Vibrio cholerae* and *Escherichia coli* were grown in Luria-Bertani (LB) broth or on LB agar plate solidified with 1.5% agar at 37 °C. All strains used in this study are listed in **Table S7**.

### Killing assays

*Aureispira* and prey were cultured overnight in ASWM. In the morning, the cells were concentrated and resuspended in low nutrient ASWM (L-ASWM; 2.4% Instant Ocean, 10mM HEPES, 0.05% peptone, pH 7.6) to an OD_600_ of 1.0. Cells were starved in L-ASWM for 2 h. After starvation, *Vc* cultures were mixed in a 1-to-1 ratio with differently treated *Aureispira* cultures: i) untreated *Aureispira* culture (referred to as ‘Live *Aureispira’*); ii) heat-inactivated *Aureispira* culture, which was incubated at 96 °C for 10 min (referred to as ‘Boiled *Aureispira*’); iii) *Aureispira* culture supernatant filtered through a 0.22 μm filter (referred to as ‘*Aureispira* supernatant’); iv) without the addition of any *Aureispira* cells (referred to as ‘*Vc* only’). Mixtures were co-cultured for 6 h or as indicated. This was followed by serial dilutions and plating or spotting onto marine agar plates for quantification and comparison.

For assays on solid agar plates, OD_600_ of *Aureispira* and prey overnight cultures were concentrated and resuspended in ASWM to an OD_600_ of 1.0. 5 μL of differently treated *Aureispira* (see above) and *Vc* cultures were mixed in a 1:1 ratio, applied on L-ASWM agar plates and incubated for 24 h or as indicated. After co-incubation, cells were scraped from the agar plate with an inoculation loop and resuspended in 200 μL L-ASWM, followed by serial dilution and plating or spotting onto marine agar plates for quantification and comparison.

### Light microscopy imaging

To image the ixotrophy behavior, *Aureispira* and *Vc* were cultured separately in ASWM, and the cultures were concentrated and resuspended in L-ASWM to an OD_600_ of 1.0. Afterwards, cells were starved in L-ASWM for 2 h. 5 μL of *Aureispira* culture and 5 μL of *Vc* culture were mixed on an 0.5% agarose pad made from L-ASWM. The samples were imaged with a 100x oil immersion objective on a Leica Thunder Imager 3D Cell Culture equipped with a Leica DFC9000 GTC CMOS camera (2,048 × 2,048 pixels, 65 x 65 nm pixel size). For the visualization of DNA, cells were stained with DAPI at a final concentration of 10 μg/μL for 30 min at room temperature in the dark. For time-lapse imaging, the samples were imaged either with the Leica Thunder microscope mentioned above or with a Zeiss wide-field microscope equipped with a Hamamatsu ORCA-ER digital camera. Images were recorded every 5 s over a time course of 30 min. For imaging in liquid culture, *Aureispira* and *Vc* WT and mutants were starved for 2 h in L-ASWM. 800 μL *Aureispira* culture and 80 μL *Vc* culture were mixed in a 35 mm uncoated μ-Dish (Ibidi) and immediately imaged with the microscope with a 40x objective. All images were further analyzed and processed with Fiji (Schindelin *et al*, 2012).

### Mutagenesis of *Vc*

To delete the flagellin genes in *Vc*, 800 bp-long sequences located upstream and downstream of the *Vc* flagellins were amplified with Q5 2X master mix (New England Biolabs) and ligated into the restriction site XhoI and SphI in pDM4 followed by transformation of the plasmid into *E. coli* SM10 λ pir by electroporation. The plasmid was inserted into the *Vc* genome by conjugation with *E. coli* SM10 λ pir as described previously (Simpson *et al*, 2019). Briefly, overnight cultured *Vc* and *E. coli* SM10 λ pir harboring pDM4 plasmid were mixed, and 10 μL of the cultures were pipetted onto a marine agar plate and incubated at 30 °C overnight. On the following day, *Vc* conjugants were selected with marine agar plates supplemented with 10 μg/mL chloramphenicol and 50 U/mL polymyxin B. A single colony of a *Vc* conjugant was picked and cultured overnight in marine broth and subsequently plated in the morning onto marine agar plates containing 10% sucrose and 50 U/mL polymyxin B to select for the colonies that lost the suicide plasmid. Colonies were screened for the presence of the suicide plasmid and deletion of the gene with PCR. The process was repeated twice to delete all six flagellin genes separated into two loci in the *Vc* genome. Used primers are listed in **Table S8**.

### Purification of flagella from *Vc*

Flagella of *Vc* were purified as described earlier (Yang *et al*, 1977) with some modifications. Briefly, a 5 mL *Vc* overnight culture in ASWM was pelleted and resuspended in 1 mL of artificial seawater [2.4% Instant Ocean (Aquarium Systems), 10mM HEPES, pH7.6] followed by vigorous vortexing for 2 min to shear off flagella. Intact *Vc* cells were removed by centrifugation at 8,000 g for 10 min at 4 °C, and the supernatant was subjected to ultracentrifugation at 100,000 g for 1 h at 4 °C to pellet the flagella. The flagella pellets were resuspended in 100 μL artificial seawater.

### Purification of grappling hooks

*cis14*::IS*Aursp1* was inoculated in 10 mL of ASWM and grown overnight at 30 °C and 200 rpm. Afterwards, the culture was transferred to 200 mL ASWM and grown overnight at 30 °C and 200 rpm before being transferred again into 2 L ASWM and grown overnight at 30°C and 200 rpm. Finally, the cells were harvested (7,000 g, room temperature, 20 min), and the cell pellet was thoroughly resuspended in 48 mL grappling hook shearing buffer (150 mM NaCl, 20 mM HEPES, pH7.5, 1x cOmplete EDTA-free protease inhibitor tablet) until all clumps were dissolved. The grappling hook shearing protocol was adapted from Gaines, M. C. et al. (Gaines *et al*, 2022). Briefly, the vial with the cell suspension was connected to a peristaltic pump and, under gentle stirring using a magnetic stirrer, continuously passed through two needles (0.8 mm x 40 mm, 4 °C, 3 h, Microlance 3, VVR and 0.5 mm x 16 mm, 4 °C, overnight, Microlance 3, VVR) at 25 rpm. The sheared suspension was then centrifuged (21,000 g, 4 °C, 20 min) to remove intact cells and debris.

The supernatant was filtered through a 0.45 μm filter (Spritzenfilter Filtropur, Sarstedt AG) and subjected to ultracentrifugation on a sucrose cushion. 1 mL Sucrose Cushion Buffer (150 mM NaCl, 20 mM HEPES, pH 7.5, 50% (w/v) sucrose) was filled into ultracentrifuge tubes (Polypropylene 10 ml Tube, Beckman Coulter), the sheared grappling hooks were gently layered on top, and the samples were centrifuged (250,000 g, 4 °C, 3 h, MLA-80 rotor). Taking care not to disturb the cushion at the bottom, all but the bottom-most 1.5 mL were discarded. The remaining solutions were pooled and centrifuged again (21,000 g, 4 °C, 20 min). The supernatant was diluted to 48 mL with grappling hook buffer (150 mM NaCl, 20 mM HEPES, pH 7.5). After an additional round of ultracentrifugation (250,000xg, 4 °C, 3 h, MLA-80 rotor), the resulting pellets were soaked in 75 μL grappling hook buffer overnight at 4 °C. Afterwards, the pellets were gently resuspended, centrifuged (21,000 x g, 4 °C, 20 min) and the supernatant was used for sucrose gradient ultracentrifugation. To this end, 2.5 mL 50 % (w/v) and 3 mL 10 % (w/v) sucrose gradient buffer (150 mM NaCl, 20 mM HEPES, pH 7.5 with 50 % or 10 % sucrose, respectively) were gently layered on top of each other in ultracentrifuge tubes (Ultra-Clear 5 mL Tube, Beckman Coulter) and a continuous gradient was generated using a gradient maker (Gradient Master, BIOCOMP). The top-most 200 μL of the gradient was discarded, and 300 μL of the grappling hook sample was gently loaded onto the gradient. The sample was then centrifuged (250,000 g, 4 °C, 3 h, deceleration = coast, SW55-Ti rotor) and 11 x 0.5 mL fractions were collected (F1-F11), with F11 being the top-most. F5 was found to contain the most grappling hooks, and thus, it was diluted to 6 mL with grappling hook buffer and subjected to another round of ultracentrifugation (250,000 g, 4 °C, 3 h, MLA-80 rotor). The pellet was soaked overnight at 4 °C in 100 μL grappling hook buffer, gently resuspended and stored at 4 °C until further use.

### Plunge freezing for cryoET and single particle cryoEM

To image intact *Aureispira* cells by cryoET, the OD_600_ of an *Aureispira* overnight culture was adjusted to 1.0 and sub-cultured in L-ASWM for 2 h. 3.5 μL of the culture were applied onto glow-discharged copper EM grids (R2/1, Quantifoil), automatically blotted and plunged into liquid ethane/propane (Tivol *et al*, 2008) using a Vitrobot Mark IV (Thermo Fisher Scientific) (Iancu *et al*, 2006). Using a Teflon sheet on one side, samples were blotted only from the back for 8-10 s.

To image attacker-prey contacts, overnight cultures of *Aureispira* and *Vc* were starved in L-ASWM for 2 h. Cell cultures were concentrated to an OD_600_ of 5-10 by centrifugation at 5,000 g for 4 min and resuspended in L-ASWM. Afterwards, concentrated *Aureispira and Vc* cultures were mixed and incubated at room temperature for 2-8 min before plunge freezing. 4 μL of the cell mixture was applied three times onto glow-discharged copper EM grids (R2/2, Quantifoil), followed by plunge freezing into liquid ethane/propane. Using a Teflon sheet on one side, samples were blotted from the back only for 6 s.

For single particle cryoEM of grappling hooks, copper EM grids (Cu 200 R 2/1, Quantifoil) were used. 3.5 μL purified grappling hook solution was applied onto the grid, and the grid was automatically blotted with filter papers from both sides and plunged into a liquid ethane/propane mixture using a Vitrobot Mark IV. Plunge-frozen grids were stored in liquid nitrogen.

### CryoFIB milling

Lamellae through plunge-frozen *Aureispira-Vc* mixture were obtained by automated sequential cryoFIB milling as described previously (Zachs *et al*, 2020). Briefly, EM grids were clipped into FIB milling autoloader-grids (Thermo Fisher Scientific) and mounted onto a 40° pre-tilted grid holder (Medeiros *et al*, 2018) (Leica Microsystems GmbH) using a VCM loading station (Leica Microsystems GmbH). For all grid transfers under cryo-conditions, a VCT500 cryo-transfer system (Leica Microsystems GmbH) was used. Grids were sputter-coated with a ∼4 nm thick layer of tungsten using an ACE600 cryo-sputter coater (Leica Microsystems GmbH) and afterwards transferred into a Crossbeam 550 FIB-SEM dual-beam instrument (Carl Zeiss Microscopy), equipped with a copper-band cooled mechanical cryo-stage (Leica Microsystems GmbH). The gas injection system (GIS) was used to deposit an organometallic platinum precursor layer onto each grid. Grid quality assessment and targeting of the cells were done by scanning EM (SEM) imaging (3-5 kV, 58 pA). Coordinates of chosen targets were saved in the stage navigator, and milling patterns were placed onto targets’ FIB image (30 kV, 20 pA) using the SmartFIB software. To mill 10 μm wide and ∼250 nm thick lamellae, a total of four currents were used, and currents were gradually reduced according to lamella thickness [rough milling (700 pA, 300 pA, and 100 pA) and polishing (50 pA)]. After the milling session, the grids were unloaded and stored in liquid nitrogen.

### CryoET

Micrographs of *Aureispira* samples were recorded on a Titan Krios 300 kV FEG transmission electron microscope (Thermo Fisher Scientific) equipped with a Quantum LS imaging filter (slit width 20 eV) and K2 direct electron detector (Gatan). A low magnification overview of the grid was recorded using SerialEM (Mastronarde, 2005; Schorb *et al*, 2019). Tilt series were collected automatically with SerialEM and covered an angular range from –50° to +70° or - 60° to +60° with 2° increments for cryoFIB-milled lamellae and *Aureispira* cells, respectively. The defocus was set to -8 μm for cryoFIB-thinned lamellae and to -5 to -8 μm for intact *Aureispira* cells. The total dose of a tilt series accumulated 160 e^-^/Å^2^ and the pixel size at the specimen level was 3.46 Å. CryoET data of *M. marina* were recorded on a Tecnai Polara 300kV TEM (FEI) equipped with post-column GIF 2002 imaging filter (slit width 20 eV) and K2 Summit direct electron detector (Gatan). Tilt-series were collected from -60° to +60° with 1° increment at a pixel size of 4.9 Å using UCSF tomography (Zheng *et al*, 2007) and with a defocus of -8 μm and a cumulative electron dose of 180 e^-^/Å^2^.

### Tomogram reconstruction and subtomogram averaging

Tilt series were drift-corrected using alignframes and CTF correction, and three-dimensional (3D) reconstructions were generated using the IMOD package (Kremer *et al*, 1996; Mastronarde, 2008). For visualization, tomograms were filtered using the tom_deconv deconvolution filter (Tegunov & Cramer, 2019). The deep learning-based segmentation tool Dragonfly (Heebner *et al*, 2022) was used for the segmentation of IsoNet-corrected (Liu *et al*, 2022) tomograms.

For subtomogram averaging of T6SS baseplates and trans-envelope complexes, T6SSs were manually identified in individual tomograms, and their long axes from baseplate toward the sheath were modelled with open contours in 3dmod (Mastronarde, 2008) to generate model points, the initial motive list and particle rotation axes. Initial averages that were used as subsequent first references were created with PEET (Nicastro Daniela *et al*, 2006). Afterwards, particles were extracted, aligned and averaged with Dynamo (Castaño-Díez *et al*, 2012). The 4 × 4-binned particles were aligned for 12 iterations (box size 88 × 88 × 88 pixels^3^). Afterwards the datasets were split in half for gold-standard Fourier shell correlation (FSC) calculations, and masks focusing either on the baseplate or the trans-envelope complex were used for alignments, respectively. Independent alignments were repeated for eight iterations using 2 × 2-binned tomograms (box sizes 176 × 176 × 176 pixels^3^). The set of particles was cleaned according to cross-correlation values, and another 8 alignment iterations were performed followed by a final alignment with tight masks generated by Relion’s mask generator (Zivanov *et al*, 2018) based on the density map from the previous averaging round with an additional extension of four pixels at the boundaries. A total of 560 T6SSs were initially selected from 146 tomograms. The final, CC-cleaned and C6-symmetrized average, resulted from 399 baseplate particles and 334 trans-envelope complex particles with a box size of 176 × 176 × 176 pixels^3^ at a pixel size of 6.91 Å.

For subtomogram averaging of the extracellular antenna, 61 closed antennae and 21 open antennae were picked manually from tomograms with Dynamo, and low-resolution averages were calculated to serve as initial reference. The cropping points from the averaging of T6SSs were now moved 65 pixels towards the extracellular space, and 4 × 4-binned particles containing both closed and open antennae were aligned for 12 iterations with initial reference of the closed antennae or the open antennae (box sizes 120 × 120 × 120 pixels^3^). The set of particles was separated into a group either containing only closed antennae or only open antennae according to cross-correlation values. The split datasets were subjected to another eight alignment iterations. Afterwards, each dataset was split in half for gold-standard FSC calculations, and independent alignments were repeated for 8 iterations using 2 × 2-binned tomograms (box sizes 240 × 240 × 240 pixels^3^) followed by a final alignment with tight masks generated by Relion’s mask generator (Zivanov *et al*, 2018) based on the density map from the previous averaging round with an additional extension of four pixels at the boundaries. The final, C6-symmetrized average, resulted from 152 closed antenna particles and 76 open antennae particles with a box size of 240 × 240 × 240 pixels^3^ at a pixel size of 6.91 Å.

For subtomogram averaging of the grappling hook distal end, the tips of the grappling hooks were manually picked from tomograms. Particle extraction, alignment and averaging were done with Dynamo (Castaño-Díez *et al*, 2012). The 4 × 4-binned particles were aligned for 12 iterations (box size 40 × 40 × 40 pixels^3^). Afterwards the dataset was split in half for gold-standard FSC calculations. Independent alignments were repeated for 8 iterations using 2 × 2-binned tomograms (box sizes 80 × 80 × 80 pixels^3^). The final, C7-symmetrized average, resulted from 425 grappling hooks particles from 71 tomograms with a box size of 80 × 80 × 80 pixels^3^ at a pixel size of 6.91 Å.

For subtomogram averaging of the grappling hook basal body, the particles were manually identified in individual tomograms, and their long axes from the OM toward the grappling hook distal ends were modelled with open contours in 3dmod (Mastronarde, 2008) to generate model points, the initial motive list and particle rotation axes. An initial low-resolution average used as a subsequent reference was calculated with PEET with 72 particles (Nicastro Daniela *et al*, 2006). Afterwards, particles were extracted, aligned and averaged with Dynamo (Castaño-Díez *et al*, 2012). The 4 × 4-binned particles were aligned for 8 iterations (box size 30 × 30 × 30 pixels^3^). The refined table was then used for another round of particle cropping with a bigger box size (72 × 72 × 72 pixels^3^). These particles were again aligned for another eight iterations. The final, C7-symmetrized average, resulted from 285 grappling hook basal body particles from 96 tomograms with a box size of 72 × 72 × 72 pixels^3^ at a pixel size of 13.82 Å.

For all the subtmogram averages, except for the one of grappling hook basal body, gold-standard FSC curves for resolution estimation were calculated with Dynamo after alignment of the half-maps in UCSF Chimera (Pettersen *et al*, 2021) using the fit map function. No FSC-curve was calculated for the grappling hook basal body as it remained at low resolution.

### Single particle cryoEM data collection and data processing

Data was collected on grids with appropriate ice thickness and particle distribution. 29,238 movies were collected on a Titan Krios G4 cryo-TEM (Thermo Fisher Scientific, ScopeM, Zürich) operated at 300 kV, equipped with a Bio-Continuum imaging filter (Gatan) and a K3 direct electron detector (Gatan). Automatic data collection was carried out using the Thermo Fisher Scientific EPU software. The scheme of 4 shots per hole was applied during data collection in counting mode, at a pixel size at specimen level of 1.065 Å/px and a defocus range of -1 to -2.5 μm. The total dose was 50 e^-^/A^2^ over 40 frames and 2 s exposure time.

The movies were motion corrected using MotionCor2 (Zheng *et al*, 2017). The resulting micrographs were imported into CryoSPARC v.4 (Punjani *et al*, 2017) and the contrast transfer function (CTF) estimated using the patch CTF estimation tool. Grappling hook stem particles were picked manually, with a box size of 300 pixels, and averaged using 2D classification. An initial rise of roughly 125 Å was estimated from the 2D class averages. The 3 best classes were then used as references for the filament tracer tool (filament diameter = 100 Å, inter-box distance = 130 Å), which picked 2,452,192 particles. The particles were extracted with a box size of 300 Å, binned to a pixel size of 3.8 Å. During 2D classification, re-centering after each iteration was not allowed and a mask with a diameter of 90 % of the box size was applied. After two rounds of classification, during which bad classes and classes of the grappling hook distal/proximal ends were removed, 1,309,380 good particles of the stem remained.

The particles were subjected to helical refinement, using real-space windowing of 0.75-0.9 and a featureless cylinder with a diameter of 90 Å as reference, without imposing any kind of symmetry. The refinement did not converge into a reasonable structure, so C7 symmetry was applied, reasoning that the symmetry of the subtomogram average of distal hook region of the grappling hooks (**Fig. S3**) would be conserved in the stem. Imposing C7 symmetry quickly caused the refinement to generate a more reasonable initial reference. From then on, C7 symmetry was applied throughout the refinement. It was attempted to identify helical parameters, but no convincing values were obtained, and the refinement was carried out without imposing helical symmetry. 3D classification, using real-space windowing of 0.75-0.9, was carried out with a target of 5 classes and a maximum resolution of 7 Å. Hard classification was applied to prevent particles from having partial assignment to more than one class, leading to better 3D classes. The 2 classes with the most particles were refined individually. Particles were re-extracted at a pixel size of 1.68 Å and subjected to helical refinement, applying a mask covering the central 50 % of the map, 2D classification and 3D classification with a target resolution of 4 Å. A 50 % mask was then applied to all other refinement jobs. An iterative process of 3D classifications, with a target resolution of 3.5 Å, helical refinement, non-uniform refinement (Punjani *et al*, 2020), global and local CTF refinement on unbinned particles resulted in 6 structures with resolutions between 3.4 and 3.7 Å. A graphical depiction of the workflow is shown in **Fig. S5**. Global resolution was determined by gold standard fourier shell correlation (GSFSC), as estimated by CryoSPARC at a threshold of 0.143 (**Fig. S7**). Local resolution was estimated at a GSFSC threshold of 0.143 across the entirety of the map and visualized in ChimeraX (Pettersen *et al*, 2021) (**Fig. S7**). Finally, non-uniform refinement was run again on each of the final maps without imposing symmetry of any kind (C1). Unless stated otherwise, all data processing steps were carried out using the CryoSPARC v4 default settings.

For data processing of the grappling hook distal/proximal ends, of the 2,452,192 particles picked by the filament tracer, terminal segments were identified visually and re-extracted with a box size of 800 pixel and a binned pixel size of 4.26 Å, resulting in 78,681 particles. 2D classes were inspected manually and divided into two categories, referred to as "base" and "neck" classes, which were processed independently. Based on the previously obtained stem structures, C7 symmetry was used in the refinement of both neck and base classes. Ab-initio refinement was performed to obtain initial volumes, which were improved upon through an iterative process of 2D classification and homogeneous, non-uniform (Punjani *et al*, 2020) and CTF refinement (all using the parameters described above). At the end of this process, the neck structure had reached a global resolution of 3.77 Å, while the base structure reached a global resolution of 3.73 Å as estimated by GSFSC in CryoSPARC (Punjani *et al*, 2017) at a threshold of 0.143 (**Fig. S7**). A graphical depiction of the workflow is shown in **Fig. S6**. Local resolution was estimated at a GSFSC threshold of 0.143 across the entirety of the map and visualized in ChimeraX (Pettersen *et al*, 2021) (**Fig. S7**). Unless stated otherwise, all data processing steps were carried out using the CryoSPARC v4 default settings. Noise was removed from the maps shown in Figures using the "hide dust" tool in ChimeraX (Pettersen *et al*, 2021).

### Structural modeling

The sharpened cryoEM maps and the corresponding sequence in FASTA format were used as input for ModelAngelo 1.0 (Jamali *et al*, 2023) using the "model_angelo build" option. The output models were inspected in ChimeraX (Pettersen *et al*, 2021), and the best fitting protein chain was identified manually. The model was then manually curated and adjusted in Coot (Emsley *et al*, 2010) (accessed through SBGrid (Morin *et al*, 2013)). The models were then subjected to iterative real-space refinements against related density maps using RosettaCM (Song *et al*, 2013) and ‘phenix.real_space_refine’64. The final models were evaluated using “phenix.molprobity” (Adams *et al*, 2010) (**Table S9**), and the correlations between models and the corresponding maps were estimated using ‘phenix.mtriage’ (Adams *et al*, 2010). To analyze potential glycosylation sites, the cryoEM density maps were masked with a 3 Å zone around the atomic model. This was used in **Fig. 2j** and **S9** to color all densities, which are more than 3 Å distant from the center axis of the atomic models in red. Molecular graphs were generated with Chimera (Pettersen *et al*, 2004) and ChimeraX (Pettersen *et al*, 2021).

### Purification of T6SSs

For purification of *Aureispira* T6SSs, a total of 50 mL *Aureispira* culture grown in ASWM were pelleted, resuspended in 3 mL of lysis buffer (150 mM NaCl, 50 mM Tris-HCl, 0.5× CellLytic B (Sigma-Aldrich), 1% Triton X-100, 200 μg/mL lysozyme, 50 μg/mL DNAse I, 1 mM phenylmethylsulfonyl fluoride, pH 7.4) and incubated at 37 °C for 1 h. Cell debris was removed by centrifugation (15,000 g, 15 min, 4 °C), and cleared lysates were subjected to ultracentrifugation (150,000 g, 1 h, 4 °C). Pellets were resuspended in 150 μL of resuspension buffer (150 mM NaCl, 50 mM Tris-HCl, supplemented with cOmplete EDTA-free protease inhibitor cocktail [Roche], pH 7.4). For protein identification, the samples were sent to the Functional Genomic Center Zurich (FGCZ) for mass spectrometry analysis. A total of 100 μL of T6SS preparation was digested with 5 μL of trypsin (100 ng/μL in 10 mM HCl) and microwaved for 30 min at 60 °C. The sample was dried and dissolved in 20 μL of 0.1% formic acid, diluted 1:5 and transferred to the autosampler vials for liquid chromatography with tandem mass spectrometry analysis. A total of 1 μL was injected. Database searches were performed by importing the acquired mass spectrometry data into the PEAKS Studio (Bioinformatics solutions), and the data were searched against the NCBI protein database restricted to proteins from the *Aureispira* genus.

### Negative stain EM

4 μL of sample solution was applied onto a glow-discharged, carbon-coated copper grid for 60 s, washed twice with water, and stained with 1% phosphotungstic acid for 20 s. The grids were examined using a Morgagni transmission electron microscope (Thermo Fisher Scientific) operated at 80 kV.

### Whole genome sequencing

Genomic DNA was extracted with Wizard Genomic DNA Purification Kit (Promega) from 1 mL of overnight *Aureispira* culture grown in ASWM. The DNA samples were sent to the Functional Genomic Center Zurich for long reads whole genome sequencing.

1 µg of bacterial DNA for each derivative was used for PacBio library preparation following PacBio protocol for multiplexing bacterial genomes for sequencing. The multiplexed PacBio libraries were further sequenced on a PacBio Sequel instrument using a single PacBio 1M SMRT cell, Sequel® Binding Kit 3.0 and Sequencing Primer v4.

PacBio long reads were first de-multiplexed using the “Demultiplex Barcodes” App in the PacBio Software SMRT Link (v 10.2.0.133434). De-multiplexed reads were then mapped to the reference genome (GCF_000724545.1_ Aureispira_sp._CCB-QB1), and small variants were identified using the “Resequencing” App in the SMRT Link. Variants were functionally annotated against the Aureispira_sp._CCB-QB1 gene models using SnpEff (v4.3). For large structural variant identification, de-multiplexed reads were aligned to the reference genome using “pbmm2” in the SMRT Link command line tools (v 10.2.0.133434). Aligned reads were analyzed using “pbsv” in the same collection of SMRT Link command line tools for identification of large insertion, deletion, inversion, translocation and duplication.

De novo assembly was performed using Canu (v2.0) and the “Microbial Assembly” App in SMRT Link (v 10.2.0.133434). Assembled contigs were compared to the reference genome using nucmer and dnadiff in MUMmer (v4.0.0beta2).

### Stable isotope probing – single-cell Raman microspectroscopy

To investigate the uptake of *Vc* components by *Aureispira*, the stable isotope probing (SIP)– Raman technique was employed. To label prey cells with deuterium (D), *Vc* cells were cultured in 50% D_2_O-containing ASWM overnight and subsequently sub-cultured in 50% D_2_O-containing L-ASWM for 2 h. The same procedures were repeated with *Vc* cells cultured in conventional H_2_O ASWM or L-ASWM, respectively, to be used as unlabeled prey cells. All *Vc* cells were washed once with H_2_O L-ASWM before mixing with an overnight *Aureispira* culture, which was starved in L-ASWM for 2 h. The *Aureispira-Vc* 1-to-1 mixture was then co-cultured at 30 °C with 200 rpm for 1 h. A 5-µL droplet containing each sample was placed on an aluminum-coated slide (EMF Corp., USA) and dried at 30 °C for 10 min. The samples were then washed using Milli-Q (MQ) water to remove traces of the medium, followed by air blowing to remove residual MQ water (Lee *et al*, 2021).

The samples were measured using a confocal Raman microspectroscope (LabRAM HR Evolution, Horiba Scientific, France). The system is based on an upright microscope (BxFM, Olympus), integrated with components for Raman measurements, including an objective (MPlan N 100×, 0.90 NA, Olympus), a 100-µm confocal pinhole, a 300-lines/mm diffraction grating (blazed at 600 nm), and a spectrometer (back illuminated deep-depleted CCD; 1,024 × 256 pixels). A 532-nm laser at 10 mW (continuous wave neodymium-doped yttrium aluminum garnet; CW Nd: YAG) was illuminated onto the individual cells, and their Raman spectra were measured with 10 s exposure time.

The measured Raman spectra were processed using a code built in-house (Lee *et al*, 2021). Smoothing (de-noising) and baseline subtraction were conducted using the Savitzky-Golay filter (polynomial order and frame length: 3 and 9) and a polynomial-based algorithm (polynomial order and threshold: 1 and 0.001), respectively. To evaluate the deuterium labeling status of *Aureispira*, the Raman intensity for the carbon-deuterium peak (C-D at 2,100-2,300 cm^-1^; integrated intensity, I2,100-2,300) was normalized by that for a lipid peak (C-H at 2,800-3,100 cm^-1^; integrated intensity, I2,800-3,100). The C–H peak, a proxy for cell biomass, was used for this normalization to consider the heterogeneity of biomass between cells and to compensate the precision difference of the focus of the Raman laser beam on individual cells.

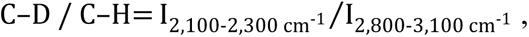

where *I* represents integrated intensity within a spectral region specified.

### RNA isolation and quantitative reverse transcription PCR (qRT-PCR)

An *Aureispira* overnight culture was sub-cultured into artificial seawater medium without any nutrients (0-ASWM) [2.4% Instant Ocean (Aquarium Systems), 10mM HEPES, pH7.6] or ASWM (rich medium) with or without supplement of *Vc* for 6 h. 1 mL of bacterial cell culture was pelleted, and the total RNA was extracted with the SPLIT RNA extraction kit (Lexogen) followed by DNase treatment with the TURBO DNase kit (Thermo Fisher). 1 μg of the RNA was reverse transcribed into cDNA with the iScript Advanced cDNA Synthesis kit (Bio-rad). 20 μL of the end products were diluted by adding 180 μL nuclease-free water and stored at -20 °C until further analysis.

5 μL of each sample was used for qRT-PCR. cDNA samples were mixed with 2.5 μL of 5 μM forward primer, 2.5 μL of 5 μM reverse primer and 10 μL of KAPA SYBR FAST (KAPA Biosystems). The qRT-PCR was done in CFX96 Real-Time System C1000 Touch Thermal Cycler (Bio-Rad). The cycling program was as follows: 1 cycle of 95 °C for 5 min, 40 cycles of 95 °C for 10 s, 62 °C for 10 s, 72 °C for 12 s, followed by a melt curve. For each experiment, 3 technical replicates were conducted and repeated with 3 biological replicates. Data processing and relative gene expression were calculated with Bio-Rad CFX Maestro (Bio-Rad).

### Evaluation of ISs excision upon starvation

*Aureispira* WT and its T6SS-negative derivatives were cultured in 10 mL ASWM overnight and sub-cultured into 50 mL ASWM on the next day. The 50 mL *Aureispira* ASWM culture was pelleted and resuspended in 50 mL 0-ASWM. On the same day, 50 mL *Vc* overnight cultures in ASWM were pelleted, resuspended in 2 mL 0-ASWM, and added to each *Aureispira* 0-ASWM culture. The OD_600_ of each culture was measured every day, and OD_600_ was adjusted to 2 by the addition of *Vc* cells. PCR was performed after the T6SS-negative derivatives showed a decreased OD_600_. Bands from the PCR products were cut out after agarose gel electrophoreses, purified, and further sent to Microsynth AG for Sanger sequencing to confirm the excision of the ISs.

### Metagenomic abundance analysis

Sea surface water was collected from Canoe Cove, Nahant, Massachusetts, USA (Lat: 42° 25 ′ 10.6″ N, Lon: 70° 54′ 24.2″ W), from 23^rd^ of July (Julian day 204) to 23^rd^ of October 2010 (Julian day 296) (Martin-Platero *et al*, 2018).

(SRR5175890 to SRR5175997 and SRR5176042 to SRR5176255) were previously determined and analyzed following the methods described in Martin-Platero et al, 2018. Coupled dynamics were analyzed by determining the Pearson coefficient of correlation for the relative abundance of each amplified sequence variant (ASV) of the *Saprospiraceae* and *Vibrionaceae* on each day of the Nahant time series taking into account time lagged correlations due to pred-ator-prey dynamic by smoothing the abundance data by a rolling average over 7 days. The threshold for significance of correlations was set to 0.03. The difference between the smoothed and direct correlation coefficient was determined and the 95^th^ percentile calculated. Every correlation above this threshold was stated as significant.

The DNA used for amplicon preparation, was used also for metagenomic sequencing. A subset of samples (Julian days 231, 235, 237, 239, 241, 243, and 247) were library prepped and sequenced by the BioMicroCenter (MIT, Cambridge, MA) on an Illumina NextSeq flowcell. Samples of each day were library prepped and sequenced by the BioMicroCenter (MIT, Cambridge, MA) using an Illumina NovaSeq S1flowcell.

The relative abundances of genomes were determined as followed: reads were i) cleaned of phiX with bowtie2 (v. 2.3.4.2) (Langmead & Salzberg, 2012) and quality trimmed with trimgalore (v.0.5.0) (https://github.com/FelixKrueger/TrimGalore) with the parameters “-e 0.05 --clip_R1 1 --clip_R2 1 --three_prime_clip_R1 1 --three_prime_clip_R2 1 --length 70 -- stringency 1 --paired --max_n 1 --phred33”; ii) error correction was done with Tadpole (v. 38.18, BBTools package https://jgi.doe.gov/data-and-tools/software-tools/bbtools/); iii) reads were filtered using kraken2 (v.2.1.3) (Wood *et al*, 2019a) classification (bacteria and archaea). In short, the kraken database was built with reference libraries of archaea and bacteria, kraken2 was run and reads were filtered with extract_kraken_reads.py (KrakenTools, https://github.com/jenniferlu717/KrakenTools; iv) Mapping was performed with bbmap (v.39.01) (BBTools package) to genomes with the parameters “fast=t minid=0.95 idfilter=0.98” and results sorted with samtools (Danecek *et al*, 2021).

For prey relative abundance, the reads were mapped to all available RefSeq genomes of the *Vibrionaceae* (218, NCBI, Nov. 2023) and summed. For the ixothrophy-positive strains, *Saprospira grandis* str. Lewin (GCF_000250635.1), *Aureispira* sp. CCB-QB1 (GCF_000724545.1), *Saprospira* sp. CCB-QB6 (GCF_028464065.1), *Aureispira* sp. CCB-E (GCF_031326345.1) and as control the ixotrophy-negative strain *Algoriphagus machipongonensis* (GCF_000166275.1) the rpkm (Reads Per Kilobase per Million mapped reads) values were calculated to account for different genome sizes. Plots were done with RStudio and smoothed by a rolling average of 3.

## Acknowledgements

ScopeM is acknowledged for instrument access at ETH Zürich. Functional Genomic Center Zurich is acknowledged for technical support. We thank Tobias Zachs for the help with cryoFIB-milling. We thank Florian Wollweber for help with light microscopy and segmentation of the cryo-tomograms. We thank Miroslav Peterek for technical support. We thank Pavel Afanasyev, Daniel Boehringer, Bilal Qureshi, and Dianhong Wang for their help with single particle cryoEM. We thank Jessie James Limlingan Malit for help with bioinformatics analysis. Charles Ericson is acknowledged for the help with sucrose gradient purification. We thank Lena Maria Leonie Keller for the help with qPCR. Wolf-Dietrich Hardt and Bidong Nguyen are acknowledged for insightful discussions on ISs. RS acknowledges support from a Gordon and Betty Moore Foundation Symbiosis in Aquatic Systems Initiative Investigator Award (GBMF9197; https://doi.org/10.37807/GBMF9197), a grant from the Simons Foundation (542395) as part of the Principles of Microbial Ecosystems (PriME) Collaborative, a grant (315230_176189) from the Swiss National Science Foundation and support from the National Centre of Competence in Research (NCCR) Microbiomes (51NF40_180575). MP was supported by the Swiss National Science Foundation (310030_212592), the European Research Council (679209 and 101000232) and the NOMIS foundation. NB and MFP were supported by the Simons Foundation (Life Sciences Project Award-572792).

## Author contributions

YWL conducted all experiments that include light microscopy, killing assays, cryoFIB milling, cryoET, qRT-PCR and characterization of IS excision. GLW generated preliminary data, supported cryoFIB, cryoET and sub-tomogram averaging. With support from YWL and JX, DA established the purification of grappling hooks, processed the cryoEM data, built an initial atomic model and analyzed the structures. Raman microspectroscopy and data analysis were performed by KSL in the lab of RS. NB and MFP analyzed the time-resolved metagenomic dataset. GF shared the initial *Aureispira* culture and gave feedback during discussions. YWL, GLW and MP wrote the manuscript with comments from all authors.

## Data and material availability

Example tomograms (EMD-XXXX–EMD-XXXX), subtomogram averages (EMD-XXXX– EMD-XXXX) and cryoEM maps (EMD-XXXX–EMD-XXX) will be uploaded to the Electron Microscopy Data Bank. Atomic coordinates of the grappling hook models will be uploaded to the Protein Data Bank. Source data are provided with this paper. Sequenced genomes will be uploaded to XXXX.

---

**Figure S1.**
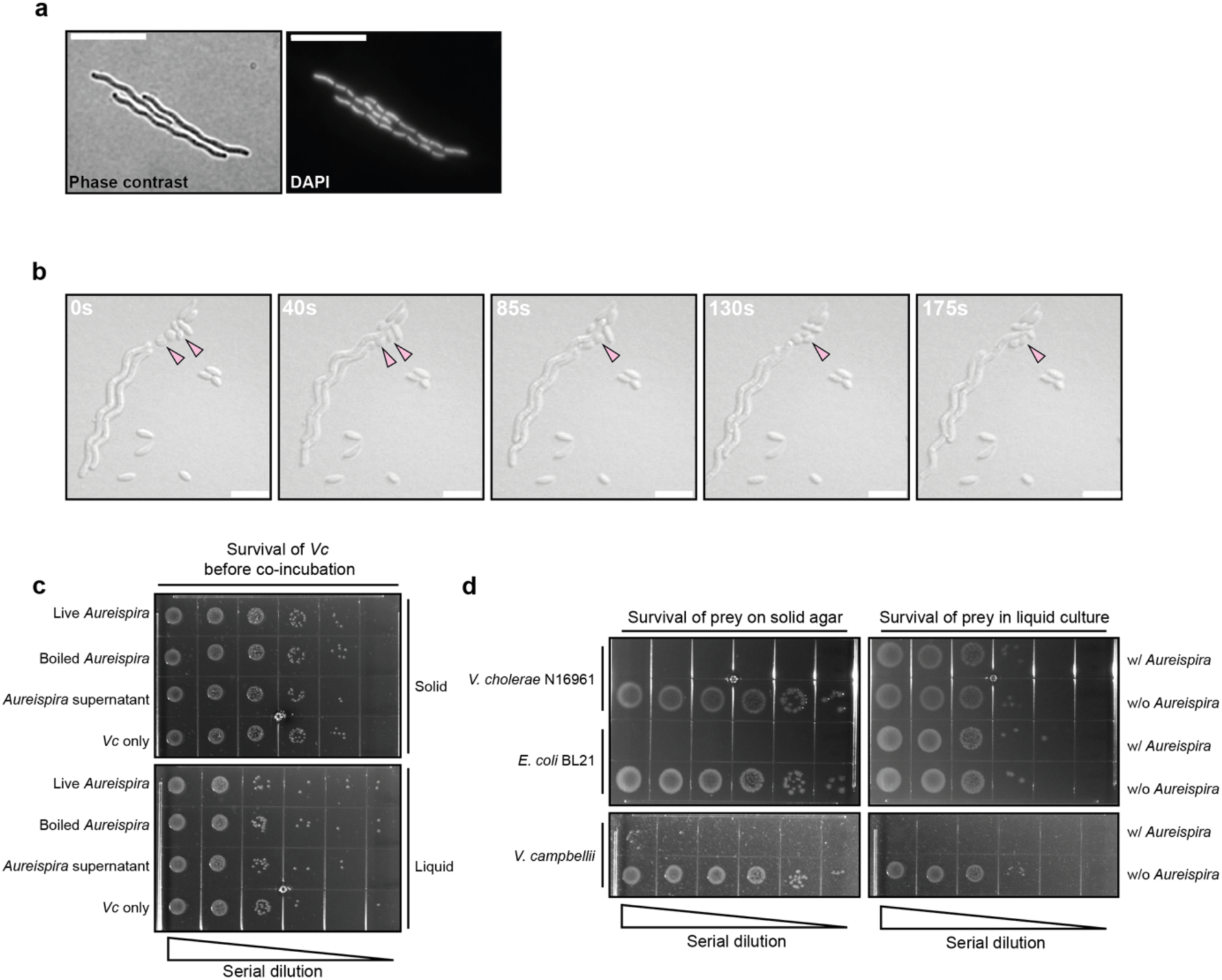
Characterization of *Aureispira* ixotrophy behavior. **a:** Phase contrast (left) and fluorescent (right) light microscopy of DAPI stained *Aureispira* cells show multicellularity of *Aureispira* filaments. Bar: 10 μm. **b:** Another example of time-lapse LM of *Aureispira* incubated with *Vc*. In the case shown here, *Aureispira* glided “head-on” towards *Vc*. After contact with *Aureispira* filaments, *Vc* cells (arrowheads) showed signs of rounding, and subsequently lysed. Bar: 10 μm. **c:** Quantification of *Vc* prey cells prior to co-incubation killing assay with differently treated *Aureispira* cultures (corresponds to Fig. 1c). The cultures were serially diluted, dropped onto agar plates, and incubated to verify identical *Vc* cell concentrations at the start of the experiment. d: Quantification of the survival diverse prey cells after co-incubation with *Aureispira* on solid agar or in liquid culture. The cultures were serially diluted, dropped onto agar plates, and incubated to verify cell concentrations.

**Figure S2.**
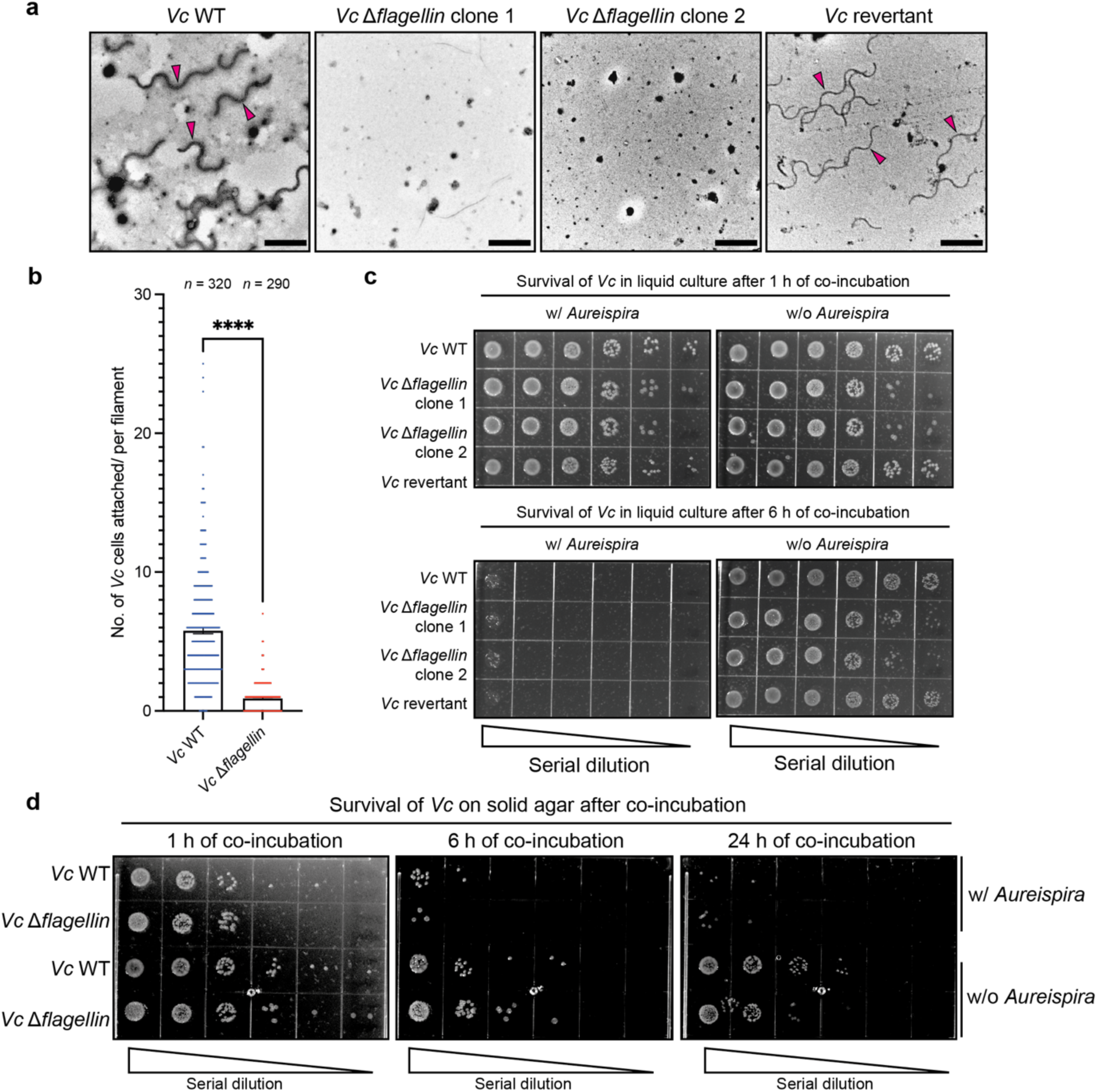
Prey flagella are involved in ixotrophy killing in liquid culture. **a:** Negative-stain electron micrographs of purified flagella from *Vc* WT and mutants. While flagella (arrowheads) were observed in both WT and revertant, no flagella were detected in the Δ*flagellin* mutants. Bar: 2 μm. **b:** Quantification of *Vc* cells attached to the surface of an individual *Aureispira* filament after co-incubation in liquid culture. The analysis of LM images showed a significant (p ≤ 0.0001) reduction of prey cell attachment to *Aureispira* when incubated with *Vc* Δ*flagellin* mutant compared to WT. Un-paired t-test was used for the statistical analysis. The sample sizes (*n*) are indicated at the top of the chart. **c:** Serial dilution drop assay of *Vc* derivatives after co-incubation with *Aureispira* in liquid culture showed that the *Vc* Δ*flagellin* mutants are still killed by *Aureispira,* but less efficient compared to flagellated *Vc* WT (see Fig. 2c for the data corresponding to co-incubation for 3 h). **d:** Serial dilution drop assay of *Vc* derivatives after co-incubation with *Aureispira* on solid media revealed no difference in killing rate between WT and *Vc* Δ*flagellin* mutant.

**Figure S3.**
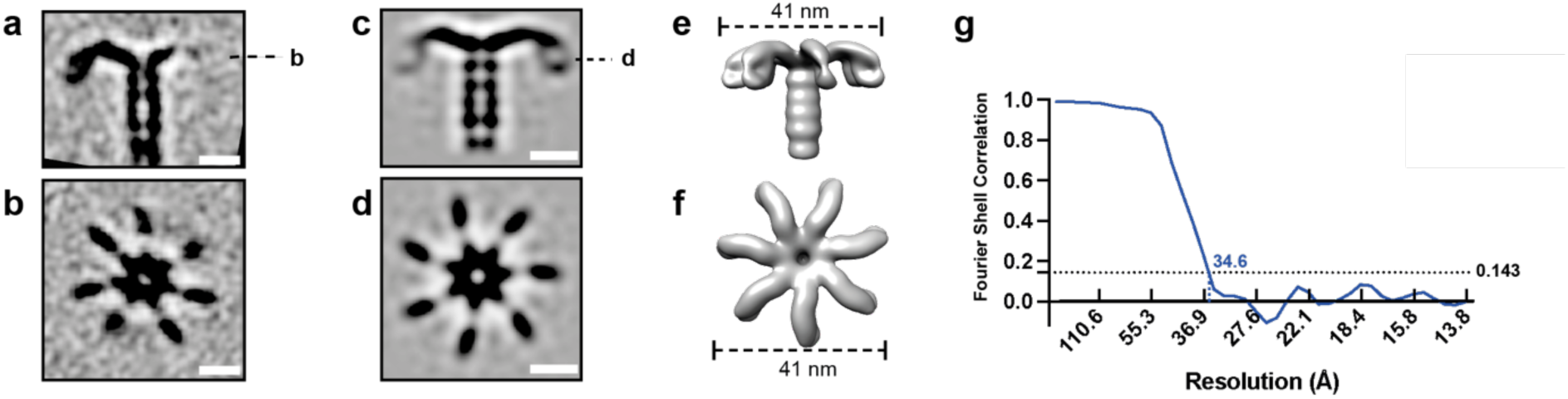
*In situ* subtomogram averaging of the grappling hook distal end. **a/b:** Shown are longitudinal (**a**) and perpendicular (**b**) sections through the unsymmetrized subtomogram average of the *Aureispira* grappling hook distal end, which consists of seven hook-like densities. Bar: 10 nm. **c/d:** Shown are longitudinal (**c**) and perpendicular (**d**) sections through the C7-symmetrized subtomogram average of the distal end of grappling hooks. Bar: 10 nm. **e/f:** Isosurface representation of the subtomogram average shown in **c/d**. **g:** Fourier Shell Correlation (FSC) analysis of the C7-symmetrized sub-tomogram average of grappling hook distal ends shown in **c-f**. The approximate resolution was 34.6 Å.

**Figure S4.**
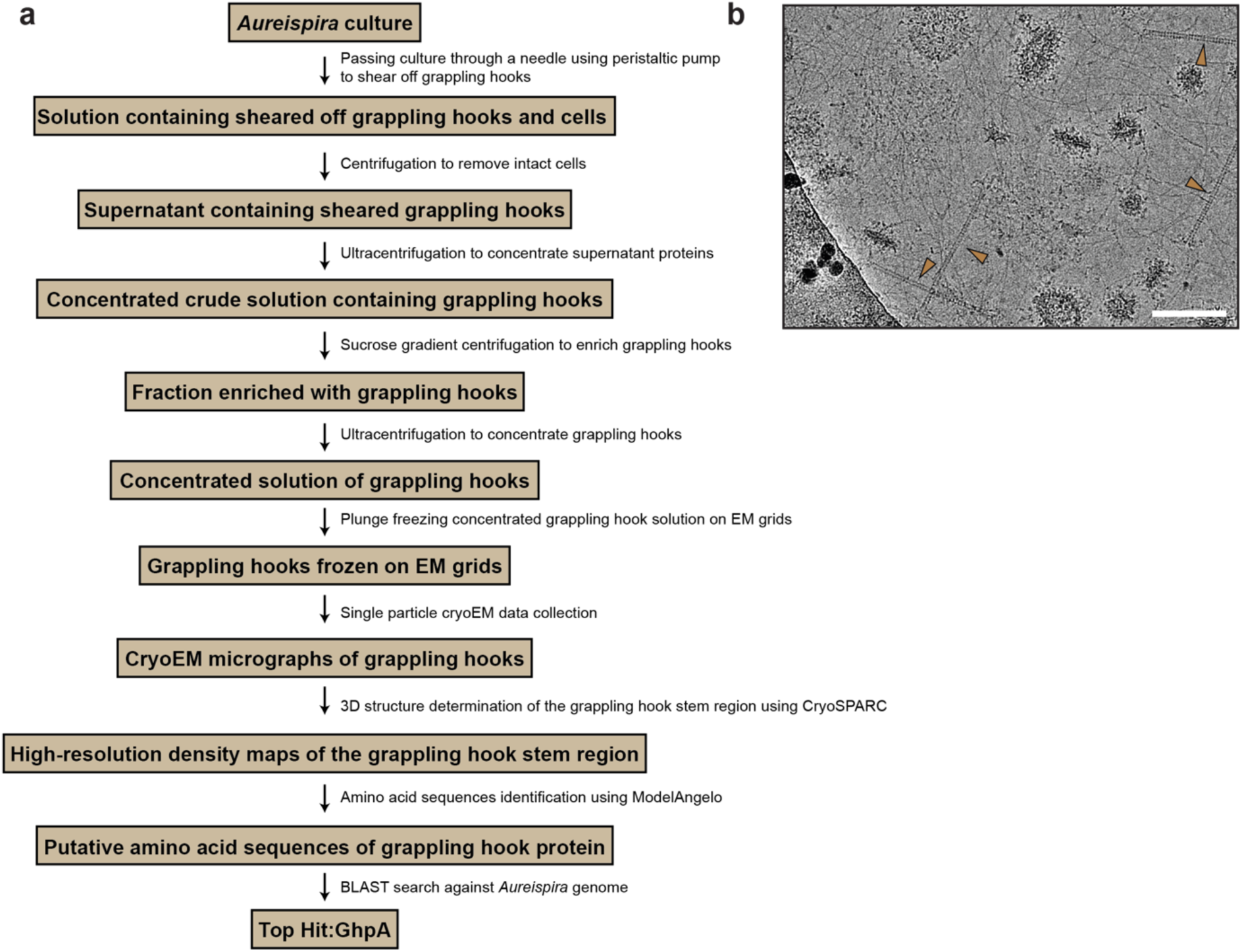
Schematic of workflow for grappling hook purification and protein identification via single particle cryoEM. **a:** *Aureispira* cells were harvested, and extracellular appendages were sheared off by passing the cell suspension through a needle using a peristaltic pump. After pelleting intact cells and cellular debris, the supernatant was subjected to ultracentrifugation to concentrate the sheared-off grappling hooks. The concentrated crude solution was then further purified using sucrose density gradient ultracentrifugation. The fraction containing grappling hooks was again concentrated by ultracentrifugation, applied to an EM grid and subsequently plunge-frozen. The frozen grids were transferred to a cryo-transmission electron microscope for single particle cryoEM data acquisition. The cryoEM data was processed using the software CryoSPARC to obtain high-resolution density maps of the grappling hook stem region. The resulting maps were used to identify the corresponding amino acid sequence by automatic atomic model building using the software ModelAngelo. The suggested amino acid sequences were subjected to a BLAST search against the *Aureispira* genome to identify candidate proteins. GhpA was identified as top hit for the grappling hook protein. A detailed description of the entire workflow can be found in Materials and Methods. **b:** Representative motion-corrected cryoEM 2D micrograph of purified grappling hooks (arrowheads). Bar: 100 nm.

**Figure S5.**
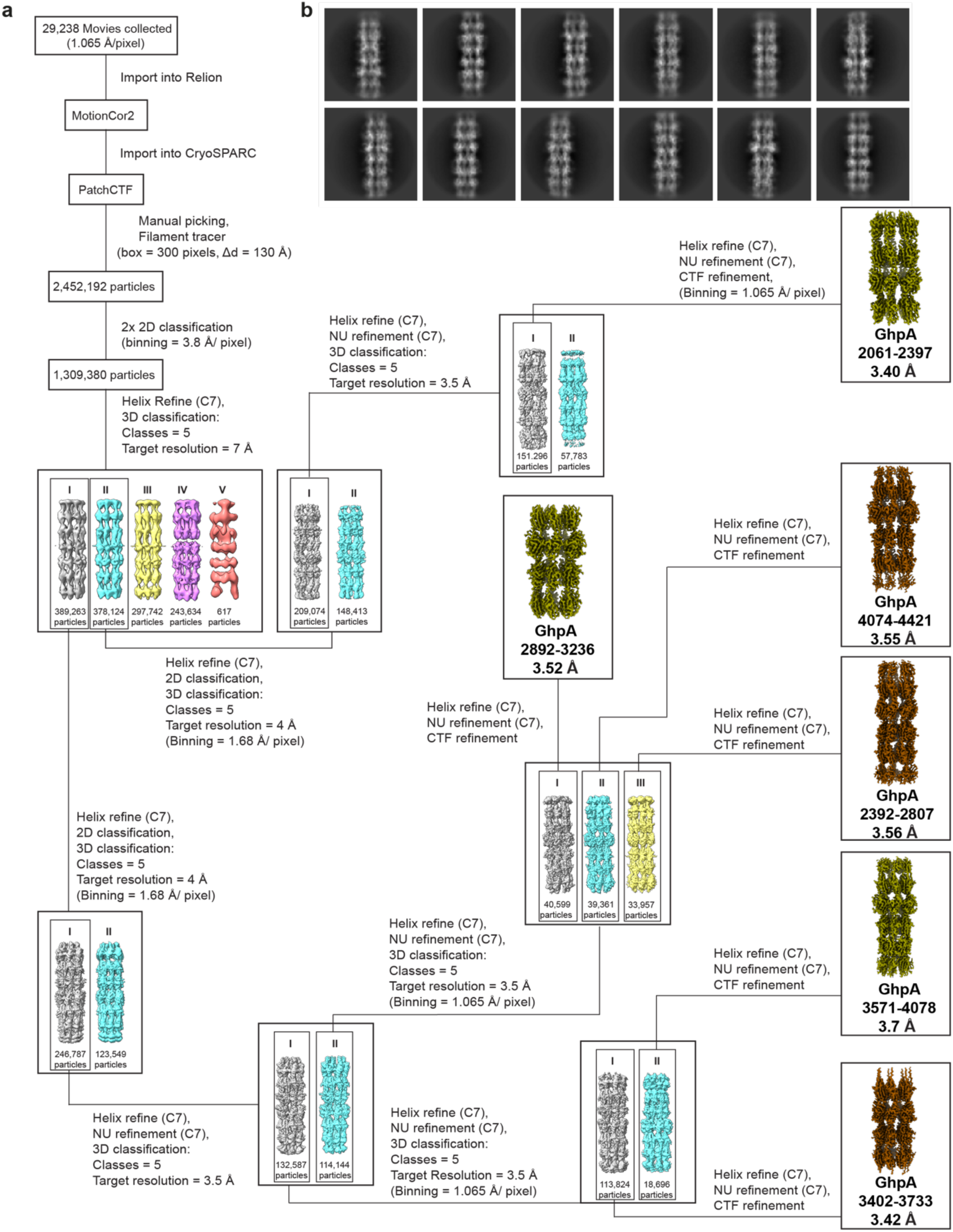
Schematic representation of cryoEM data processing for the six structures obtained of the grappling hook stem region. **a:** Schematic of the data processing workflow used to obtain cryoEM density maps of six fragments of the grappling hook stem region. All steps except for the initial motion-correction were carried out in CryoSPARC (Punjani *et al*, 2017). Key parameters of the processing steps are stated next to the corresponding step. Only 3D classes with more than 100 particles are shown. Δd: inter-box distance, NU Refinement: non-uniform refinement (Punjani *et al*, 2020). A detailed description of the workflow can be found in the Materials and Methods section. **b:** Representative 2D classes of the grappling hook stem. The individual Ig-like domains and the hollow lumen are visible.

**Figure S6.**
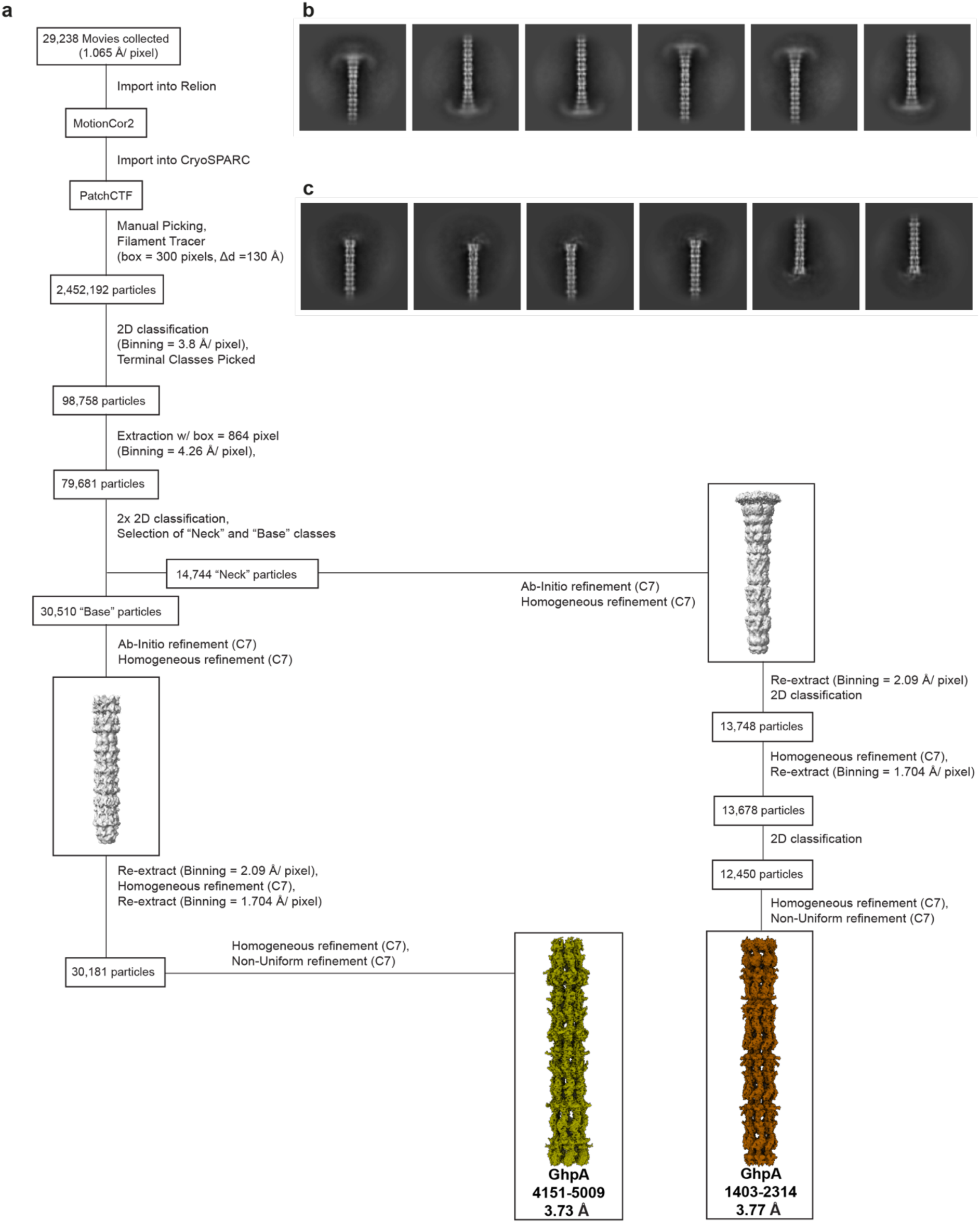
Schematic representation of cryoEM data processing workflow for the grappling hook neck and base structures. **a:** Schematic of the data processing workflow used to obtain cryoEM density maps of the grappling hook neck (GhpA 1403-2314) and base region (GhpA 4151-5009). Key parameters of the processing steps are stated next to the corresponding step. A more detailed description can be found in Materials and Methods section. **b:** Representative 2D classes of the grappling hook neck (GhpA 1403-2314). The hook structures are smeared out due to flexibility. **c:** Representative 2D classes of the grappling hook base (GhpA 4151-5009). The densities corresponding to the proximal region of the base were smeared out, presumably due to the fact that grappling hooks were sheared off during purification, inducing structural flexibility in this region.

**Figure S7.**
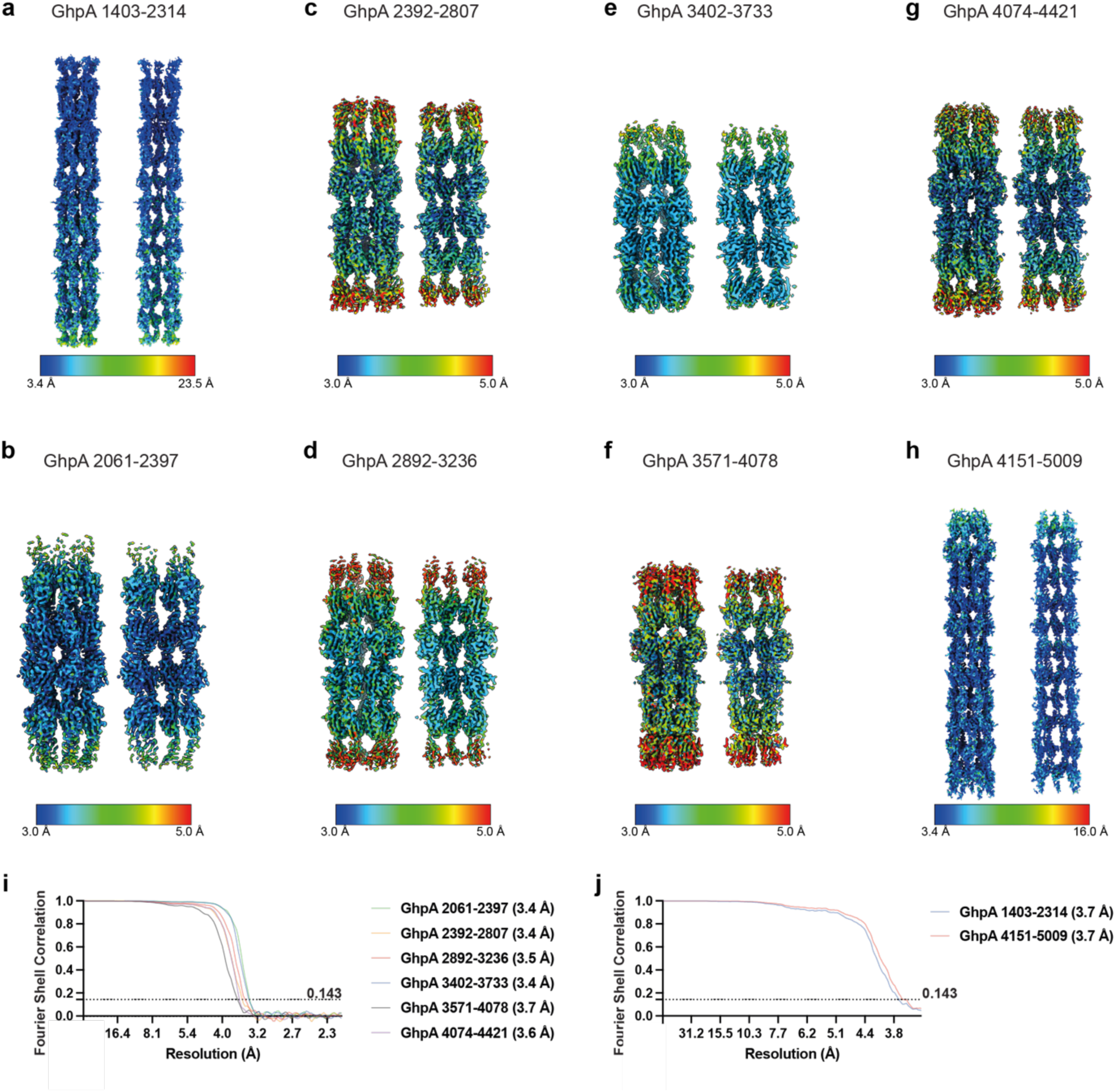
Local resolution estimation and Fourier shell correlation curves of the density maps obtained for the grappling hook stem region. **a:** CryoEM density map of grappling hook neck region with indicated local resolution. The coloring corresponds to resolutions between 3.4 Å (blue) and 23.5 Å (red). **b-g:** CryoEM density maps of fragments of grappling hook stem region with indicated local resolutions. The coloring corresponds to resolutions between 3.0 Å (blue) and 5.0 Å (red). **h:** CryoEM density map of grappling hook base region with indicated local resolution. The coloring corresponds to resolutions between 3.4 Å (blue) and 16.0 Å (red). **i:** Fourier shell correlations (FSCs) of two half maps obtained for the individual fragments of the grappling hook stem module calculated with a tight mask. The estimated “gold-standard FSC” resolution at a threshold of 0.143 is indicated on the right. **j:** Fourier shell correlations (FSCs) of two half maps obtained for the grappling hook base and neck modules calculated with a tight mask and corrected by noise substitution. The estimated “gold-standard FSC” resolution at a threshold of 0.143 is indicated on the right.

**Figure S8.**
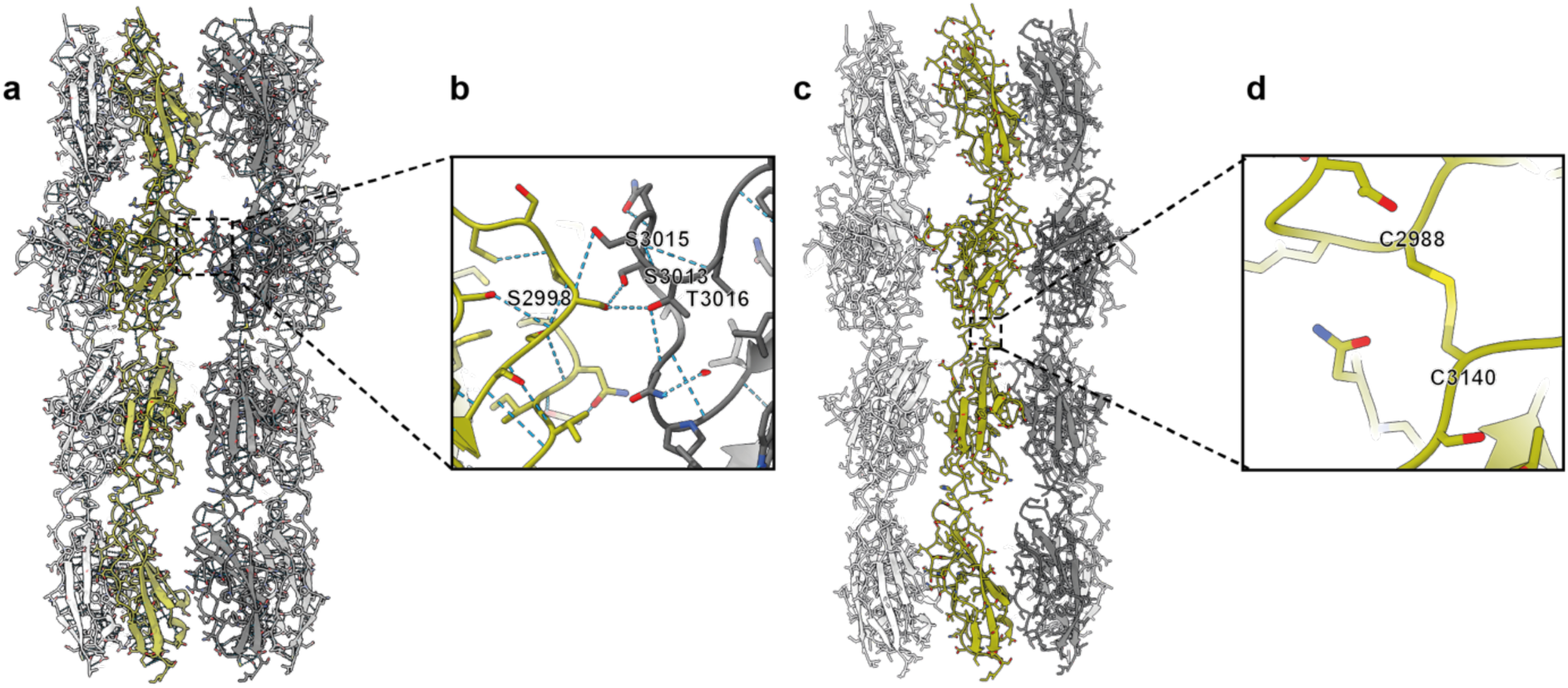
Atomic model of GhpA reveals intra- and inter-protomer interactions. **a/c:** Different orientations of atomic model of GhpA stem region (residue 2,892-3,236) shown in ribbon representation. **b:** Hydrogen bonds are found to facilitate interactions between each GhpA protomers. Shown is the region corresponding to the dashed box in **a**. The structure is visualized with yellow or gray for the peptide backbone of two different GhpA protomers, respectively: red for oxygen atoms, blue for nitrogen atoms and light blue for the hydrogen bonds (dashed lines). **d:** Disulfide bonds are found between pairs of nearby Ig-like domains. Shown is the disulfide bond between residue Cys-2,988 and Cys-3,140. The structure is visualized with green for the peptide backbone, red for oxygen atoms, blue for nitrogen atoms and yellow for the disulfide bond. The region shown corresponds to the dashed box in **c**.

**Figure S9.**
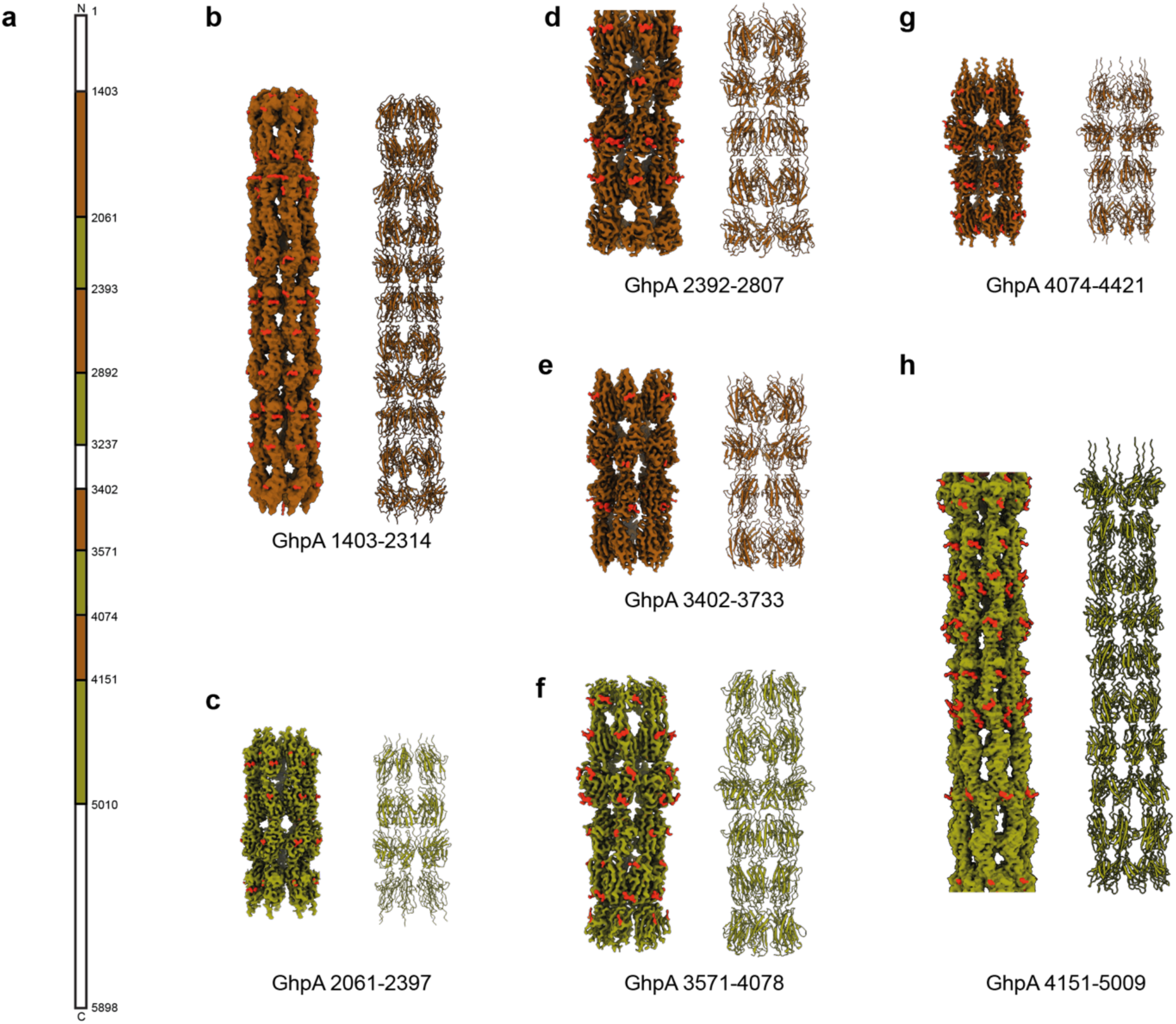
Potential glycans are found in all cryoEM density maps along the grappling hook stem. **a:** Schematic representation of GhpA protein. Indicated fragments (brown and green) correspond to individual single-particle cryoEM density maps. Amino acid residues which have been built in the corresponding atomic models are indicated on the right. **b-h:** CryoEM density maps (left) with corresponding atomic models (right) of different regions along the grappling hook stem. The cryoEM density map was masked with a 3 Å zone around the atomic model. This masking was used to color all densities, which are more than 3 Å distant from the center axis of the atomic model in red. These protrusions potentially correspond to O-glycosylation sites.

**Figure S10.**
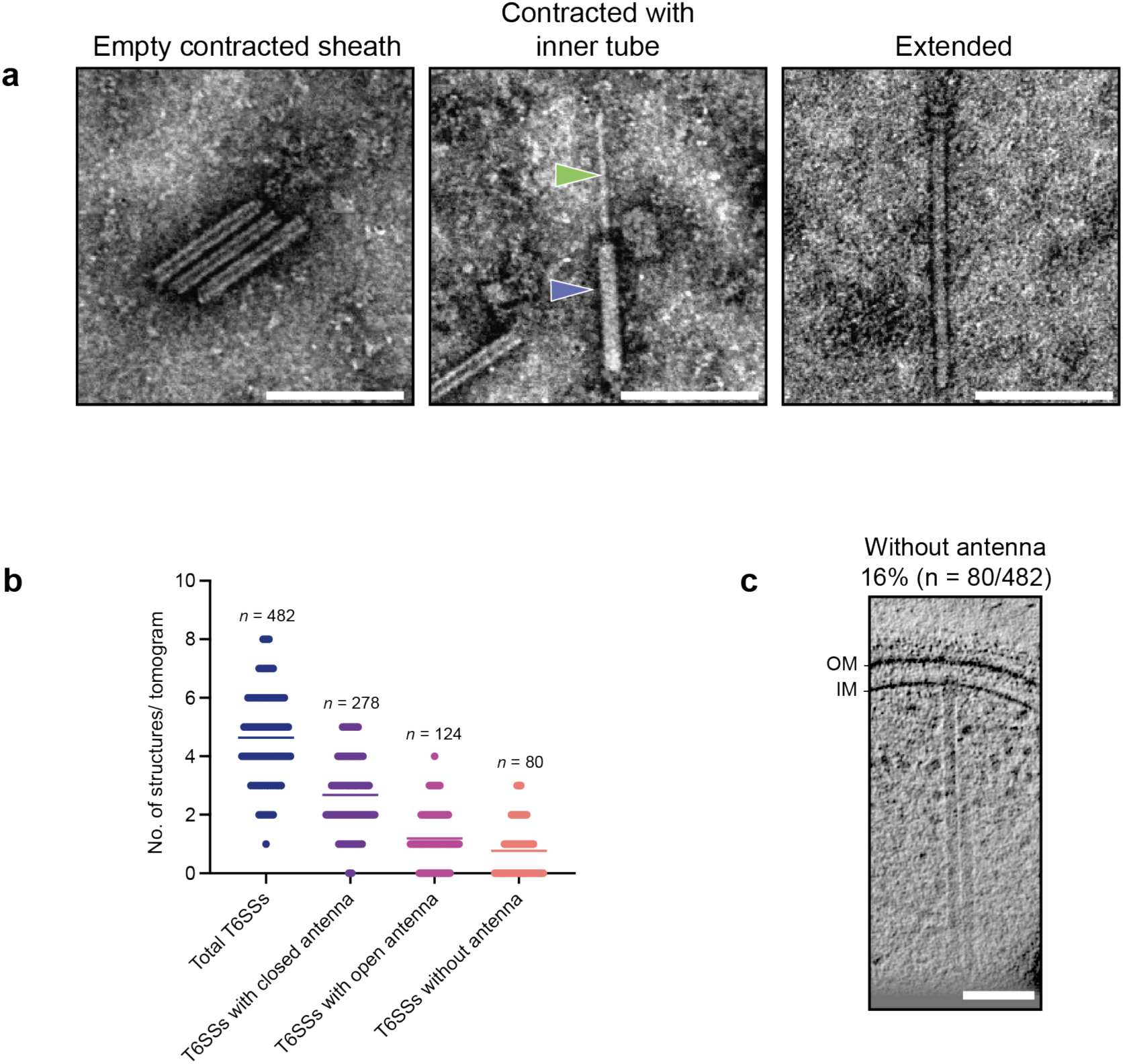
Characterization of *Aureispira* T6SS. **a:** Negative stain electron micrographs of empty contracted sheaths (left), contracted sheath (blue arrowhead) with attached inner tube (green arrowhead; center) and extended T6SS (right), which were purified from *Aureispira* cultures. Bar: 200 nm. **b:** Quantification of the states of extracellular antennae connected to *Aureispira* T6SSs as observed in cryo-tomograms. In total, 104 tomograms were analyzed. Thin lines indicate the average number. **c:** Slice through cryo-tomogram showing an *Aureispira* T6SS without extracellular antenna, which was observed in 16% of detected T6SSs. Bar: 100 nm. Thickness of the slice: 13.8 nm.

**Figure S11.**
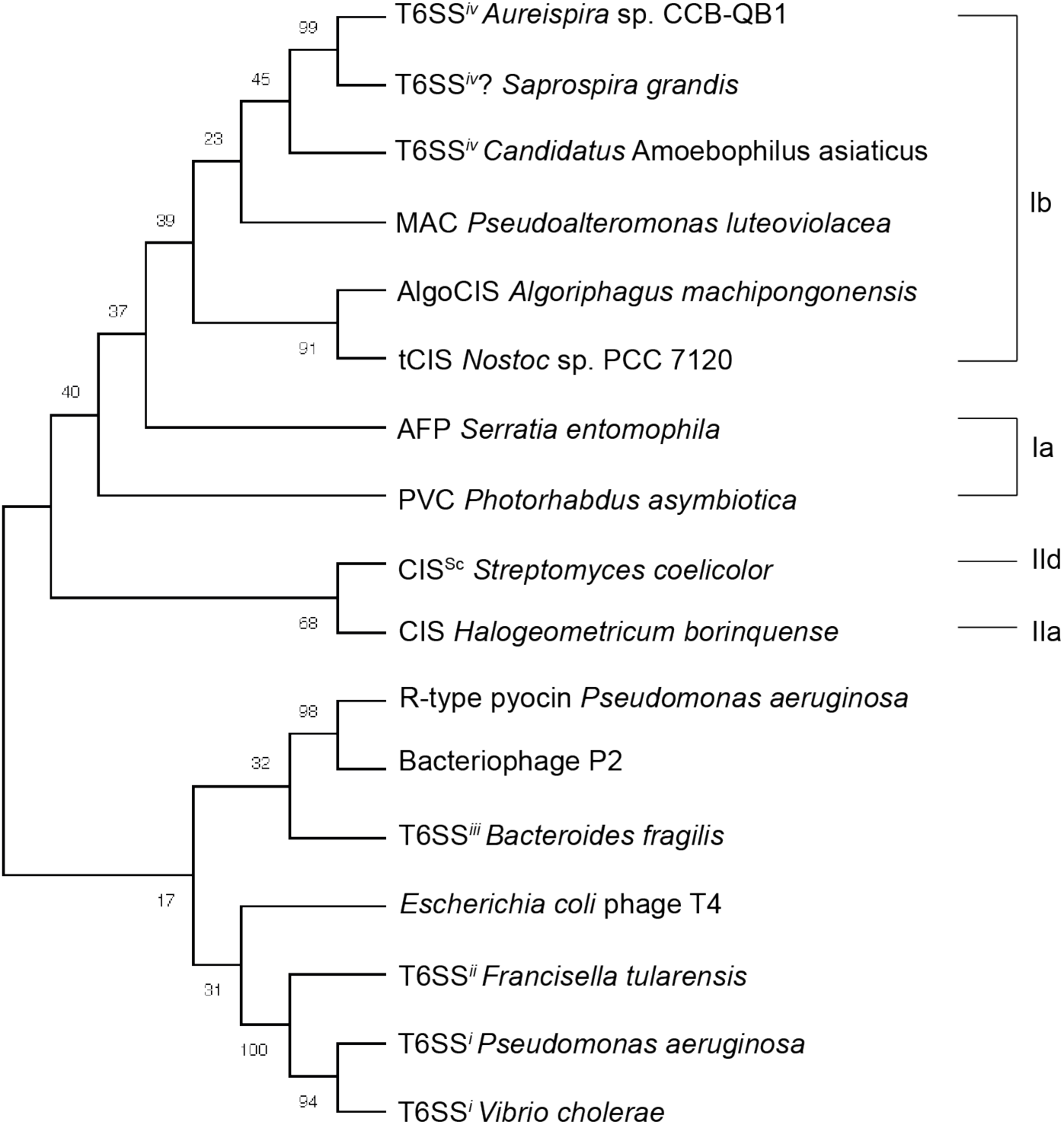
Phylogenetic analysis indicates that T6SS*^iv^* sheath proteins form a monophy-letic group. Sheath protein (Cis2) from CISs from different organisms were used to calculate the phylogenetic tree. The classification of CIS according to Chen et al. (Chen *et al*, 2019) is indicated on the right. The bootstrap values are shown on the branches. *Aureispira* Cis2 clusters with Cis2 from *Saprospira grandis*, which is another ixotrophy-positive strain in the family *Saprospiraceae* (Lewin, 1997), harboring a CIS gene cluster.

**Figure S12.**
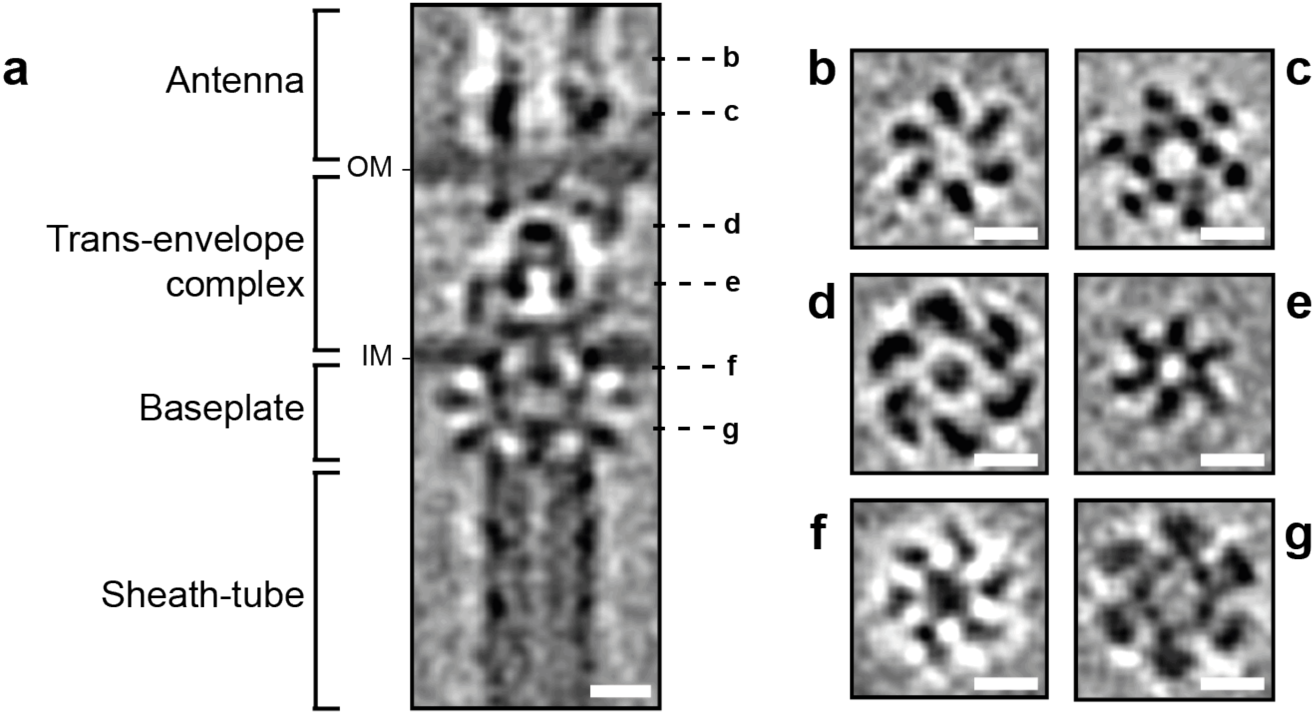
*In situ* sub-tomogram average of *Aureispira* T6SSs revealed C6-symmetry of the extracellular antenna, the trans-envelope complex and the baseplate. **a:** Longitudinal slice through the unsymmetrized subtomogram average of *Aureispira* T6SSs, covering the base of the extracellular antenna, the trans-envelope complex, the IM-anchoring baseplate as well as the sheath-tube module. The position of perpendicular slices (**b-g**) is indicated on the right. Bar: 10 nm. **b/c:** Perpendicular slices through the unsymmetrized average of the base of the extracellular antenna. Bar: 10 nm. **d/e:** Perpendicular slices through the unsymmetrized average of the trans-envelope complex. Bar: 10 nm. **f/g:** Perpendicular slices through the unsymmetrized average of the baseplate. Bar: 10 nm.

**Figure S13.**
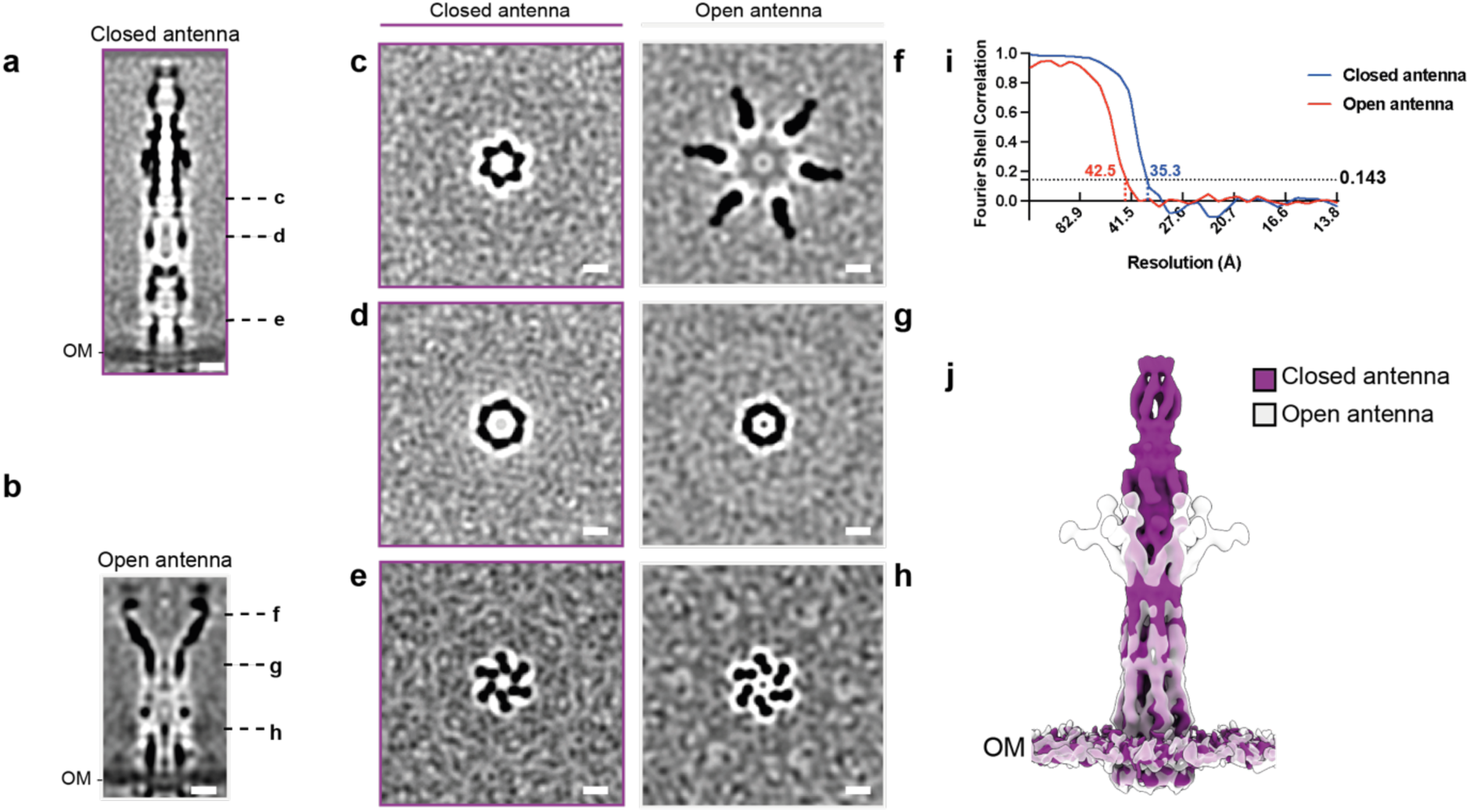
*In situ* subtomogram averages of antennae in open and closed conformations. a/b: Longitudinal slices through C6-symmetrized subtomogram averages of antennae in closed. (**a**) and opened (**b**) conformations. Positions of perpendicular slices (**c-h**) are indicated on the right. Shown are 0.7 nm thick slices. Bar: 10 nm. **c-e:** Perpendicular slices through the C6-symmetrized average of closed antennae. Bar: 10 nm. **f-h:** Perpendicular slices through the C6-symmetrized average of open antennae. Bar: 10 nm. **i:** FSC of two half-maps from sub-tomogram averaging of the antennae in both conformations. The approximate resolutions (FSC: 0.143) were 35.3 Å and 42.5 Å for the closed and open antennae, respectively. **j:** Isosurface representation of the density map of open antennae fitted into the density map of closed antennae. While the open antenna (grey) widened significantly at half of its length compared to the antenna in the closed conformation (purple), the OM-connected bases of both conformations appeared to be similar.

**Figure S14.**
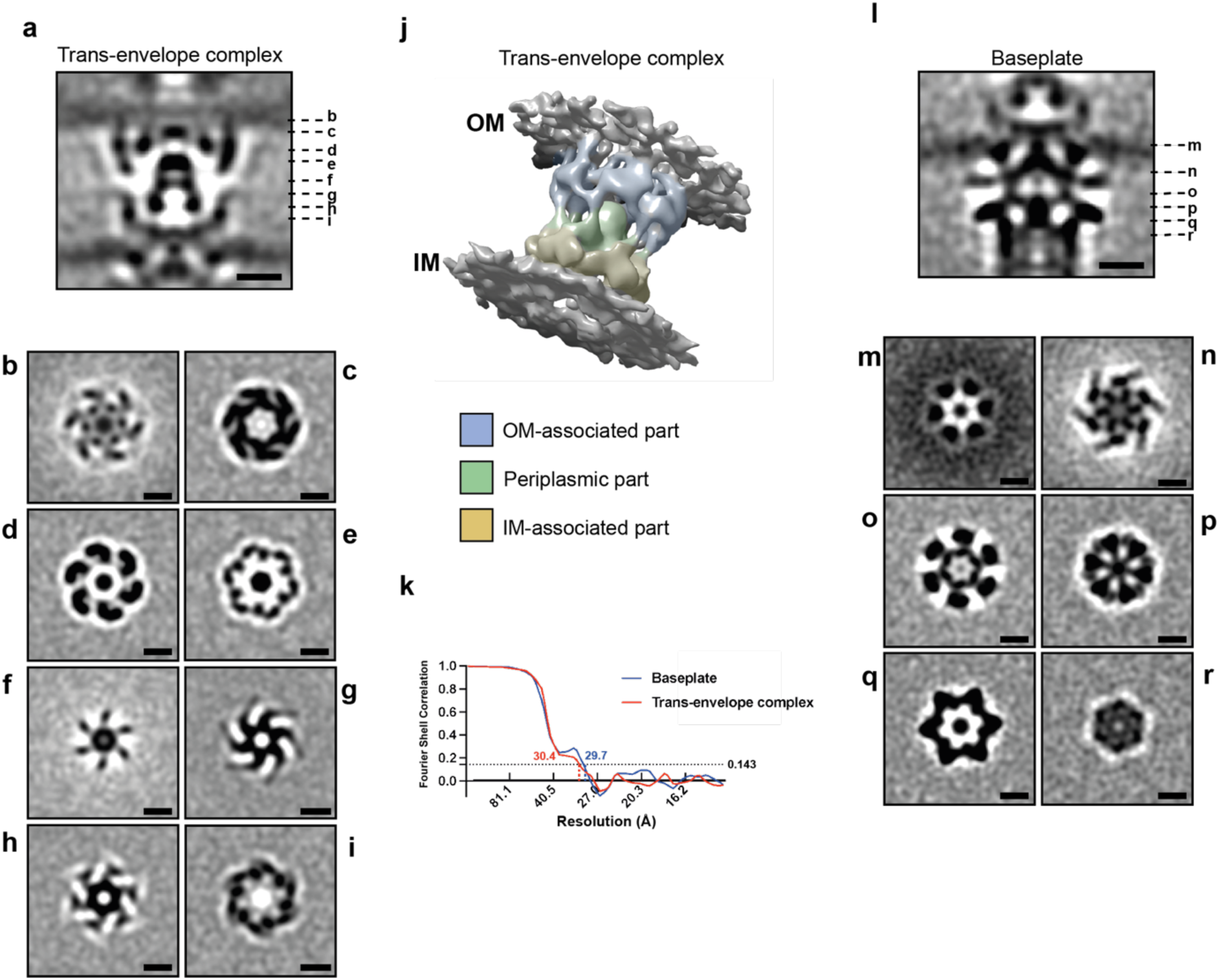
*In situ* subtomogram averages of the *Aureispira* T6SS trans-envelope complex and baseplate. **a-i:** Longitudinal (**a**) and perpendicular (**b-i**) slices through C6-symmetrized subtomogram average of trans-envelope complex. Positions of perpendicular slices (**b-i**) are indicated on the right in **a**. Shown are 0.7 nm thick slices. Bar: 10 nm. **j:** Isosurface representation of subtomogram average of trans-envelope complex. The density map was colored to indicate the three main modules: an outer membrane-associated domain, a periplasmic domain and an inner membrane-associated domain (yellow). **k:** FSC of two half-maps of subtomogram averages of the baseplate and the trans-envelope complex. The approximate resolution (FSC: 0.143) was 29.7 Å for the baseplate and 30.4 Å for the trans-envelope complex. **l-r:** Longitudinal (**l**) and perpendicular (**m-r**) slices through C6-symmetrized subtomogram average of baseplate region. Positions of perpendicular slices (**m-r**) are indicated on the right in **l**. Shown are 0.7 nm thick slices. Bar: 10 nm.

**Figure S15.**
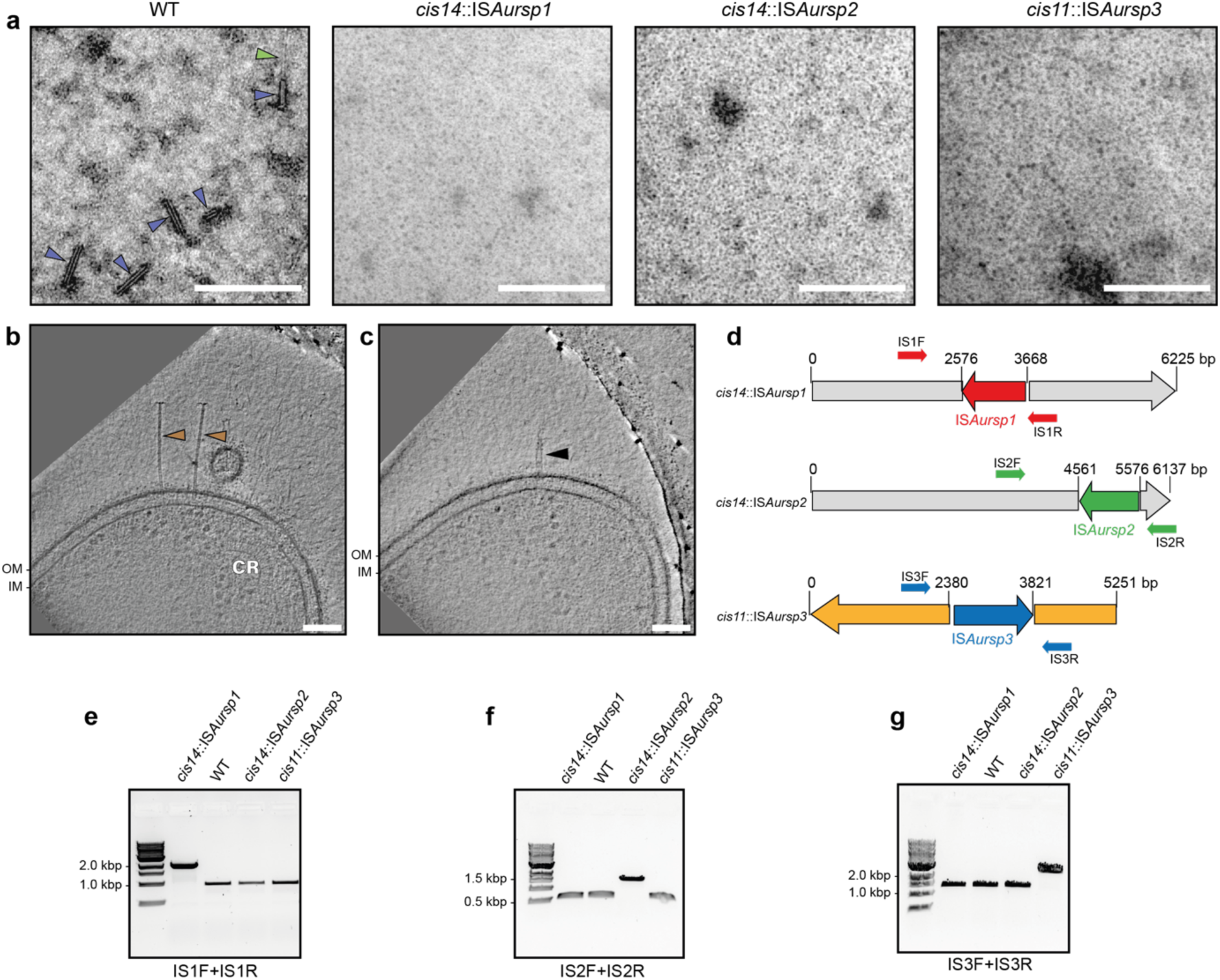
Characterization of IS mutants. **a:** Negative-stain electron micrographs of crude sheath purification confirm the absence of T6SSs in the IS mutants. Blue arrowhead: contracted sheaths. Green arrowhead: inner tube. Bar: 500 nm. **b/c:** Cryo-tomograms of the T6SS-negative *Aureispira* derivative *cis14*::IS*Aursp1*, showing the absence of T6SSs, while grappling hooks and antennae were still present. Brown arrowheads: grappling hooks. Black arrowhead: closed antenna. Bar: 100 nm. Thickness of the slices: 13.8 nm. **d:** Schematic representation of the location of ISs in *cis14* and *cis11*. The base pairs corresponding to the IS insertion sites are indicated above. The primers used in **e-g** are indicated as arrows flanking the respective IS. **e-g:** PCR confirmed IS insertion in the T6SS-negative mutants. The genomic DNA of each T6SS-negative mutant and WT was isolated and subjected to PCR. Used primer pairs are shown below and the binding sites of the primers are indicated schematically in **d**.

**Figure S16.**
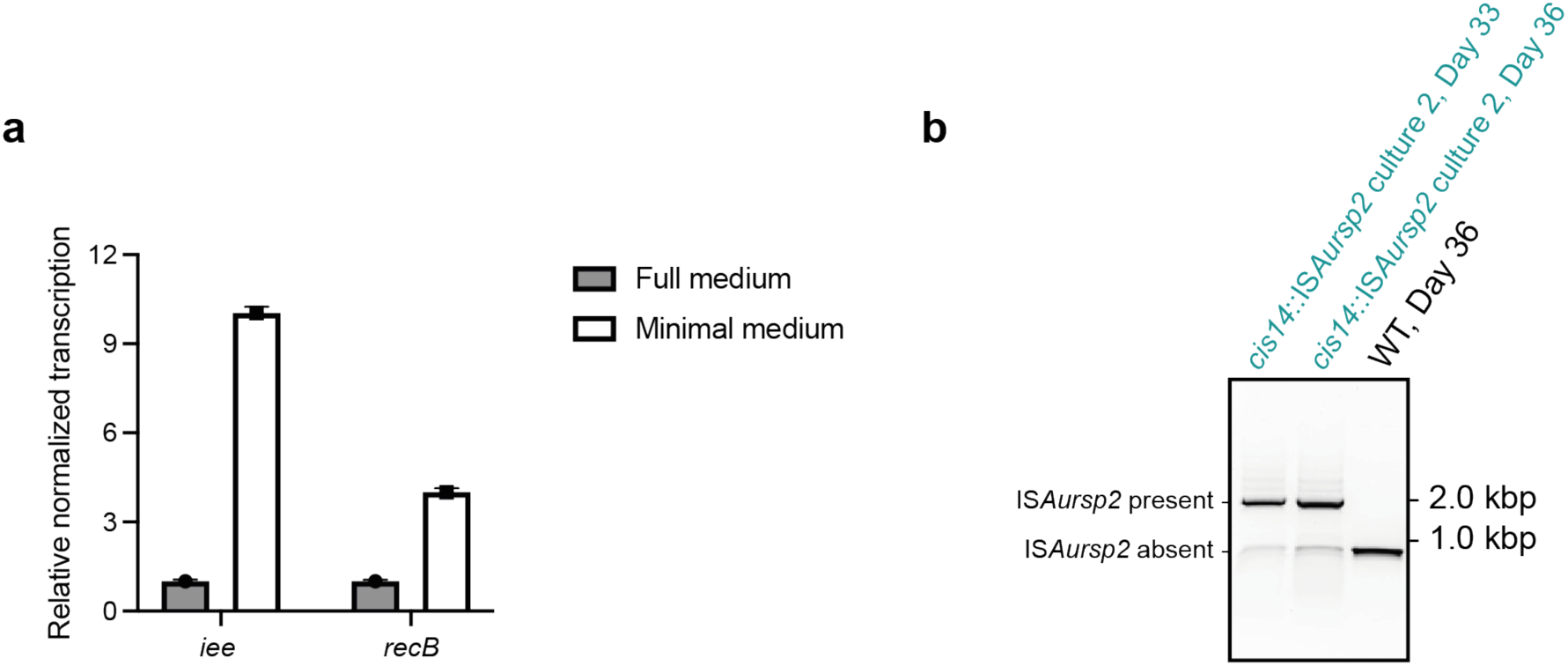
Further characterization of IS excision during starvation. **a:** Quantitative reverse transcription polymerase chain reaction (qRT-PCR) shows upregulation of genes involved in IS excision under starvation. Total RNA from *Aureispira* cultured in minimal medium or in rich medium was extracted, and qRT-PCR was used to determine the relative normalized transcription of insertion sequence excision enhancer (*iee*) and *recB*. The expression level was normalized with the transcript of *rpoB* (a gene involved in transcription). **b:** PCR confirms another example of IS excision events in *cis14*::IS*Aursp2* culture during starvation. *Aureispira-Vc* culture under starvation was boiled to release DNA, and the total DNA was used for PCR. Binding sites of the primers (IS2F and IS2R) are indicated schematically in **Fig. S15d**.

**Figure S17.**
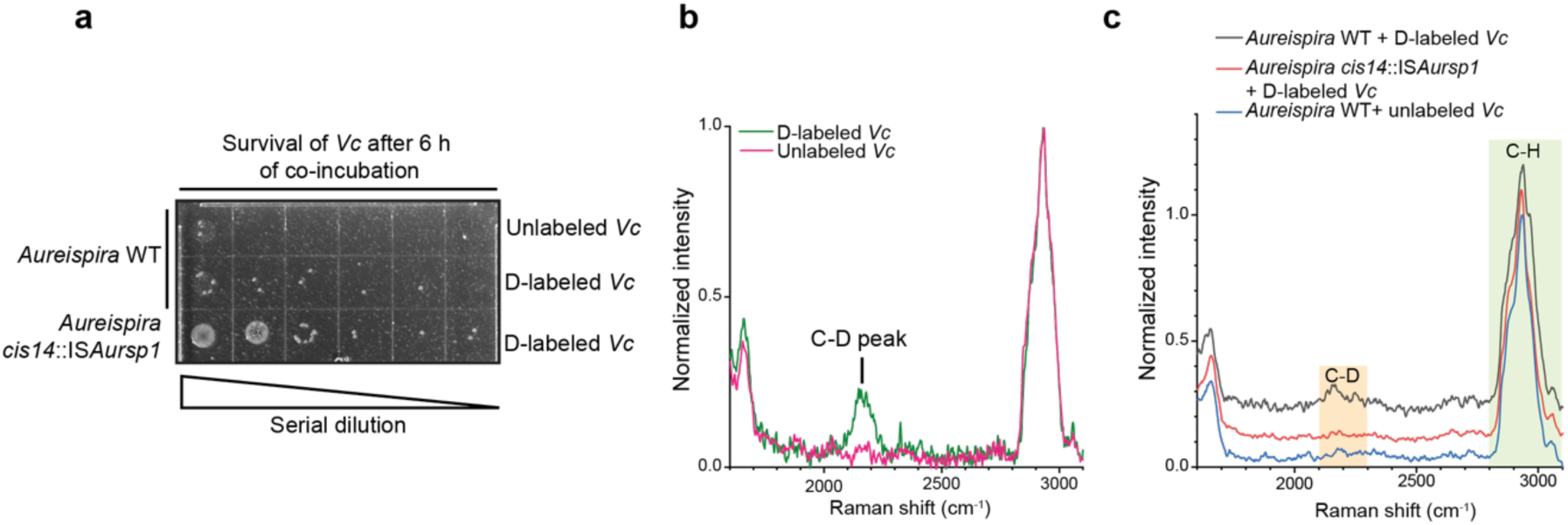
Raman microspectroscopy reveals that prey cell components are taken up by *Aureispira*. **a:** Deuterium (D)–labeled *Vc* cells were killed by *Aureispira* as efficiently as unlabeled *Vc*. *Aureispira* was co-cultured with either D–labeled or unlabeled *Vc* for 6 h and the survival of *Vc* was analyed by serial dilution and dropping on an agar plate. **b:** *Vc* cells were successfully labeled with deuterium. Representative Raman spectra of D– labeled and non–labeled *Vc* cells. The spectra were normalized using “(Raw intensity - minimum intensity within the spectral window) / (maximum intensity - minimum intensity)”. **c:** Representative Raman spectra of *Aureispira* cells under three different conditions: *Aureispira* WT co-incubated with D–labeled *Vc* (black), *Aureispira cis14*::IS*Aursp1* co-incubated with D–labeled *Vc* (red), and *Aureispira* WT co-incubated with unlabeled *Vc* (blue). The spectra were normalized and, for visual clarity, all spectra are offset on the y-axis (by 0.1).

**Figure S18.**
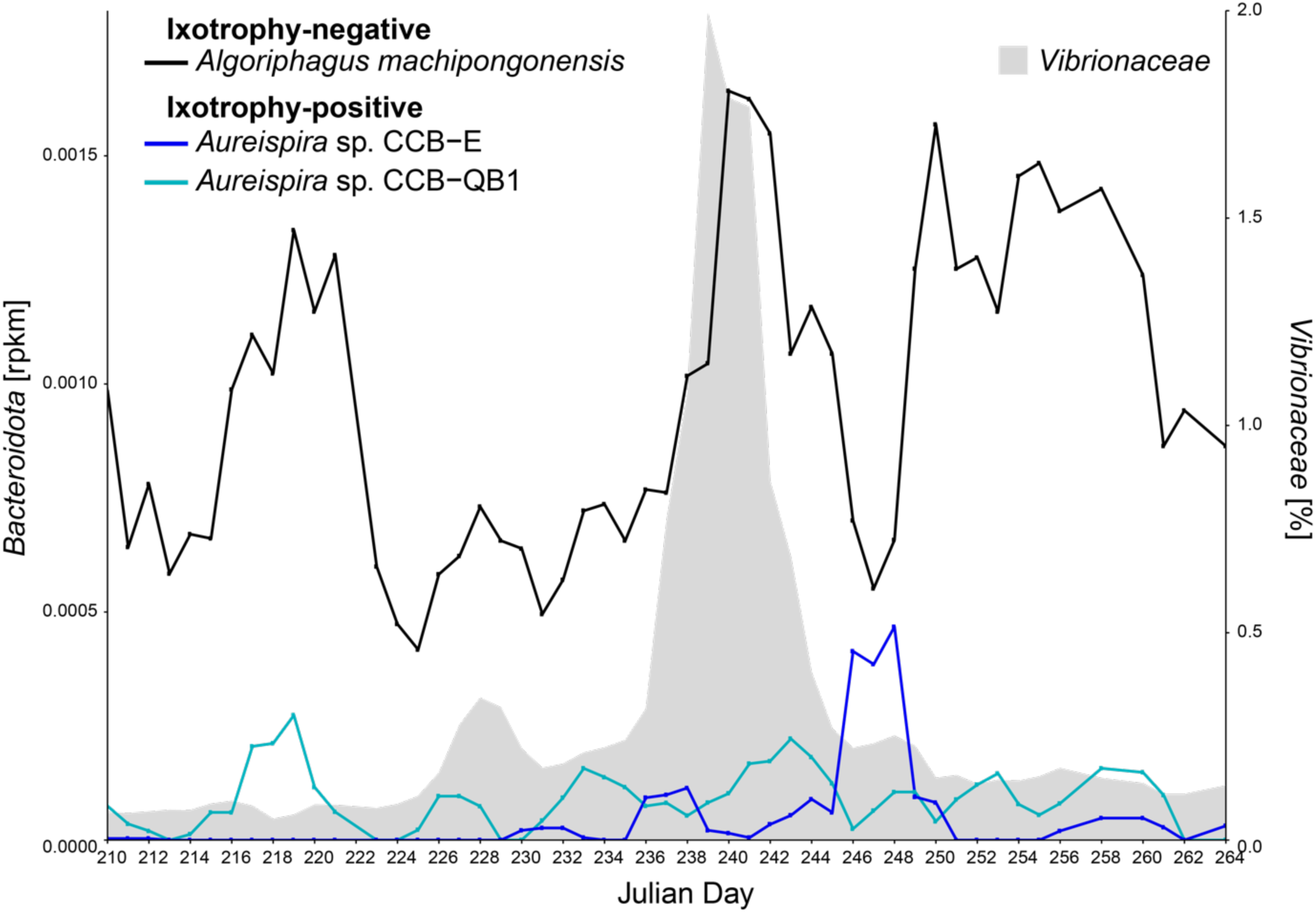
Predator-prey dynamic between *Aureispira* and *Vibrionaceae* in the ocean. Analysis of a time-resolved metagenomic dataset (Nahant time series) shows correlation between the abundance of *Vibrionaceae* (gray) and *Aureispira* species (cyan and blue). The correlation, however, is less pronounced compare to the analyzed *Saprospira* strains (see Fig. 5c), which is potentially due to the low abundance of *Aureispira* in the dataset.

**Figure S19.**
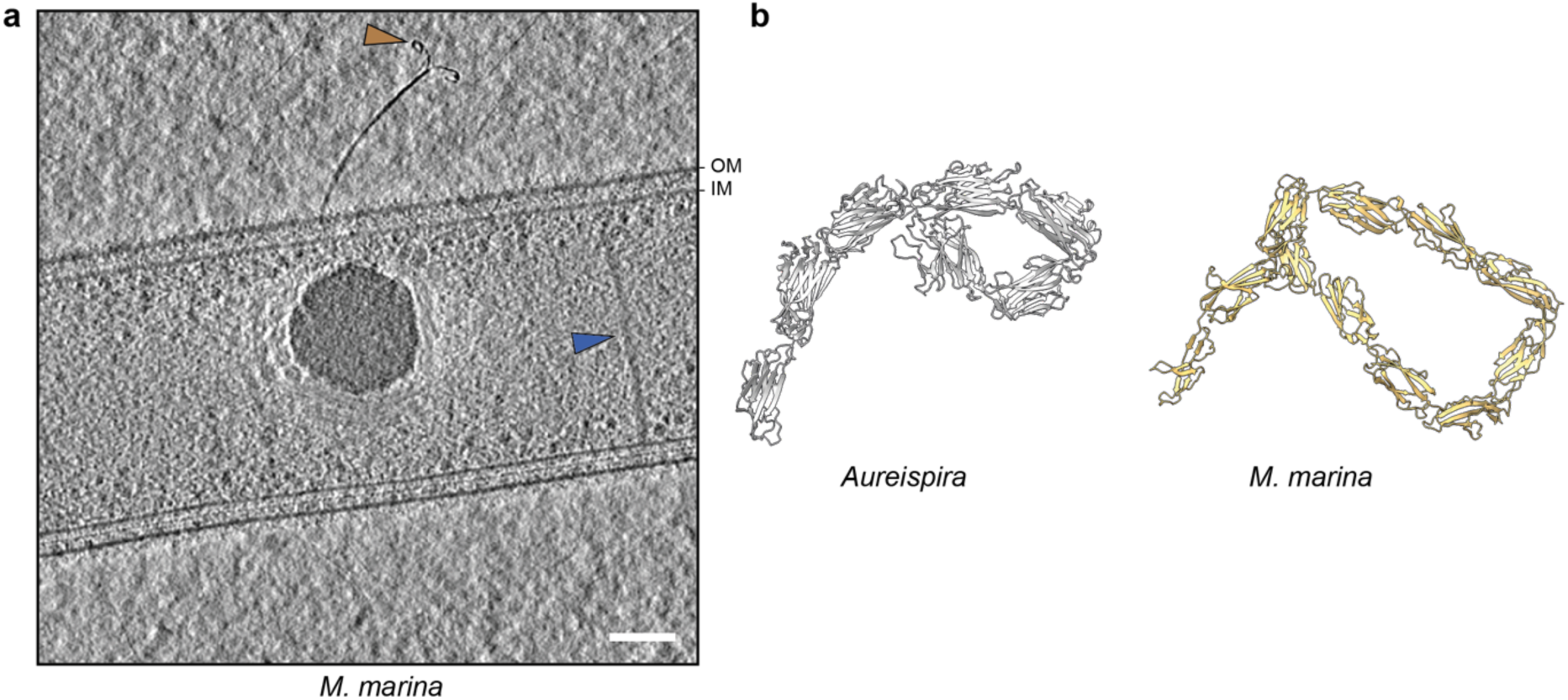
Characterization of grappling hooks in another *Bacteroidota* species. **a:** A slice through a cryo-tomogram of *M. marina* shows grappling hook-like structures on the bacterial surface. Brown arrowhead: grappling hook. Blue arrowhead: extended T6SS. Bar: 100 nm. Thickness of the slice: 19.8 nm. **b:** Alphafold2 predicted atomic model of the N-terminal region of *M. marina* GhpA homolog shows a larger hook-like architecture compared to the predicted model of the N-terminal region of *Aureispira* GhpA. The prediction of a larger hook fits the observed structure in the cryo-tomograms (see brown arrowhead in **a**). The first 1,000 amino acid residues of *M. marina* GhpA were used for the Alphafold2 prediction.

**Figure S20.**
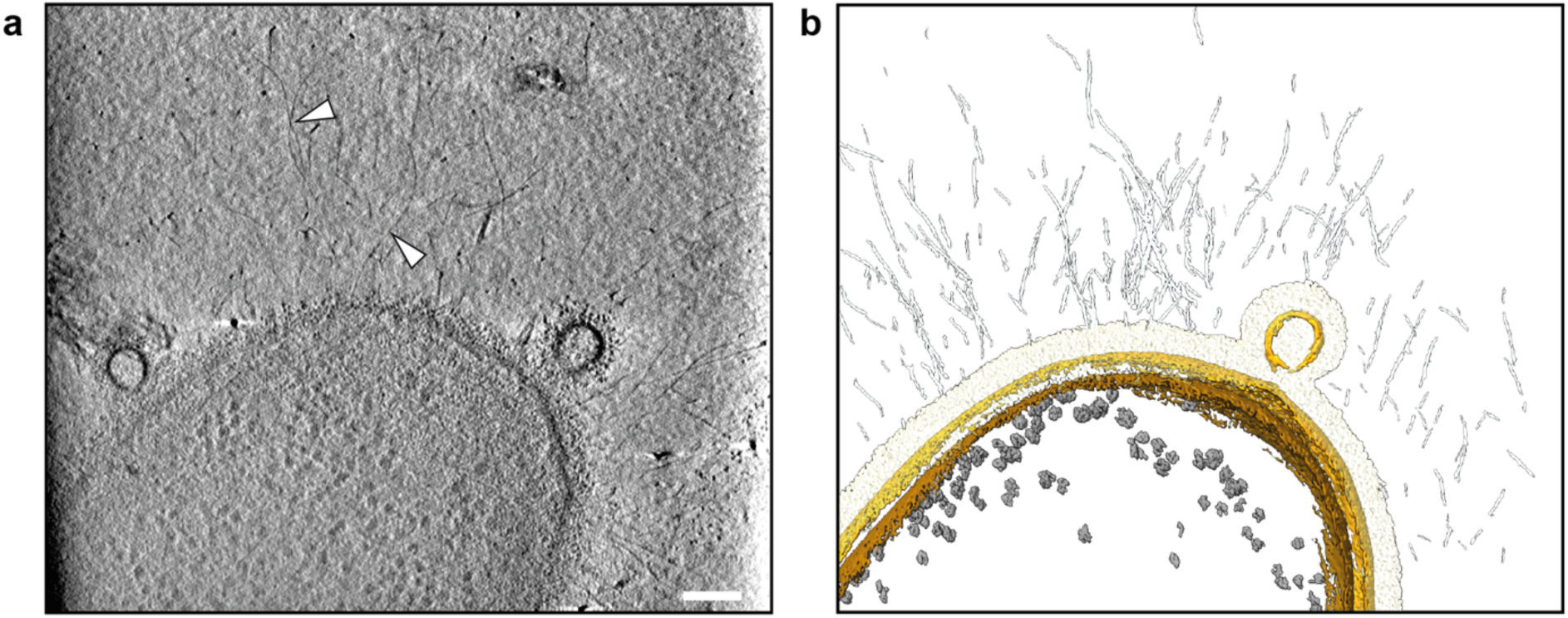
CryoET revealed putative SprB filaments on the surface of *Aureispira*. **a:** A slice through a cryo-tomogram of *Aureispira* reveals potential SprB filaments (arrowheads) on the bacterial surface. Bar: 100 nm. Thickness of the slice: 13.8 nm. **b:** A volume segmentation of the tomogram shown in **a** visualizes the 3D organization, including IM (brown), OM (yellow), extracellular surface density (light yellow), potential SprB filaments (light gray), and ribosomes (dark gray).

**Figure S21.**
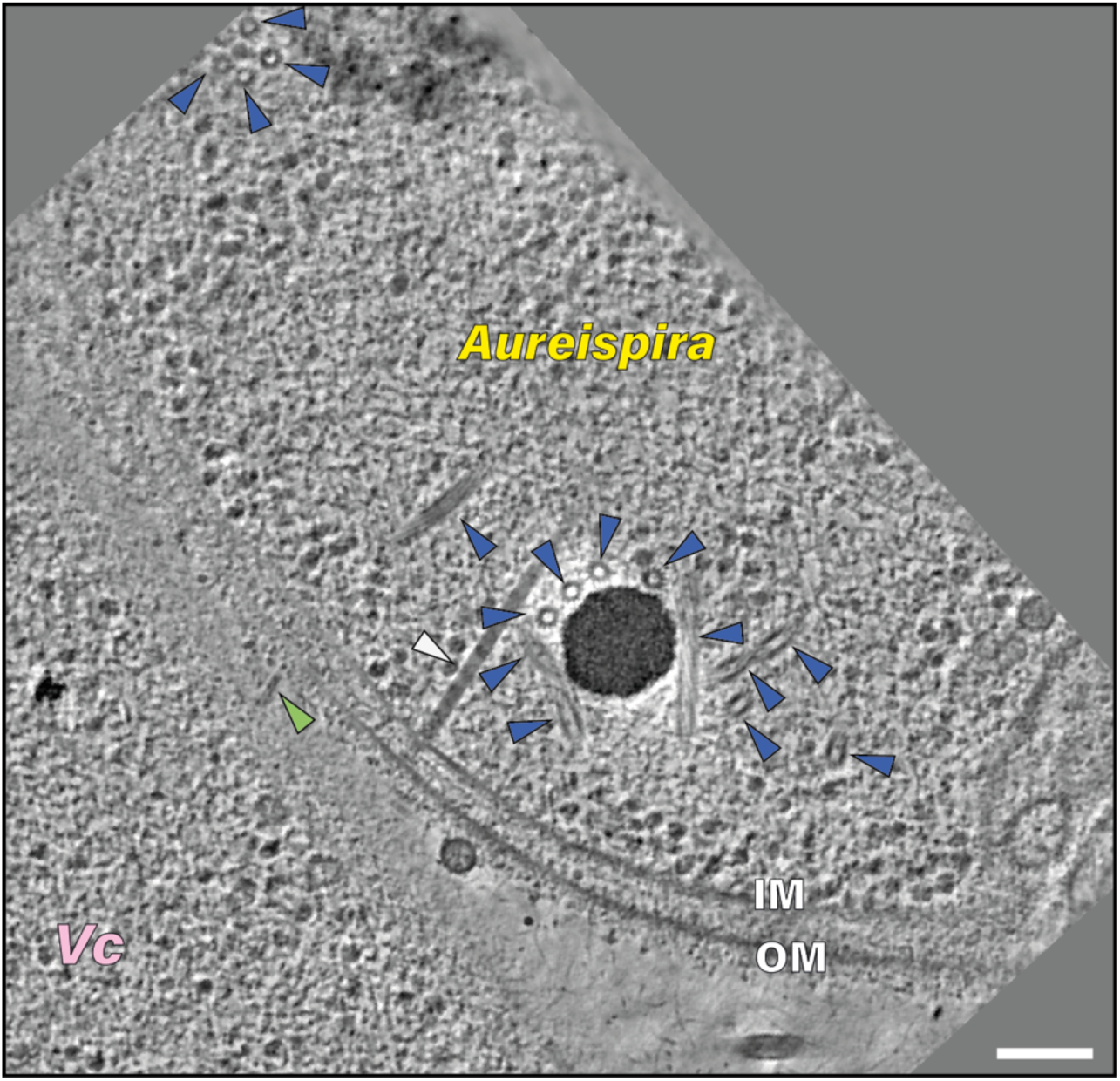
More than 16 contracted sheaths were found in an *Aureispira* cell co-incubated with *Vc*. A slice through a cryo-tomogram of an *Aureispira* cell co-incubated with *Vc*. In this example, 16 contracted T6SS sheath structures (blue arrowheads) were observed. White arrowhead: extended T6SS. Green arrowhead: T6SS inner tube penetrating the *Aureispira* cell envelope towards the *Vc* cell. OM: outer membrane. IM: inner membrane. Bar: 100 nm. Thickness of the slice: 13.8 nm.

## Supplementary movies

**Movie S1: Time-lapse LM movie of *Aureispira* gliding towards *Vc* cells, which subsequently lyse after cell-cell contact with *Aureispira*. The movie corresponds to Fig. 1a. Bar: 5 μm. The time point (in mins) of each frame is indicated in the upper left corner.**

**Movie S2: Time-lapse LM movie of *Aureispira* filaments gliding towards *Vc* cells. After cell-cell contact, *Vc* cells showed signs of rounding and lysed. The movie corresponds to Fig. S1b. Bar: 5 μm. The time point (in mins) of each frame is indicated in the upper left corner.**

**Movie S3: Cryo-tomogram of an *Aureispira* cell, followed by segmentation (corresponds to Fig. 1e/f), shows unique structures: grappling hooks (brown), extended T6SSs sheath(blue), T6SS baseplate (orange), T6SS trans-envelope complex (turquoise) and extracellular antennae (gray). Outer membrane (yellow), inner membrane (light brown). Bar: 100 nm.**

**Movie S4: Cryo-tomogram of FIB-milled *Aureispira-Vc* lawn, followed by segmentation (corresponds to Fig. 3l/m), shows a contracted T6SS sheath (blue) and an associated expelled inner tube (green). The inner tube is seen penetrating the OM of *Vc*, apparently resulting in membrane vesiculation. Outer membrane (yellow), inner membrane (light brown), and ribosomes (gray). Bar: 100 nm.**

**Movie S5: Time-lapse LM movie of T6SS-negative *Aureispira* gliding towards *Vc* cells. Even after extensive cell-cell contact, no prey cell lysis can be observed. Bar: 10 μm. The time point (in mins) of each frame is indicated in the upper left corner.**

## Supplementary material

**Supplementary Material S1: 35 pairs of *Saprospiraceae* and *Vibriocaceae* sequences showed predator-prey like interaction with high confidence from a marine time-resolved metagenomic dataset.**

**Table S1.** *Aureispira* CIS structural components compared with components of other CISs.

| *Aureispira* | *Nostoc* sp. PCC 7120 | *Algoriphagus machipongonensis* | *Pseudoalteromonas luteoviolacea* | *Candidatus Amoebophilus asiaticus* | Proteobacteria | Putative Function |
| --- | --- | --- | --- | --- | --- | --- |
| T6SS<sup>iv</sup> | tCIS | eCIS | eCIS | T6SS<sup>iv</sup> | T6SS<sup>i</sup> |  |
| Cis1a/b/c | Cis1 | Alg1 | JF50_12680 | Aasi_1077 | Hcp | Inner tube |
| Cis2 | Cis2 | Alg2 | JF50_12675 | Aasi_1074 | TssB/C | Outer sheath |
| Cis6 | Cis6 | Alg6 | JF50_12690 | Aasi_1078 | - | Spike plug |
| Cis7 | Cis7 | Alg7 | JF50_12695 | Aasi_1079 | - | Tube initiator |
| Cis8 | Cis8 | Alg8 | VgrG | Aasi_1080 | VgrG | Spike |
| Cis9 | Cis9 | Alg9 | JF50_12705 | Aasi_1082 | TssE | Baseplate sheath adaptor |
| Cis10 | Cis10 | Alg10 | - | Aasi_1081 | PAAR | Spike tip |
| Cis11 | Cis11 | Alg11 | JF50_12710 | Aasi_0557 | TssF/G | Baseplate wedge |
| Cis12 | Cis12 | Alg12 | JF50_12715 | Aasi_1083 | TssF | Baseplate wedge |
| Cis13 | Cis13 | - | JF50_12720 /12580 | Aasi_0556 | - | Tail fiber |
| Cis14 | Cis14 | Alg14 | JF50_12725 | Aasi_1806 | - | Tape measure |
| Cis15 | - | Alg15 | JF50_12570 | - | ClpV | ATPase |
| Cis16A/B | Cis16A/B | Alg16A/B | JF50_12575 | Aasi_1072 | TssA | Distal cap |

**Table S2.**
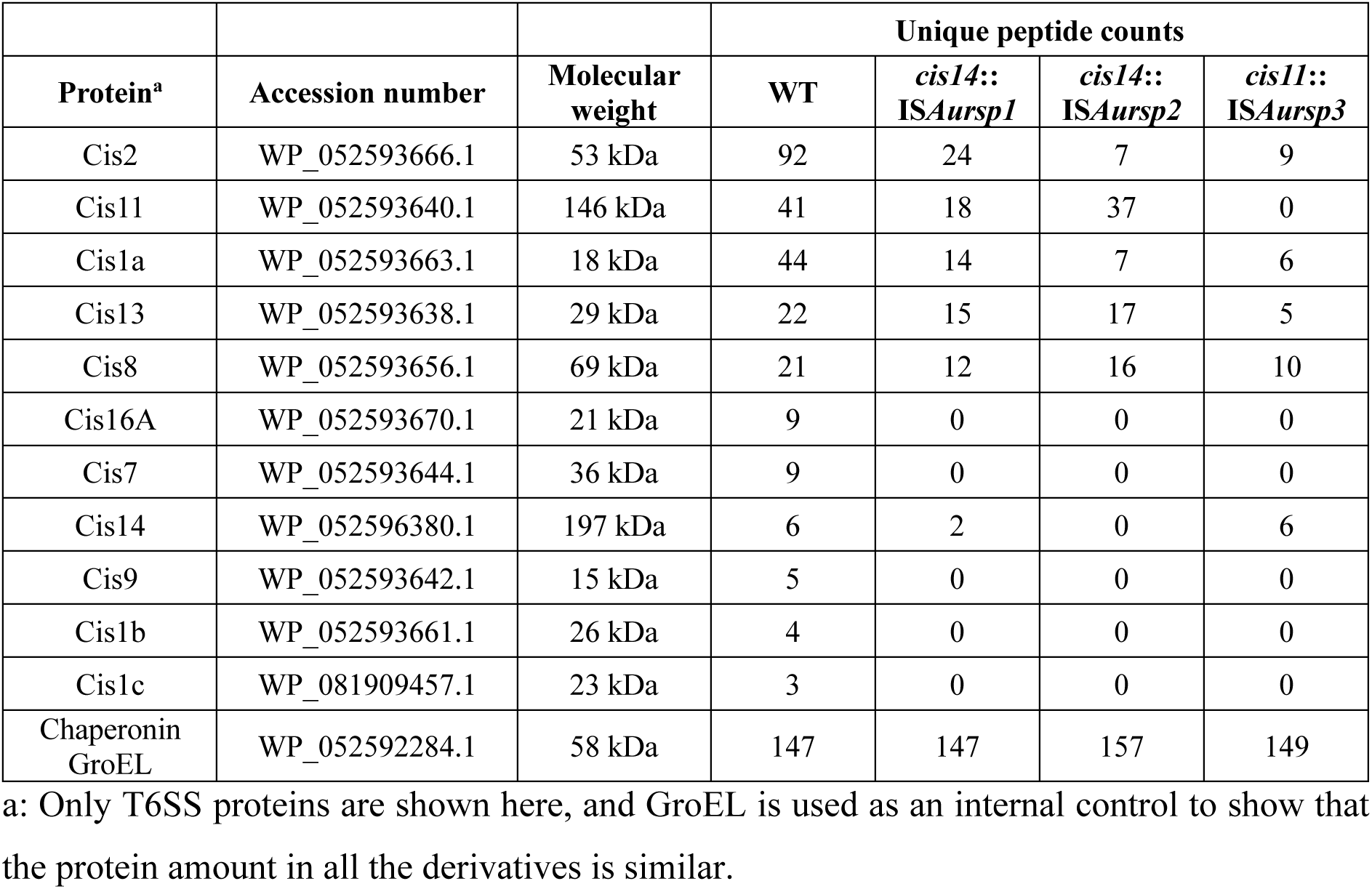
Mass spectrometry analysis of crude purification of *Aureispira* T6SSs.

**Table S3.** Copy numbers of the ISs in the genome of *Aureispira* derivatives.

| ISs<br>Derivatives | *ISAursp1* | *ISAursp2* | *ISAursp3* |
| --- | --- | --- | --- |
| WT | 2 | 3 | 2 |
| *cis14::ISAursp1* | 2<br>+1 (disruption of *cis14*) | 3 | 1 |
| *cis14::ISAursp2* | 2 | 3<br>+1 (disruption of *cis14*) | 1 |
| *cis11::ISAursp3* | 2 | 3 | 1<br>+1 (disruption of *cis11*) |

**Table S4.**
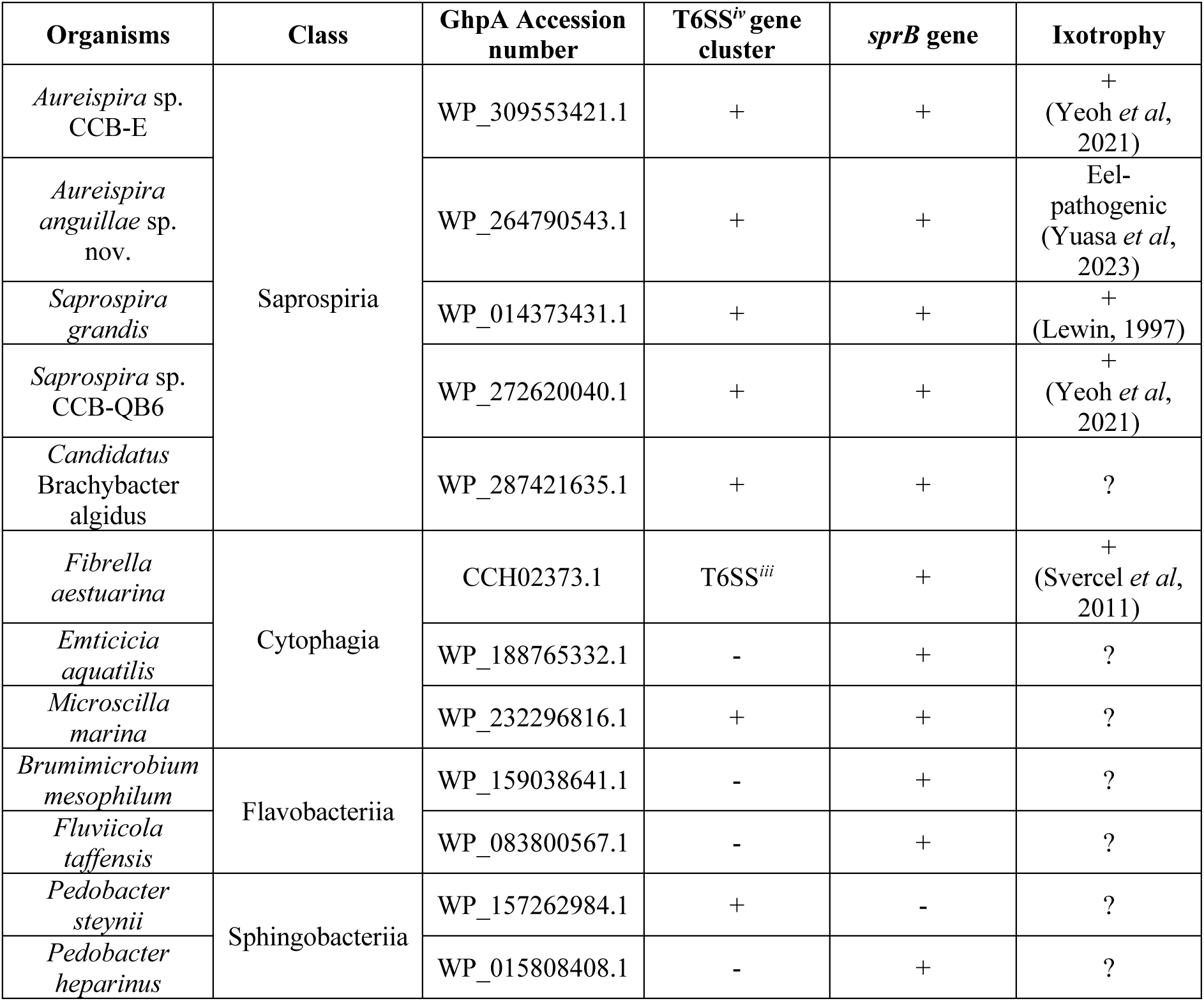
Bacteria with homologs of *ghpA* often also harbor genes for T6SS*^iv^* and *sprB*.

| Organisms | Class | GhpA Accession number | T6SS<sup>iv</sup> gene cluster | *sprB* gene | Ixotrophy |
| --- | --- | --- | --- | --- | --- |
| *Aureispira* sp. CCB-E | Saprospiria | WP_309553421.1 | + | + | +(Yeoh *et al*, 2021) |
| *Aureispira anguillae* sp. nov. |  | WP_264790543.1 | + | + | Eel-pathogenic (Yuasa *et al*, 2023) |
| *Saprospira grandis* |  | WP_014373431.1 | + | + | +(Lewin, 1997) |
| *Saprospira* sp. CCB-QB6 |  | WP_272620040.1 | + | + | +(Yeoh *et al*, 2021) |
| *Candidatus Brachybacter algidus* |  | WP_287421635.1 | + | + | ? |
| *Fibrella aestuarina* | Cytophagia | CCH02373.1 | T6SS<sup>iii</sup> | + | +(Svercel *et al*, 2011) |
| *Emticicia aquatilis* |  | WP_188765332.1 | - | + | ? |
| *Microscilla marina* |  | WP_232296816.1 | + | + | ? |
| *Brumimicrobium mesophilum* | Flavobacteriia | WP_159038641.1 | - | + | ? |
| *Fluviicola taffensis* |  | WP_083800567.1 | - | + | ? |
| *Pedobacter steynii* | Sphingobacteriia | WP_157262984.1 | + | - | ? |
| *Pedobacter heparinus* |  | WP_015808408.1 | - | + | ? |

**Table S5.** T9SS main components in *Aureispira* genome.

| Protein<sup>a</sup> | Accession number<br>in *Aureispira* | Function |
| --- | --- | --- |
| OM translocon complex |  |  |
| Sov/SprA | WP_052598092.1 | OM translocon (Lauber *et al*, 2018) |
| Plug | WP_052591900.1 | Block the translocon when no substrate binding (Lauber *et al*, 2018) |
| PorK/GldK | WP_052594856.1 | OM protein forming a ring structure with PorN/GldN (Gorasia *et al*, 2016; Song *et al*, 2022) |
| PorN/GldN | WP_052594849.1 | Periplasmic protein forming a ring structure with PorK/GldK (Gorasia *et al*, 2016; Song *et al*, 2022) |
| PorW/SprE | WP_052598013.1 | Possibly connecting the ring to the translocon (Gorasia *et al*, 2022a) |
| IM/periplasmic rotary motor |  |  |
| PorL/GldL | WP_052594853.1 | IM rotor (Hennell James *et al*, 2021, 2022) |
| PorM/GldM | WP_052594851.1,<br>WP_052594847.1,<br>WP_052594843.1,<br>WP_052594812.1 | Rotor spanning from IM to periplasm (Leone *et al*, 2018; Hennell James *et al*, 2021, 2022) |

**a:** The names shown here are from the two extensively studied T9SS model organisms, *Flavobacterium johnsoniae* and *Porphyromonas gingivalis*.

**Table S6.** Known ixotrophic predator, their prey and their isolation sites.

| Predator | Prey | Place of isolation | Reference(s) |
| --- | --- | --- | --- |
| *Saprospira albida* | Cyanobacteria<br>*Microcystis aeruginosa* | A hypertrophic lake in South Africa | (Ashton & Robarts, 1987) |
| *Saprospira grandis* | *Vibrionaceae* bacteria | Subtidal sand, polyethylene sheeting and dead didemnid ascidian colonies in the Bahama Island | (Lewin, 1997) |
| *Saprospira* sp.<br>SS98.5 | Diatom<br>*Chaetoceros ceratosporum* | Kagoshima Bay in Japan | (Furusawa *et al*, 2005) |
| *Saprospira* sp.<br>PdY3 | Cyanobacteria<br>sprcies | Dianchi Lake (freshwater lake) in Kunming City of Yunnan Province in China | (Shi *et al*, 2006) |
| *Fibrella aestuarina* | Cyanobacteria<br>*Nostoc muscorum* | Mud from tidal flats in Fedderwardersiel from the Northe Sea coast of Germany | (Svercel *et al*, 2011) |
| *Aureispira* sp. and<br>*Saprospira* sp. | *Vibrio* sp. | Seaweed (genus *Ulva*) from Queens Bay of Penang in Malaysia | (Furusawa *et al*, 2015b, 2015a; Yeoh *et al*, 2021) |

**Table S7.**
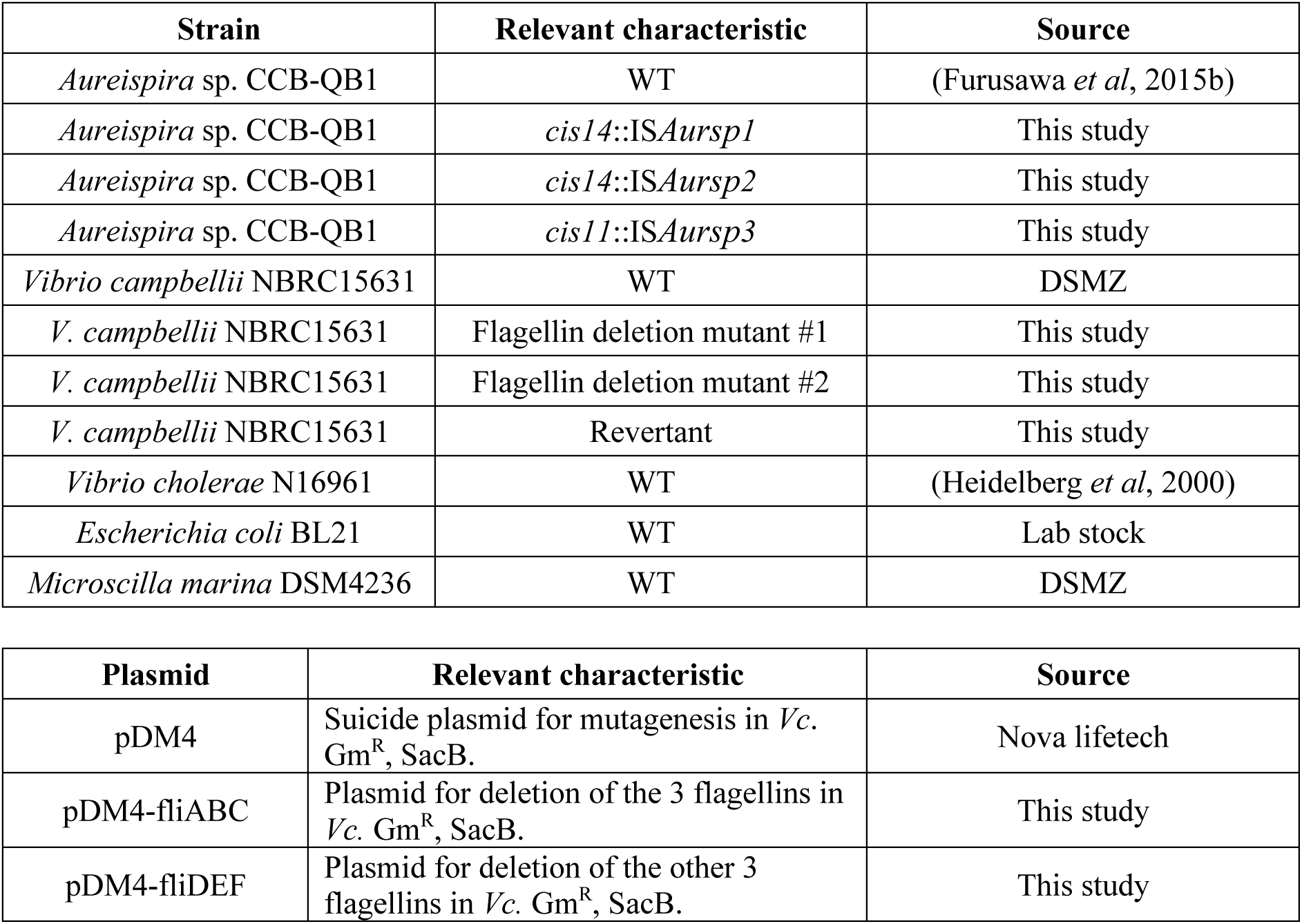
Bacterial strains and plasmids used in this study.

| Strain | Relevant characteristic | Source |
| --- | --- | --- |
| *Aureispira* sp. CCB-QB1 | WT | (Furusawa *et al*, 2015b) |
| *Aureispira* sp. CCB-QB1 | *cis14::ISAursp1* | This study |
| *Aureispira* sp. CCB-QB1 | *cis14::ISAursp2* | This study |
| *Aureispira* sp. CCB-QB1 | *cis11::ISAursp3* | This study |
| *Vibrio campbellii* NBRC15631 | WT | DSMZ |
| *V. campbellii* NBRC15631 | Flagellin deletion mutant #1 | This study |
| *V. campbellii* NBRC15631 | Flagellin deletion mutant #2 | This study |
| *V. campbellii* NBRC15631 | Revertant | This study |
| *Vibrio cholerae* N16961 | WT | (Heidelberg *et al*, 2000) |
| *Escherichia coli* BL21 | WT | Lab stock |
| *Microscilla marina* DSM4236 | WT | DSMZ |

| Plasmid | Relevant characteristic | Source |
| --- | --- | --- |
| pDM4 | Suicide plasmid for mutagenesis in *Vc.* Gm<sup>R</sup>, SacB. | Nova lifetech |
| pDM4-fliABC | Plasmid for deletion of the 3 flagellins in *Vc.* Gm<sup>R</sup>, SacB. | This study |
| pDM4-fliDEF | Plasmid for deletion of the other 3 flagellins in *Vc.* Gm<sup>R</sup>, SacB. | This study |

**Table S8.**
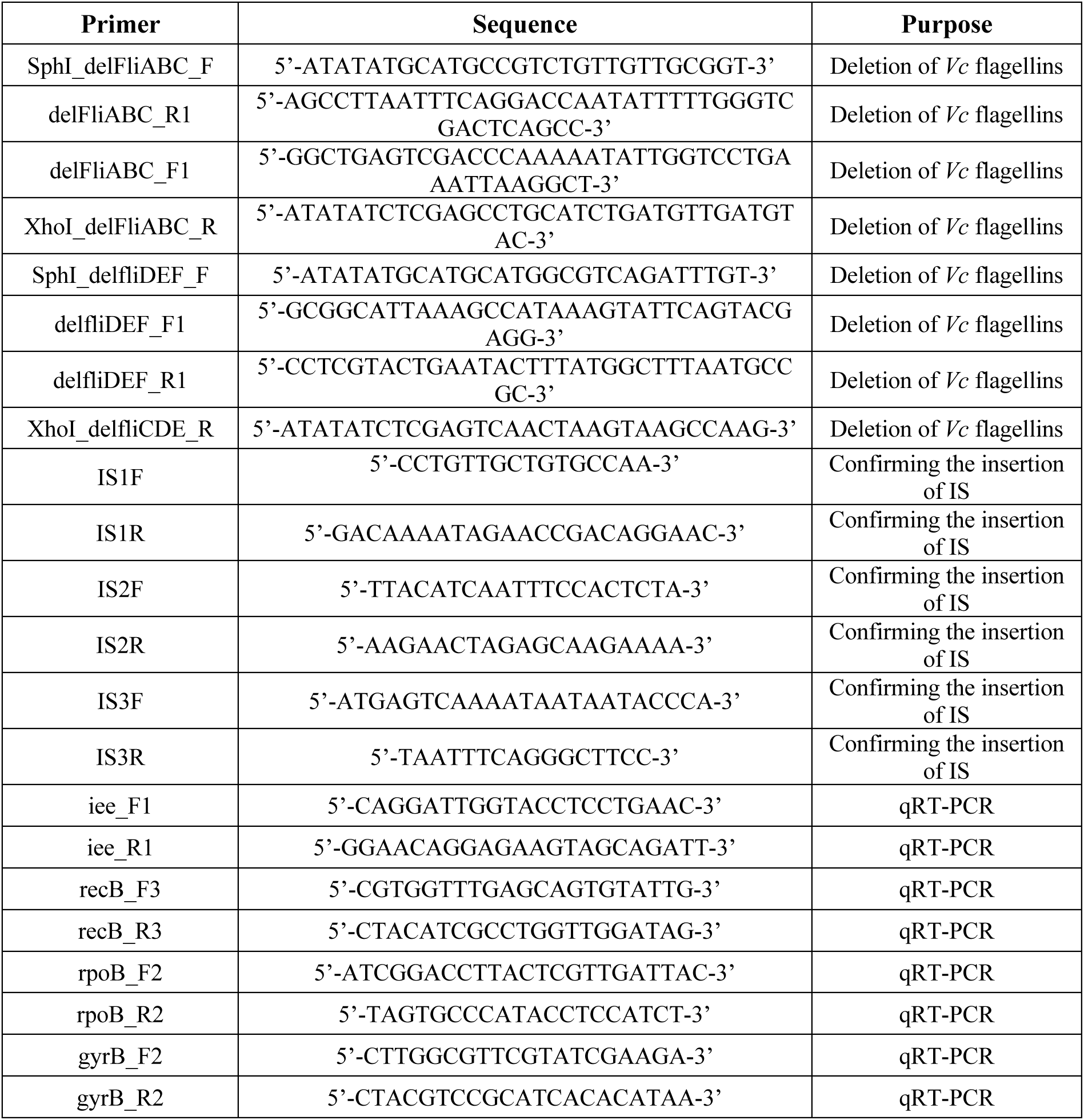
Primers used in this study.

**Table S9.**
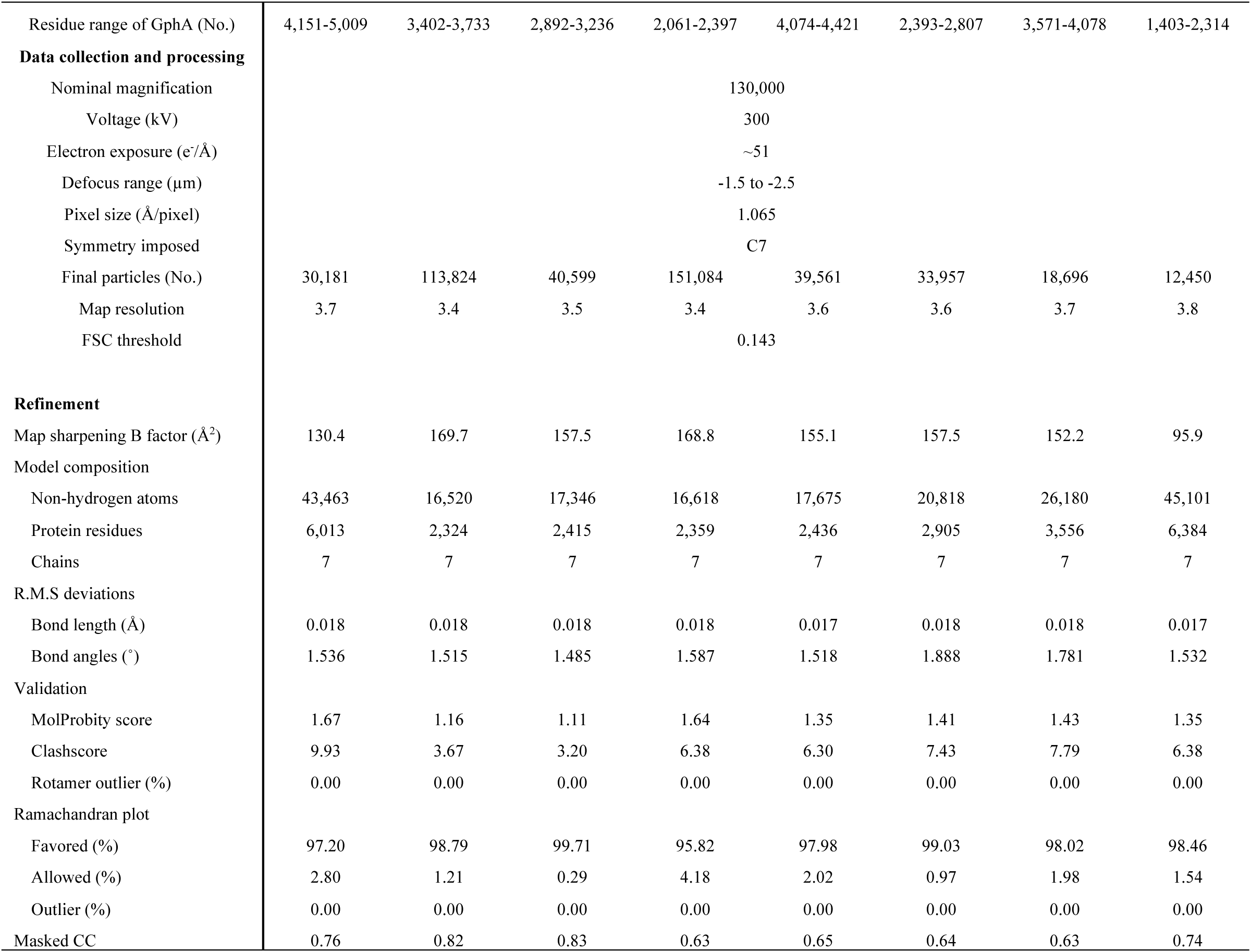
Single particle cryoEM data statistical analysis.

